# Engineering dynamic gates in binding pocket of penicillin G acylase to selectively degrade bacterial signaling molecules

**DOI:** 10.1101/2023.05.09.538545

**Authors:** Michal Grulich, Bartlomiej Surpeta, Andrea Palyzová, Helena Marešová, Jiří Zahradník, Jan Brezovsky

**Author notes:** Corresponding Authors, (M.G); (J.B.). These authors contributed equally (Michal Grulich, Bartlomiej Surpeta).

## Abstract

The rapid rise of antibiotic-resistant bacteria necessitates the search for alternative, unconventional solutions, such as targeting bacterial communication. Signal disruption can be achieved by enzymatic degradation of signaling compounds, reducing the expression of genes responsible for virulence, biofilm formation, and drug resistance while evading common resistance mechanisms. Therefore, enzymes with such activity have considerable potential as antimicrobial agents for medicine, industry, and other areas of life. Here, we designed molecular gates that control the binding site of penicillin G acylase to shift its preference from native substrate to signaling molecules. Using an ensemble-based design, three variants carrying triple-point mutations were proposed and experimentally characterized. Integrated inference from biochemical and computational analyses demonstrated that these three variants had markedly reduced activity towards penicillin and each preferred specific signal molecules of different pathogenic bacteria, exhibiting up to three orders of magnitude shifts in substrate specificity. Curiously, while we could consistently expand the pockets in these mutants, the reactive binding of larger substrates was limited, either by overpromoting or overstabilizing the pocket dynamics. Overall, we demonstrated the designability of this acylase for signal disruption and provided insights into the role of appropriately modulated pocket dynamics for such a function. The improved mutants, the knowledge gained, and the computational workflow developed to prioritize large datasets of promising variants may provide a suitable toolbox for future exploration and design of enzymes tailored to disrupt specific signaling pathways as viable antimicrobial agents.

## INTRODUCTION

Antibiotic-resistant bacteria have become an issue of global priority and will require effective approaches and solutions in the coming years.^1^ Increasing bacterial resistance derives from several sources, including multiple bacterial mechanisms that help them to cope with the antibiotics or drug-like molecules,^2^ promoted by the direct action of antibiotics on microorganisms, which exerts strong pressure to evolve multiple resistance approaches to survive.^3^ It is further promoted by the still widespread misuse of antibiotics in various areas of life to maximize profits or unjustified prophylactic prescription of antibiotics.^4,5^ Finally, it was shown that resistance genes could be transferred between the same or even different bacterial species without exposure to the antimicrobial agent.^6–9^ Therefore, collaborative efforts on a global and local scale are required to cope with the above-mentioned challenges. While misuse can be gradually reduced by education, appropriate control, and regulations, substituting antibiotics in various fields or overcoming already widespread resistance mechanisms and inefficiency of known antibiotics constitute non-trivial tasks. It is further complicated by the void in discovering new generations of antibiotics, with the last inventions arising nearly 30 years ago.^10,11^ Therefore, the constant search for new alternatives is crucial. Such approaches should preferably be able to escape the existing resistance mechanisms and limit the potential development of resistance in the future.

Interestingly, one recently studied approach targets a bacterial communication process called quorum sensing (QS). In gram-negative bacteria, this process is primarily induced by N-acyl-homoserine lactones (AHLs) used as signaling molecules for synthase/transcriptional regulator systems. In these systems, the synthase enzyme produces the signaling molecule, while the receptor recognizes the signal and affects the expression of QS-dependent genes, including the synthase.^12^ This process enables the collective response of bacteria and controls the expression of genes responsible for virulence, swarming, spreading resistance, or biofilm formation in a cell-density-dependent manner. This communication can be disrupted by the quorum quenching (QQ) process, achieved either by inhibiting the specific synthase involved in the signaling or its receptor, or by enzymatically inactivating the signaling molecule. While inhibiting intracellular proteins may suffer from similar resistance mechanisms as for current antibiotics, signal inactivation constitutes a more promising approach due to its extracellular targeting, markedly reducing the selective pressure.^13^ Amongst three enzyme classes that process AHLs, amidases/acylases are of significant interest due to their irreversible hydrolytic action.^14^ Notably, acylases capable of hydrolyzing AHLs have shown promise in many applications in different fields, including medicine, industry, agriculture, aquaculture, and wastewater treatment among others.^14–19^ Given the above facts, the increased application potential of QQ enzymes is evident in various areas. However, it is worth noting the obstacles that limit their applicability. When considering broad-scale utilization, aspects such as established procedures for mass production, immobilization, high tolerance to industrial processes, and shelf-life are critical. Although numerous enzymes have been shown to exhibit non-negligible QQ activity, not all are suitable for wide-scale adoption. For example, *Pseudomonas aeruginosa* acyl-homoserine lactone acylase (paPvdQ), despite being the most investigated AHL acylase with verified QQ activity, ^19–21^ lacks an optimized procedure for industrial-scale production. Furthermore, PF2571 AHL acylase was shown to possess high activity towards long AHLs, and its variant has been modulated to cleave the shorter substrates efficiently. However, while these enzymes show high potential for preventing food spoilage, the lack of optimized and cost-effective production is currently considered a major application obstacle.^18,22^

Instead of devising processes for improving the applicability of individual AHL-degrading enzymes, we propose to focus on engineering this activity for already well-established *Escherichia coli* penicillin G acylase (ecPGA), best known for its role in the synthesis of semi-synthetic β-lactam antibiotics.^23–26^ ecPGA represents a perfect template for engineering due to the availability of multiple high-quality crystal structures, broad knowledge of its properties, and multiple protocols for expression and purification, as documented by successful rational engineering cases.^27,28^ Previously, we have shown that ecPGA possesses degrading activity towards short- to medium-long AHLs.^29^ Moreover, by contrasting its mechanism of action with paPvdQ, we have revealed dynamic determinants that govern the AHL-degrading activity and constitute promising targets for engineering the potency of ecPGA towards QQ applications. These include (i) residues forming the gate preventing opening of the bottom part of ecPGA’s acyl-binding cavity that compromise productive binding of longer substrates, and (ii) the dynamics of molecular gates themselves that control the access to the acyl-binding site.^29^ Since bacterial species, including the beneficial microbiota, often utilize distinct AHLs,^13^ we believe that relevant QQ solutions for various applications could be devised by engineering these molecular determinants.

In this work, we aimed at obtaining ecPGA variants with specificity shifted from native penicillin G (PenG) to varied bacterial signaling molecules, AHLs. Using computationally-driven workflow targeting the structure and dynamics of molecular gates controlling access and depth of acyl-binding pocket of this enzyme, we designed several ecPGA variants with experimentally validated shift in specificity and preference for particular AHLs used by different pathogenic bacteria. This work confirms the feasibility of customizing ecPGA towards tailored AHLs-degrading specificity. By in-depth analysis of the mutants, we provide molecular details underlying the improved activity and altered specificity. Furthermore, we point out additional limitations in our strategy that can guide subsequent engineering efforts.

## RESULTS

### Identifying hot spots in acyl-binding sites of ecPGA to tailor its QQ activities

A recent investigation found insufficient accommodation of moderate-length C08-HSL (N-octanoyl-L-homoserine lactone) in the wild-type ecPGA acyl-binding pocket.^29^ This, together with arrangements in the paPvdQ enzyme that are selective towards longer AHLs (Figure 1A,B),^30^ inspired the exploration of residues whose modification could expand the ecPGA pocket, while improving the stabilization of the bound substrates by altering the dynamics of two molecular gates at the entrance and at the bottom of the pocket. Using molecular dynamics (MD) simulations that were generated in our previous work,^29^ we identified residues that frequently formed the pocket of wild-type ecPGA (Figure 1C,D, Figure S1) and excluded catalytic residues Ser1β, Gln23β, Ala69β, Asn241β, and gating residues Arg145α, Phe146α and Phe24β (Figure S1). This procedure left us with two hot-spot resides, namely Met142α gating residue preventing opening of the deeper parts of the pocket, to expand the pocket (Figure 1C), and Ile177β to stabilize the molecular gates Phe146α and Phe24β at the entrance (Figure 1D). Additionally, we considered the third position with the capacity to further elongate the engineered pocket. In this case, we used the binding pose of the transition state analog present in the paPvdQ structure (PDB ID: 4M1J) to estimate the direction of pocket expansion (Figure 1A-C). From the residues forming the target region, Pro49β, Phe57β, Phe138α, and Trp154β, we chose the most mutable residue identified by HotSpot Wizard 3.0 tool,^31,32^ namely Phe138α, as the third hot-spot (Figure S1).

**Figure 1.**
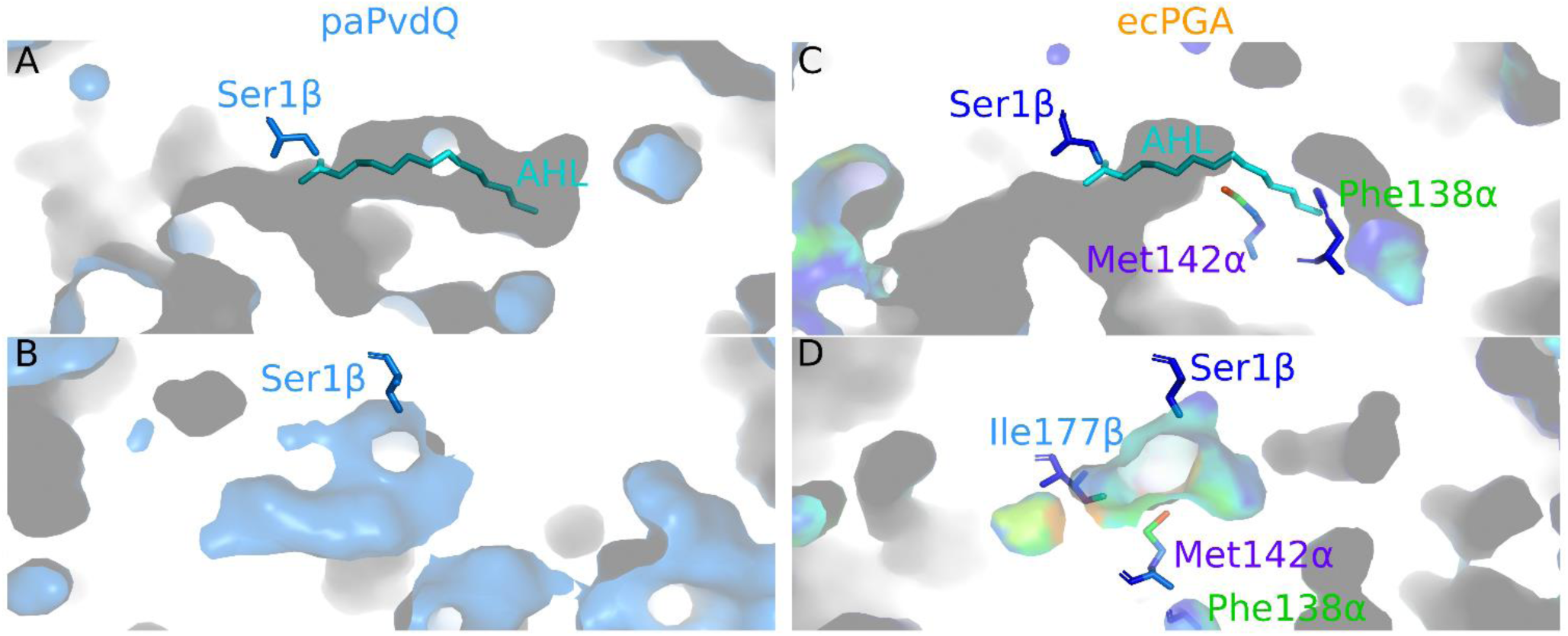
Computational design of triple-point ecPGA mutants. **A**) Side view of the paPvdQ binding pocket profile with transition state analog shown as cyan sticks. **B**) Entrance to the paPvdQ binding pocket. **C**) Side view of the wild-type ecPGA binding pocket with paPvdQ transition stage analog shown as cyan sticks superimposed, residues selected for mutagenesis – Phe138α and Met142α – are shown as sticks. **D**) Entrance to the ecPGA binding pocket, residue Ile177β selected for mutagenesis to stabilize molecular gates at the entrance is shown as sticks. The coloring of ecPGA was based on the conservation of contribution to the binding pocket formation from mdpocket^33^ analyses (C-D).

### Computational saturation mutagenesis and filtering identified six variants with enhanced pocket ensembles

Mutagenesis was performed in a two-step workflow using the FoldX^34^ tool (Figure 2A,B). First, we considered mutations of residues Met142α and Ile177β. For Met142α, we allowed 19 substitutions to all standard amino acids. In the case of Ile177β, we limited the mutagenesis to residues bulkier than the native isoleucine without introducing additional hydrogen bonding capacity, reducing the possible substitutions at this position to phenylalanine only. By introducing the mutations in both possible orders, we generated 60 structures for each mutant. Using TransportTools^35^ comparative module, we analyzed the pocket conformation in the structural ensembles of mutants and identified a consistently present supercluster that best represented the acyl-binding pocket. By considering the geometric properties of engineered pockets and the predicted effects of mutations on ecPGA stability, we pre-selected only the most promising candidates for the second design step. The criteria for selection, irrespective of the order of introduced mutations, were as follows: (i) a large-margin destabilization penalty (defined here as average energy change extended by the corresponding standard deviation) per mutation that cannot exceed 2 kcal/mol, (ii) conformational behavior of the engineered pocket from the comparative analysis was improved over the wild-type pocket considering the number of frames with an open pocket, average length, average and maximal bottleneck radii, and average throughput (Table S1). Such a procedure resulted in prioritizing two mutations for the next step – Met142αAla & Ile177βPhe and Met142αSer & Ile177βPhe, which were analyzed together with their reverse combinations (Table S2).

**Figure 2.**
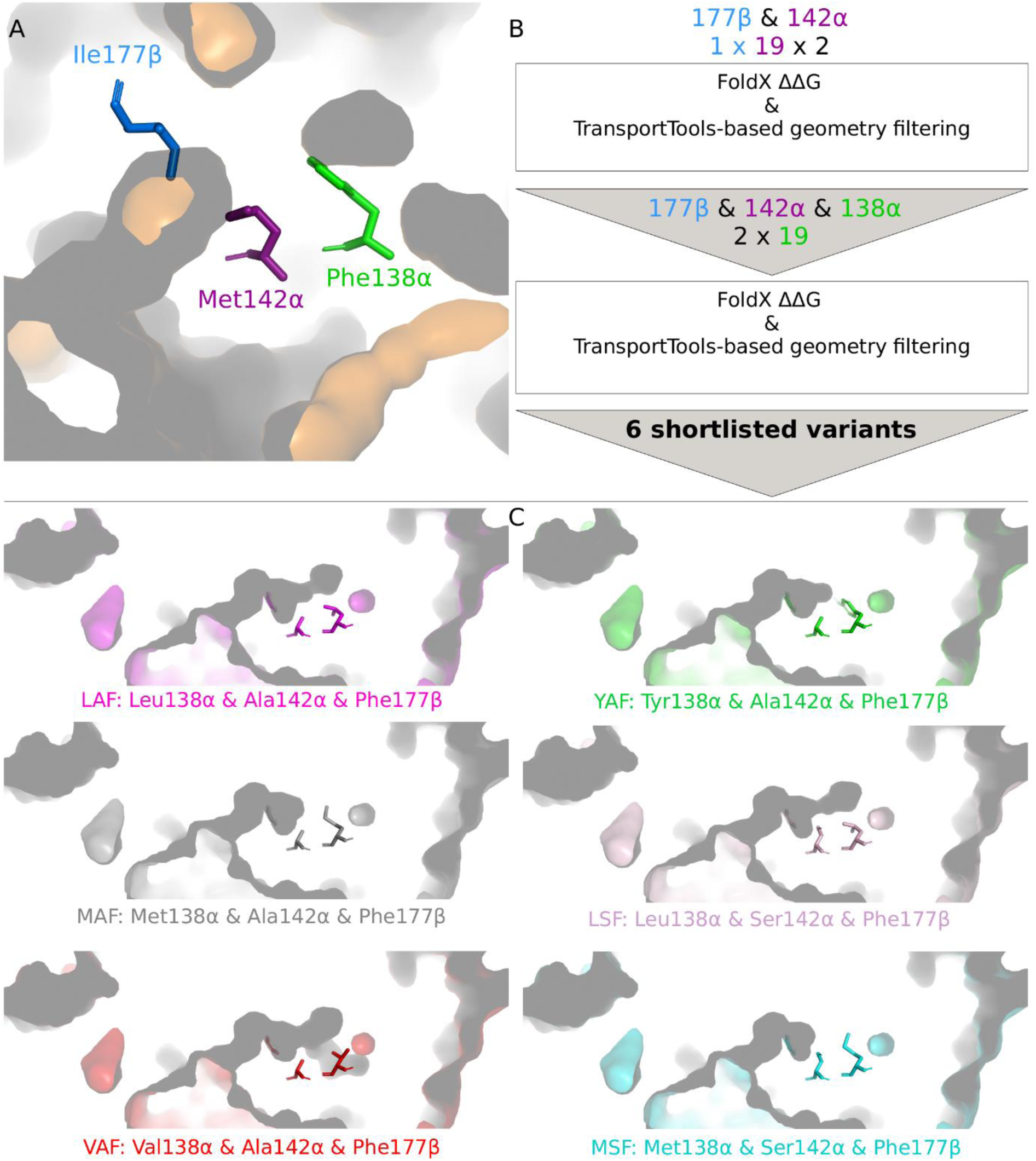
Schematic representation of the *in-silico* mutagenesis procedure. **A**) wild-type ecPGA pocket with residues selected for mutagenesis shown as blue, purple, and green sticks. **B**) Step-wise procedure of possible variants generation and filtering based on their predicted stability and geometries of engineered pockets. **C**) Six best candidates selected for subsequent detailed molecular investigation.

Then, we introduced 19 substitutions instead of the third hot-spot residue, Phe138α, into all generated structures of two double-point mutants and followed an analogous procedure for their evaluation, i.e., stability was estimated with FoldX and pocket ensemble geometry was investigated with TransportTools. The second filtering round consisted of the same maximal destabilization penalty of 2 kcal/mol per mutation, and stricter filters based on TransportTools geometries, requiring significantly improved average length, average and maximal bottleneck radii, and near permanent occurrence (Table S3). Reverse order mutations were already considered when calculating properties from the ensembles, which simplified the filtering process. Such filtering yielded seven three-point variants, including two very similar variants, VAF and VSF, with much higher overall destabilization penalties at over 5 kcal/mol (Table S4). While both displayed geometrical improvements, we decided to retain only the VAF variant with the deepest pocket to avoid the costly processing of two potentially unstable variants and produce a set of the six best candidates for further examination (Table S4 and Figure 2C).

Computational evaluation of substrate binding in selected variants predicted three variants with specificity tailored towards AHLs used by different pathogenic bacteria. The six variants selected were used as receptors for molecular docking experiments with a set of signaling molecules of different sizes, being C06-HSL (N-hexanoyl-L-homoserine lactone), C06-3O-HSL (N-3-oxo-hexanoyl-L-homoserine lactone), C08-HSL, C08-3O-HSL (N-3-oxo-octanoyl-L-homoserine lactone), C10-HSL (N-decanoyl-L-homoserine lactone), and C12-3O-HSL (N-3-oxo-dodecanoyl-L-homoserine lactone). Following strict filtering based on adherence to the reaction mechanism and favorable binding score, we selected appropriate and pre-organized complexes for the follow-up MD simulations (Table S5).

These simulations revealed improved binding and stabilization for various AHLs for the designed mutants compared to the wild-type enzyme. Starting from the complexes that exhibited favorable substrate stabilization for the first reaction step, we performed repetitive, 5 ns long MD simulations for each system. Here, we focused on the ability of the variant to maintain reactive stabilization for the nucleophilic attack reaction (Figure 3). Building on top of the known mechanism-based criteria to estimate reactivity,^27,36,37^ we predicted potential efficiency of variants by counting the fraction of the simulations that sampled proper AHL stabilization (Supplementary File 2). From here on, this metric is referred to as the reactive stabilization score (RSS), whose significance was verified by leave-one MD simulation-out approach (Table S6). By contrasting this score with the stabilization observed for wild-type ecPGA for individual AHLs, we estimated relative activity improvements for each investigated variant (Figure 4). Overall, our analysis indicated that the effects of the mutations that were introduced when contrasted to the wild-type enzyme (i) was primarily neutral or only slightly favorable for all ecPGA variants with shorter substrates (C06-, C06-3O-HSL), (ii) most of the designed mutants presented improved activities for medium-size substrates (C08-, C08-3O-HSL), and (iii) few candidates presented markedly improved activities for the longest C12-3O-HSL, which is also the most problematic one for the wild-type ecPGA with a shallow pocket.^29^

**Figure 3.**
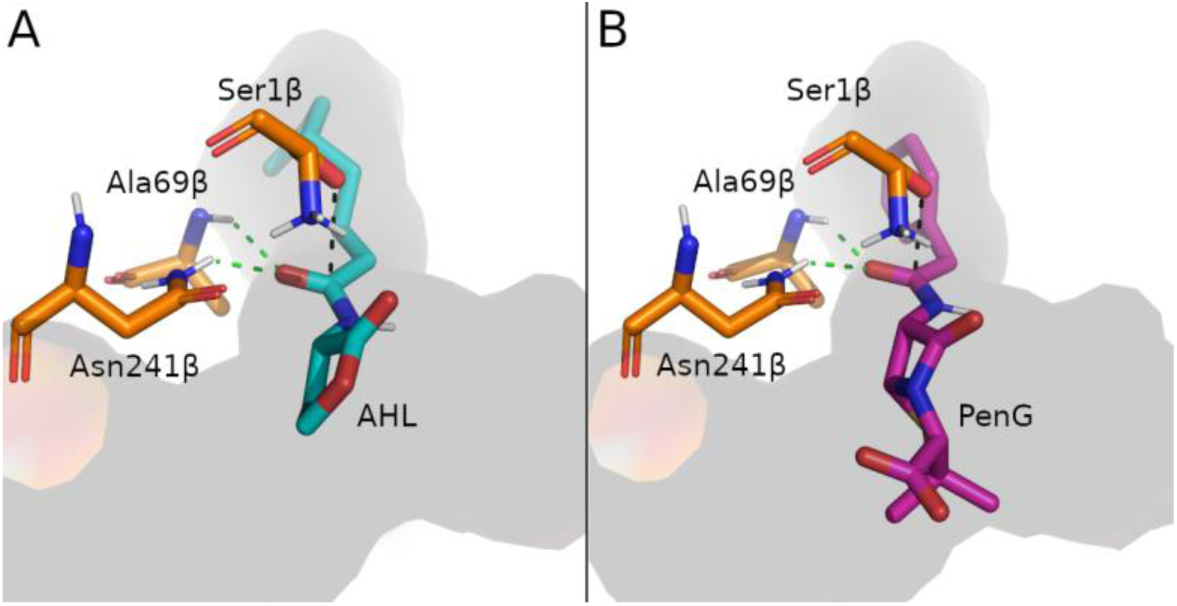
Geometric criteria based on the nucleophilic attack reaction mechanism. Those included substrates interactions with the oxyanion hole-stabilizing residues Ala69β and Asn241β (≤ 3.0 Å), nucleophile attack distance (≤ 3.3 Å) and angle (between 75-105°). **A**) Stabilization of AHL (cyan) and **B**) analogous stabilization of PenG (magenta).

**Figure 4.**
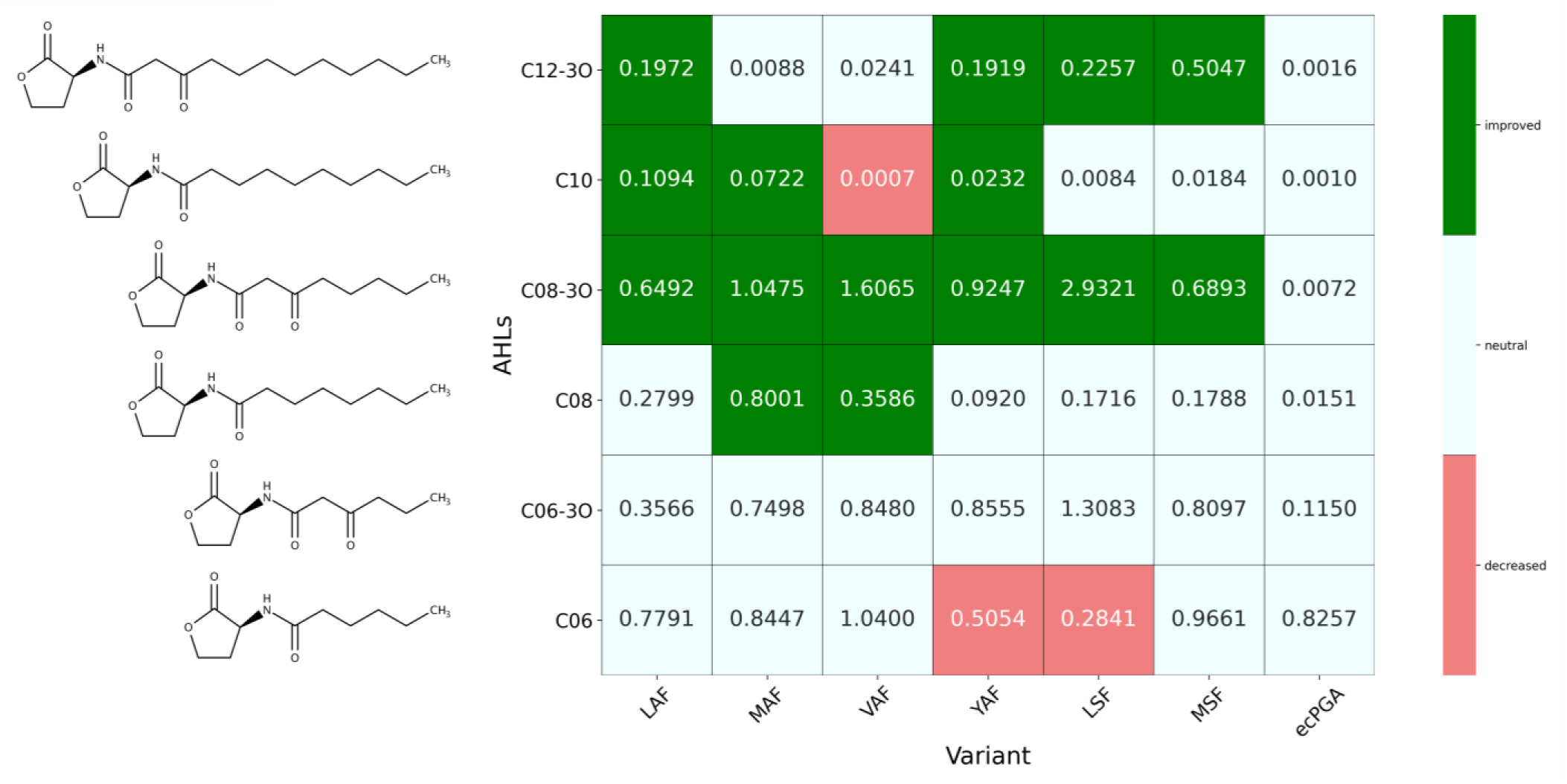
Predicted activities based on reactive stabilization score of six pre-selected ecPGA variants with a set of AHLs. Rows correspond to the studied AHLs, and columns to particular ecPGA variants, including the wild-type ecPGA. The coloring scheme was applied according to the reactive stabilization score relative to the wild-type ecPGA (relative RSS ranges: decreased: 0 – 0.75, neutral: 0.75 – 20, improved: >20).

We decided to take three of the most interesting candidates for experimental evaluation. The VAF mutant was chosen due to its unique, narrow specificity towards medium-size AHLs, including C08-HSL, a signaling molecule of pathogenic *Burkholderia cenocepacia*.^19^ The YAF mutant was selected for its overall improvement for longer substrates (C08-3O-HSL – C12-3O-HSL), in contrast to other mutants, especially the LSF mutant with seemingly similar properties but lacking the improvement for C10-HSL, the signaling compound of pathogenic *Vibrio* species.^38^ Finally, the MSF mutant was shortlisted, due to its preference for C12-3O-HSL, the signaling molecule of *P. aeruginosa*, and the preferred substrate of prototypical paPvdQ.^19,30^

### Experimental characterization confirmed activity profiles of the three best mutants agreed with the predictions

The VAF, YAF and MSF variants were successfully expressed and displayed thermostability comparable to wild-type ecPGA (Figure S2).^39,40^ Further, these variants were subjected to a hydrolytic activity assays with AHLs and PenG. Experimental assays indicated markedly modulated activities of the mutants towards AHLs (Table 1 and Supplementary File 3). Moreover, all variants were incapable of degrading PenG in comparison with the wild-type enzyme (activity 2600 ± 50 U/L and specific activity 1020 ± 75 U/gcdw), indicating notable shift in preference from antibiotics to bacterial signaling molecules. In order to emphasize the importance of the observed changes, we compared the ability of the purified VAF, YAF and MSF variants with wild-type ecPGA to catalyze kinetically controlled synthesis of ampicillin and amoxicillin, a pair of the most produced semi-synthetic B-lactam antibiotics by ecPGA.^41^ We observed reduced capabilities of VAF, YAF and MSF to synthesize ampicillin and amoxicillin as compared to wild-type ecPGA (Figures S3-S4 and Table S7). Regarding activity towards AHLs, the introduced mutations had a primarily neutral effect on enzyme activity with shorter substrates compared to wild-type ecPGA. The only exception was the YAF mutant, which exhibited a significant drop of the catalytic efficiency below 10% of the wild-type, caused both by the decreased *kcat* and increased *KM*. This observation contrasts with the computational prediction of the neutral effect, indicating that the activity drop originates from subsequent reaction stages not investigated in the simulations. Overall, we observed satisfactory agreement between the experimental assay and predictions for medium and long substrates. The YAF mutant performed well with the long substrates, reaching a 3-fold improvement for C08-3O- and C12-3O-HSLs. The VAF mutant exhibited notably improved activities with medium substrates, i.e., a nearly 2- and 5-fold improvement for C08-3O- and C08-HSLs, respectively. Interestingly, the VAF mutant had relatively low activity for long substrates, similar to the MSF mutant with C10-HSL, which, conversely, performed well for the longer C12-3O-HSL. These differences were subsequently supported by a *Chromobacterium violaceum* CV026-based bioassay used to confirm the activity of the three mutant variants as QS signal quenchers (Figure S5). Intrigued by these tendencies we investigated the origins of these non-trivial effects in more detail.

**Table 1.** Experimental results of enzyme kinetic parameters for wild-type and designed ecPGA variants.

| AHL/variant | wild-type ecPGA | VAF | YAF | MSF |
| --- | --- | --- | --- | --- |
| **$k_{cat}$ [$s^{-1}$]** |  |  |  |  |
| C06 | 0.0090 ± 0.0003 | 0.0096 ± 0.0010 | 0.0031 ± 0.0002 | 0.0099 ± 0.0006 |
| C06-3O | 0.0100 ± 0.0002 | 0.0098 ± 0.0006 | 0.0030 ± 0.0003 | 0.0090 ± 0.0005 |
| C08 | 0.0085 ± 0.0003 | 0.0178 ± 0.0010 | 0.0147 ± 0.0010 | 0.0098 ± 0.0009 |
| C08-3O | 0.0077 ± 0.0004 | 0.0091 ± 0.0010 | 0.0120 ± 0.0010 | 0.0110 ± 0.0008 |
| C10 | 0.0045 ± 0.0004 | 0.0045 ± 0.0004 | 0.0040 ± 0.0002 | 0.0044 ± 0.0004 |
| C12-3O | 0.0043 ± 0.0002 | 0.0041 ± 0.0003 | 0.0055 ± 0.0002 | 0.0050 ± 0.0006 |
| **$K_M$ [mM]** |  |  |  |  |
| C06 | 0.59 ± 0.07 | 0.52 ± 0.05 | 2.42 ± 0.15 | 0.49 ± 0.01 |
| C06-3O | 0.51 ± 0.03 | 0.53 ± 0.03 | 1.34 ± 0.09 | 0.62 ± 0.02 |
| C08 | 0.70 ± 0.04 | 0.28 ± 0.03 | 0.38 ± 0.03 | 0.49 ± 0.01 |
| C08-3O | 0.88 ± 0.04 | 0.57 ± 0.02 | 0.41 ± 0.04 | 0.42 ± 0.01 |
| C10 | 2.98 ± 0.20 | 2.98 ± 0.20 | 1.44 ± 0.08 | 3.01 ± 0.30 |
| C12-3O | 2.50 ± 0.10 | 2.81 ± 0.20 | 1.06 ± 0.04 | 1.12 ± 0.20 |
| **$k_{cat}/K_M$ [mM$^{-1}$ s$^{-1}$]** |  |  |  |  |
| C06 | 0.0153 ± 0.0010 | 0.0185 ± 0.0020 | 0.0013 ± 0.0006 | 0.0200 ± 0.0040 |
| C06-3O | 0.0196 ± 0.0020 | 0.0185 ± 0.0040 | 0.0022 ± 0.0005 | 0.0146 ± 0.0040 |
| C08 | 0.0122 ± 0.0020 | 0.0636 ± 0.0040 | 0.0387 ± 0.0040 | 0.0200 ± 0.0030 |
| C08-3O | 0.0087 ± 0.0010 | 0.0160 ± 0.0020 | 0.0300 ± 0.0010 | 0.0262 ± 0.0010 |
| C10 | 0.0015 ± 0.0002 | 0.0015 ± 0.0003 | 0.0028 ± 0.0008 | 0.0015 ± 0.0003 |
| C12-3O | 0.0017 ± 0.0003 | 0.0015 ± 0.0003 | 0.0052 ± 0.0009 | 0.0045 ± 0.0005 |

The data is represented as the mean ± stdev.

### Engineered variants maintain their ability to open the entrance to the acyl-binding pocket and arrange their catalytic machinery for the productive binding of AHLs but prohibits proficient hydrolysis of native substrate PenG

Interestingly, comparison of the experimental results and computational predictions formulated in a form of RSS exhibited good correspondence between the ranking provided by our computational predictions and experimental *KM* values, with a Spearman correlation coefficient of -0.69 (Figure S6), suggesting the developed score as a relevant metric to guide rational engineering of penicillin G acylases. Furthermore, we noticed lack of rank correlation of our metric with *kcat* or *kcat*/*KM* parameters indicating that the following reaction steps not considered in our simulations might be rate limiting (Figures S7-S8).^42^

To determine the molecular basis of substrate specificity change upon the introduced mutations, we performed simulations of VAF, YAF, MSF variants as well as wild-type ecPGA with PenG (Figures S9-S12). Interestingly, we observed RSS of 0.33, 0.95 and 0.43 for VAF, YAF and MSF variants, respectively, accounting for nearly 5- to 4-fold decrease in this metric for VAF and MSF mutants as compared to 1.74 RSS detected for ecPGA, confirming consistency between computational predictions and experimental assays (Table S8). Next, we followed with free enzyme MD simulations of 0.5 µs in length for three replicates for each variant and contrasted those results with previously produced simulations of ecPGA and paPvdQ.^29^ The simulations indicated that engineered variants maintained the overall folds well, confirming the absence of severe destabilizing effects due to the introduced mutations (Figure S13), consistently with experimentally derived melting temperatures comparable to wild-type enzyme (Figure S2). Furthermore, we inspected the tendency of the catalytic machinery, consisting of Ser1β, Ala69β, Asn241β, and Gln23β, to adopt properly pre-organized conformations for productive stabilization of AHLs. Analogously to our previous work,^29^ we measured distances between functional atoms of these residues and performed principal component analysis (PCA) to characterize conformations adopted by these crucial residues (Figure S14). These residues often adopt the favorable state for productive binding, characterized by the serine hydroxyl oxygen approximately equidistant to the remaining functional (stabilizing) atoms. This state was visited in 59 % and 51 % of YAF and MSF simulations, respectively, even surpassing the 36% occurrence in paPvdQ simulations (Table S9). For mutant VAF, the appropriate state was not dominantly adopted, with approximately 21% occurrence, similar to 24 % occurrence in ecPGA (Table S9). Additionally, we inspected the capabilities of the acyl-binding site opening to track potential undesirable gating events introduced by mutations. From this perspective, all variants exhibited sufficiently frequent openings to bind AHLs comparable to wild-type ecPGA (Figures S15-S18).

The engineered variants demonstrated uniform expansion of their pockets, although this altered the dynamics of their entrances and the stability of the deepest pocket regions. To evaluate the effect of mutations on the targeted acyl-binding pockets, we compared their behavior between the variants, wild-type ecPGA and paPvdQ. Starting from the average volumes that we found in the simulations, all engineered pockets were successfully expanded over the volume observed for the ecPGA pocket, often reaching or even exceeding the volume of the paPvdQ pocket (Figure 5A). By inspecting the details of the geometrical profiles of the pockets (Figure 5B and Figure S19), we observed increased pocket depth in all variants, almost reaching the depth of pocket in paPvdQ, and approximately twice as deep as the ecPGA wild-type.

**Figure 5.**
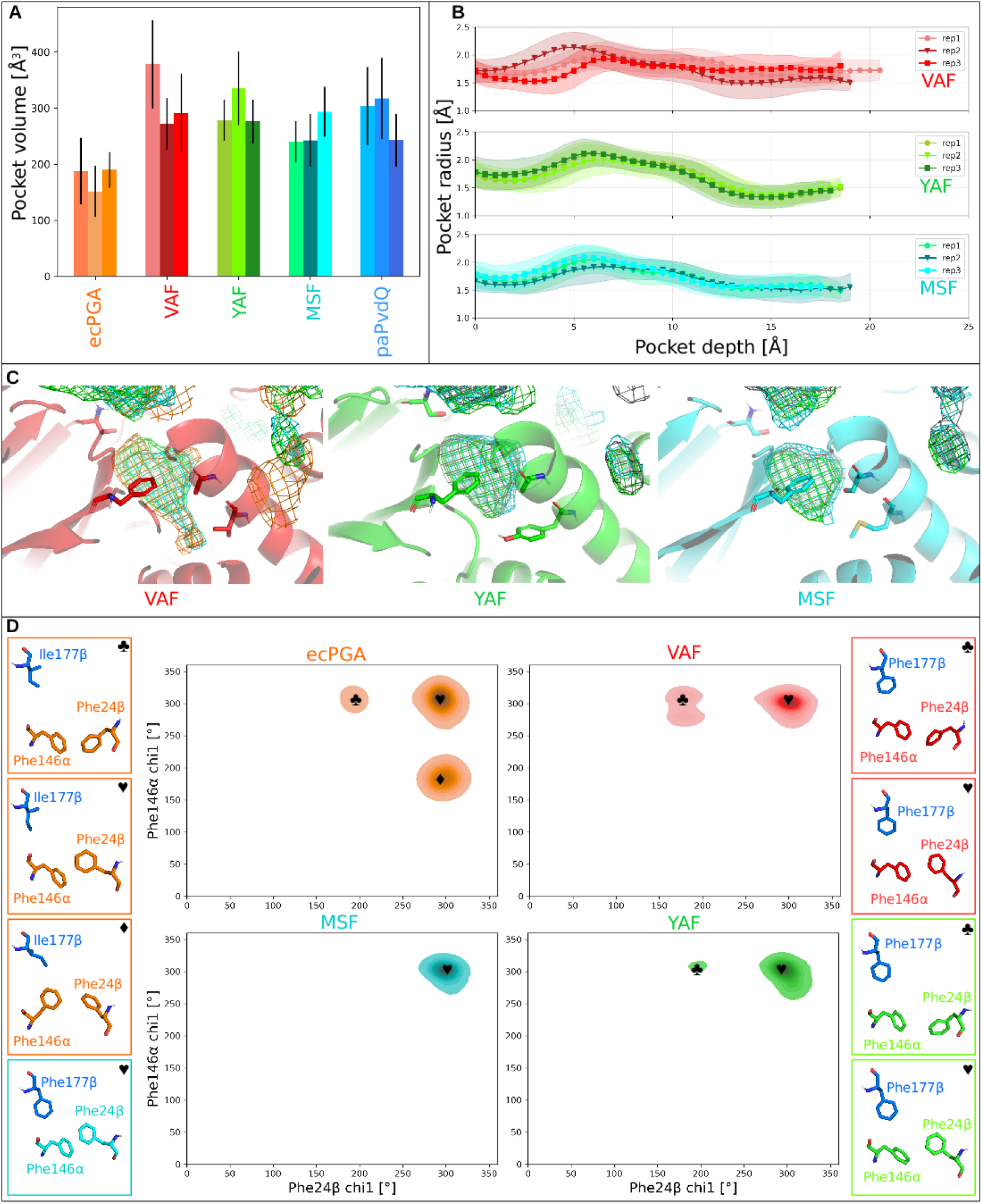
Detailed investigation of structure and dynamics of engineered binding pockets. **A**) Average binding pocket volumes calculated from free enzyme MD simulations using mdpocket. **B**) Profiles of the binding pocket in the free enzyme MD simulations, based on CAVER tunnel calculations analyzed from properly pre-organized structures. **C**) Isomesh represents the open voids that occurred in at least 50 % of analyzed frames, with mutated residues shown as sticks. **D**) Modulated dynamics of the gating residues of designed mutants compared to wild-type ecPGA enzyme. Graphs show conformations sampled by the gating residues as maps of chi1 angles of Phe24β versus Phe146α. Structures located in frames around the maps show (sticks) representative conformations for each state.

Interestingly, the VAF mutant pocket presented the least stable profile, particularly at the bottom portion (the depth of 15-20 Å). Here, structural fluctuations resulting in the opening of extra space were regularly observed, while well-defined and stable pocket bottoms were present in the other two mutants (Figure 5C). To understand the potential source of the differential pocket stability, we measured the distances between the three pairs of mutated residues in all variants (Figure S20). This data displays different behavior of mutated sites, i.e., mutants YAF and MSF present analogous behavior, where the residues at the 138α and 142α positions formed the closest pair, followed by the 142α-177β pair, and finally the 138α-177β pair. Conversely, in the mutant VAF, we observed two essential differences, namely (i) separation between the 138α-142α pair was increased, similarly to wild-type, reaching the ranges seen for the 142α-177β pair, and (ii) the distances between both the 138α-142α and 138α-177β pairs were less stable, often adopting much larger distances. This more dynamic cross-talk between substituted residues correlates with the observed increased fluctuation of the binding pocket volume for this mutant (Figure 5A) and more considerable deviation in the bottom part of the pocket, as illustrated by the profile analysis (Figure 5B).

Additionally, the initial part of the pocket profile in the engineered variants provided intriguing observations (Figure 5B). Here, we noticed a constriction located below the 5 Å depth that was not present in wild types. Since the constricted area was within the proximity of the mutated site (177β), we analyzed the effect of this perturbation on the conformational dynamics of gating residues Phe24β and Phe146α. The wild-type ecPGA is known to frequently visit two main states of Phe146α (♥ and ♦) while occasionally also adopting an alternative conformation of Phe24β (♣).^29^ Interestingly, we observed markedly restricted dynamics of the gating residues in all three variants (Figure 5D), predominantly displaying the single conformation of Phe146α (♥). While the alternative conformation of Phe24β (♣) was rarely visited in the case of VAF and YAF variants, The MSF variant did not display this conformation. To understand how the Ile177βPhe substitution affects the behavior of the gate, we inspected the distribution of spatial separation between the center of mass of the 177β residue and the two gating residues (Figure S21). The most striking difference was the limited range of the distances between gating Phe146α and the introduced Phe177β observed in all mutants when compared to the wild-type ecPGA, where the separation between these two residues reached over 10 Å. Curiously, the restricted mobility of this pair was less pronounced in the VAF mutant, in which the distribution of the respective distances was broader and exhibited bimodal behavior. The increased mobility of the entrance and the initial part of the pocket in this mutant is in agreement with the increased fluctuations detected for this mutant in the pocket profile, volume and residues forming the bottom part of the cavity, rendering this pocket the most dynamic and unstable among the evaluated variants.

### Suboptimal dynamics of the enlarged pockets prevent productive stabilization of longer AHLs in VAF and MSF mutants

One of the most intriguing discrepancies between the structural analysis of the investigated variants and their preferences for various AHLs is the minimal activity of the VAF mutant that was observed with C10- and C12-3O-HSLs, despite its most extensive and flexible binding pocket. Our previous analyses of the ecPGA-C08-HSL complex identified the acyl-binding pocket as being too shallow, causing suboptimal stabilization of this substrate by the oxyanion hole residues. The increased distance of C08-HSL to Ala69β (from 2.0 to 2.5 Å) was found to be a relevant indicator reflecting the increased tendency of the substrate to leave the binding site.^29^ Hence, we analyzed these distances in our simulations to evaluate the proper stabilization of AHLs in all variants (Table S10). In line with our expectation, the VAF mutant exhibited markedly longer distances to the oxyanion-hole stabilizing Ala69β with both of the longer substrates, reaching 2.9 ± 0.1 and 2.6 ± 0.3 Å for C10- and C12-3O-HSL, respectively. These distances were much higher when compared to the remaining active complexes. Unlike in our previous study with ecPGA-C08-HSL, the instability of longer AHLs was not caused by a problematic fit into a too-shallow pocket, but instead by the increased dynamics of the pocket.

Additionally, we observed a similar increase in the Ala69β distance for these AHLs with the MSF mutant (Table S10). The pocket of this mutant remained stable following the introduction of Ser142α, as this residue made frequent hydrogen bonds with Tyr52β, stabilizing the bottom section of the pocket. We envision two possible mechanisms behind the enhanced expulsion of longer AHLs from the binding pocket of MSF mutant, being i) the stability introduced into the pocket could limit the range of adjustments necessary to accommodate such AHLs, as corroborated by the conformational changes observed at the bottom of the paPvdQ pocket upon C12-HSL binding,^30^ and ii) such an arrangement results into the exposure of multiple polar groups inside the pocket, decreasing complementarity with the hydrophobic tails of the bound C10- and C12-3O-HSL. Surprisingly, these effects translated to the less frequent formation of a reactive binding pose and lower activity for only the C10-HSL, while the combined inference into C12-3O-HSL showed improved catalytic performance, which is quite controversial considering that the later AHL is considerably longer. Attempting an explanation of this inconsistency, we realized that although both C10- and C12-3O-HSL represent rather long substrates, they also differ by the presence of a 3-oxo substituent. Thus, we analyzed hydrogen bonding interactions formed with this group. Aside from rare interactions with Ser1β and Gln23β, the most frequent interaction of the 3-oxo group was noted with the amide group of Ala69β (Table S11). When inspecting the occurrence of these hydrogen bonding events and the reactive complexes, we noted the recurring recovery of reactive poses after the hydrogen bonds are formed with AHLs carrying the 3-oxo substituent (Supplementary File 4 and Figure S22). This observation indicates that although the binding pocket carrying Ser142α is not optimal for the longer substrates, the 3-oxo substitution in C12-3O-HSL, absent in C10-HSL, partially offsets the repulsion, helping the substrate stay in the vicinity of the catalytic machinery and promoting the restoration of the productive complex. This is further supported by analogous behavior observed in simulations of the LSF mutant carrying the same mutation (Figure 4).

## DISCUSSION

Enzymes carrying AHL-degrading activity have gained significant interest as promising antimicrobial agents in the last decade, mainly due to their environmentally friendly character, potential to escape common resistance mechanisms, and their application potential in medicine, various fields of industry, biotechnology, and agriculture.^14,15^ Furthermore, the current antibiotic discovery void accentuates the need to investigate antibiotic alternatives to effectively fight bacterial infections.^1,10^ Several studies have previously investigated engineered AHL-degrading acylases, examining improvements in QQ potency or modified substrate specificity, including PvdQ acylase,^19^ MacQ bifunctional acylase,^43^ and PF2571 AHL acylase^22^. Furthermore, significant efforts have been invested in developing and improving methods for the efficient engineering of these enzymes for effective QQ activity.^44,45^ However, to our knowledge, most of the previous systems are not yet fully developed for efficient and sustainable applications on a broader scale.

To address this issue, and building on top of our previous findings of AHL-degrading activity of industrially potent penicillin G acylases towards short- to medium-size AHLs,^29^ we engineered ecPGA specificity towards a group of AHLs of different sizes. We followed a workflow for the computational design of pocket ensembles facilitated by the recently developed TransportTools library^35^ that enabled us to compare differences in pocket geometries across the large dataset in a unified and reproducible manner. We used this workflow to introduce triple-point mutations in ecPGA’s binding pocket to alter its ability to accommodate AHLs while deprioritizing mutants with increased destabilization. Furthermore, the RSS that we developed to select candidates from MD simulations for experimental evaluation displayed an appreciable correlation to the results of biochemical assays, suggesting its ability to screen for analogous enzymes displaying AHL-degrading activity (Figure S6). Moreover, the computer designs have marked potential, considering that experimentally they displayed the most notable improvement of the activity towards particular bacterial signaling molecules, namely, the VAF mutant being most active for C08-HSL used by *B. cenocepacia*, the YAF mutant displaying the best improvement for C10-HSL, the signaling compound of pathogenic *Vibrio* species, and finally, the MSF mutant the most active towards C12-3O-HSL, the signaling molecule of *P. aeruginosa*, and the preferred substrate of prototypical paPvdQ (Table 1). In parallel, activities of all constructed mutants with the native substrate of ecPGA – PenG become experimentally undetectable. Nonetheless, computational inference into reactive stabilization of these complexes enabled us to estimate the magnitude of the shifts in substrate preference from wild-type ecPGA caused by the designed mutations (Tables S6 and S8): VAF ∼ 125 times (C08-HSL/PenG), YAF ∼ 44 times (C10-HSL/PenG), and MSF ∼ 1287 times (C12-3O-HSL/PenG).

Detailed analyses of three experimentally constructed variants exposed changes in the structure-dynamic-function relationships that are relevant to the specificity of these enzymes towards AHLs of different lengths. The VAF mutant, which lacked enhanced activity with longer AHLs and presented the highest dynamics and plasticity of the binding site, was also the variant with the highest predicted destabilization energy in the initial design despite its maintained thermal stability (Figure S2). This highlighted the multifaceted consequences of such local destabilizing effects during rational engineering campaigns. Conversely, YAF and MSF mutants featured stable binding pockets capable of proper stabilization of longer AHLs, although we also identified drawbacks of potential overstabilization of the pocket in variants carrying Met142αSer mutation. Furthermore, we detected modulated dynamics of the Phe146α and Phe24β gating residues compared to wild-type ecPGA, tracking the source of this perturbation to the introduction of the Phe177β residue, which interacts more closely with the gating residue Phe146α, restricting its conformational freedom. Although limiting the dynamics of the gate and driving it closer to more stable gates observed in the prototypical paPvdQ,^29^ the perturbation had not narrowed down the entrance as designed. Instead, the entrance remained as open as that seen in the wild-type, whereas the subsequent part of the pocket became more constricted (Figures 2C and 5B). We hypothesize that this constriction may instead compromise the efficient binding of the AHLs that need to adopt strained conformations to fit such a curved pocket. Although revealing unfavorable effects that were introduced with the Ile177βPhe substitution, this uncovered an additional hot-spot contribution to the dynamic control of gating in ecPGA, opening new opportunities for modulation of AHL-degrading activity and specificity, bringing us closer to efficient QQ agents. When all our data is taken into account, we have demonstrated the ability to design industrially proficient ecPGA with shifted specificity from native substrate penicillin G to bacterial signaling molecules of different acyl-chain length. Importantly, although the AHL-degrading activities for designed mutants remain basal and their further QQ potential has to be validated on bacterial colonies, we have demonstrated the ability to modulate the specificity of these enzyme for desired bacterial signaling compounds, and the identified several relevant targets enabling further engineering of ecPGA, leading for efficient antibacterial agents with practical application potential.

## METHODS

### Computational

#### Preparation of templates for mutagenesis

The three best quality structures of ecPGA were selected from the PDB database (PDB IDs: 1GK9, 1GM7, and 1GM9),^46^ and water molecules, and organic and inorganic compounds were removed. Structures were further modified to achieve a wild-type composition; 1GM7 Ala241β was converted to native Asn, and Sme16α was replaced by Met for 1GM7 and 1GM9 structures using PyMOL 2.0.1 (The PyMOL Molecular Graphics System, Version 2.0 Schrödinger, LLC.) mutagenesis wizard. The *RepairPDB* module of FoldX 4^34^ was then applied to improve the quality of the structures, indicated in the structures validation reports.

#### *In-silico* site-saturation mutagenesis and filtering

The *BuildModel* module of FoldX 4 software was used to construct the mutant structures. Each mutagenesis experiment was performed using the module’s default settings, namely ionStrength 0.05, pH 7, temperature 298K, vdWDesign 2.0, and was repeated ten times (numberOfRuns 10). The predicted stability of the introduced mutation was averaged from the mutagenesis obtained independently for each of the structures, considering both orders of introducing mutations for two-point mutations. Overall, an ensemble of 60 structures was generated for each mutant (30 x 2) and further evaluated with CAVER 3.02^47^ software to calculate the tunnel covering the binding pocket entrance. The starting point for the CAVER calculations was set up as a center of mutated residues, namely Phe138α, Met142α, and Ile177β, with *probe_radius* of 0.9 Å, *shell_radius* of 3 and *shell_depth* set up to 7. The results of the CAVER calculation for each mutant were processed separately by the comparative analysis module of TransportTools 0.9.4 software.^35^ Tunnel clusters were joined into superclusters using the average-linkage clustering method with a 4.0 Å cutoff. Filtering was applied to consider only superclusters containing tunnels that occurred in at least two structures per ensemble that were generated for each variant (*min_snapshot_num = 2*). The variants were then evaluated based on the stability estimated by FoldX software and the geometric filtering based on comparative TransportTools analysis. The variables we considered were the number of frames with an open pocket, average length, average and maximal bottleneck radii, and average throughput.

#### Preparation of protein and substrate structures for molecular docking

At this stage, only the variants we obtained from the best-quality template of the wild-type enzyme (PDB-ID 1GK9) were used for further modeling. Designed and repaired wild-type structures were protonated by H++ web-server^48,49^ with default settings, namely pH 7.5, salinity 0.15, and internal and external dielectric constants of 10 and 80, respectively. Protonation of the crucial Ser1β residue was adjusted to resemble the state ready for the first stage of the reaction. Autodock4.2 was used to perform molecular docking experiments to obtain protein-substrate complexes for all designed variants and the wild-type with the following set of AHLs: C06-HSL, C06-3O-HSL, C08-HSL, C08-3O-HSL, C10-HSL, and C12-3O-HSL and PenG. Structures of AHLs were created in Avogadro,^50^ while the PenG was obtained from penicillin G acylase crystal structure with its complex.^46^ Molecules were optimized using Gaussian09 software^51^ at HF/6-31G* level of theory. Furthermore, charges were derived from the RESP-A1 charge model with a multi-conformational RESP fitting procedure using the RESP ESP web server.^52,53^

#### Molecular docking calculations

The grid box for docking was defined using the center of mass of Ser1β and Phe138α residues and was adjusted manually using Autodock PyMOL plug-in^54^ to cover the entire binding site and catalytic machinery. The box dimension was set up to be 17.5 x 16.25 x 21.25 Å in size. Substrate molecules were treated as flexible during docking, while the receptor was considered rigid except for the mutated residue at position 138α. Receptor and substrate structures were prepared using *prepare_receptor4.py* and *prepare_ligand4.py* scripts from AutoDockTools4 utilities.^55^ Docking procedure was performed using the default settings of the Lamarckian Genetic Algorithm search, which was repeated 250 times for each complex. Representative complex structures were selected based on the favorable binding score from the Autodock4.2 scoring function and were filtered by the criteria defined by proper substrate stabilization for the nucleophile attack reaction.^27,29,36^ This criteria included oxyanion hole stabilization interactions (distances between the carbonyl oxygen of the substrate molecule and hydrogens from Ala69β-NH and Asn241β-NδH), nucleophile attack (distance between carbonyl carbon of the substrate and hydroxyl oxygen of Ser1β), and additional stabilization provided by Gln23β (distance between amide hydrogen of the substrate and backbone oxygen Gln23β-O) that is known to play a crucial role in the tetrahedral intermediate stabilization. Complexes with distances satisfying a cutoff of 3.0 Å and favorable binding score were used as inputs for the following MD simulations.

#### Molecular dynamics simulations of protein-substrate complexes

Pre-selected complex structures were merged with the crystallographic waters, and placed in an octahedral box containing TIP3P water molecules with a minimum distance of 10 Å between the solute and the edge of the box. Na+ and Cl-ions were added to the system to neutralize the charge and obtain a salt concentration of approximately 0.1 M. Water molecules that were too close to the solute were removed to avoid potential instabilities in the system during simulation. Parameters and topologies were obtained using the *tleap* module from AmberTools17^56^, applying the ff14SB force field.^57^ The protonation of the N-terminal serine was designed to resemble the nucleophile-attack ready state (^+^H3N-Ser-O^-^).^29,58^ Charges were obtained analogously to the procedure performed for substrates, and the different atom types were derived analogously to the Cornell et al. force field.^59^ Energy minimization and MD simulations were performed using the *pmemd* of AMBER 16 package for complexes with HSLs and pmemd/*pmemd.CUDA* engines of AMBER 18 package for complexes with PenG.^56,60^ All complexes were energy minimized in five consecutive stages, each composed of 1000 steps of the steepest descent method followed by 1000 steps of the conjugate gradient method, with gradually decreasing restraints on the protein heavy atoms (initially 500 for heavy atoms, and later restraints applied only on the backbone atoms 500, 125, 25, 0.001 kcal/mol^-1^ Å^-2^) and substrate (initially 500 for heavy atoms, continued by restraints used only on the amide bond atoms, 50, 25, 5, 0.01 kcal/mol^-1^ Å^-2^). Equilibration was performed in 4 stages: (i) 100 ps NVT heating stage from 0 to 100 K using a Langevin thermostat with 5 kcal/mol^-1^Å^-2^ harmonic restraints on the heavy atoms, followed by three NPT stages using a Langevin thermostat and Monte Carlo barostat: (ii) 300 ps in 100 K with 5 kcal/mol^-1^Å^-2^ harmonic restraints on the heavy atoms, (iii) 600 ps gradually heating the system to 310 K in the first 200 ps applying harmonic restraints on the backbone heavy atoms using force constant of 5 kcal/mol^-1^Å^-2^ and distance restraints on the crucial hydrogen bonds stabilizing the substrates (substrate carbonyl carbon to oxyanion hole stabilizing residues Ala69β-NH and Asn241β-NδH) with additional stabilization of the substrate amide hydrogen (by Gln23β-O) using force constant of 25 kcal/mol^-1^ Å^-2^, and (iv) 1 ns gradually decreasing the distances that restraint was applied in stage (iii) from 25 to 0 kcal/mol^-1^Å^-2^. Furthermore, 5 and 50 ns NPT production runs were performed for AHL and PenG complexes, respectively, using a Langevin thermostat at 310 K, applying a Monte Carlo barostat. All MD simulation stages were run using a 2 fs time-step. The trajectory was generated by saving coordinates every 10 ps, and simulation steps (minimization, equilibration) were performed at least 30 times, with the exception of the production, which was continued only for the replicates that maintained the substrate in close proximity to the productive state and properly stabilized by the active site machinery. Otherwise, such replicates were considered non-productive (with zero frames sampling reactive stabilization of the substrate), and terminated after equilibration stage. Root mean square deviation (RMSD) of protein and substrate separately were used to inspect the stability of the system and complex itself during production runs (Supplementary File 2 and Figures S9-S12). Crucial distances required for estimating enzymes’ capabilities to maintain reactive stabilization of the complex were measured using *cpptraj* from AmberTools17. Heat maps representing the relative change of the potency of each variant were generated using the Seaborn Python library.^61^ RSS was defined according to following equation:

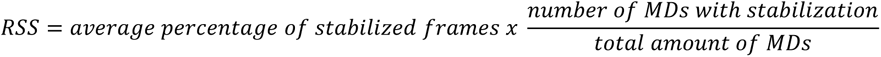

Where: *average percentage of stabilized frames* – average percentage of the frames with the proper stabilization sampled, calculated from all replicates for particular complex; *number of MDs with stabilization* – total amount of simulated replicates for particular complex in which the proper stabilization was detected; *total amount of MDs* – overall amount of simulated replicates performed for particular complex.

#### Molecular dynamics simulations of free enzymes

Wild-type ecPGA and paPvdQ enzyme structures (PDB ID: 1GK9 and 4M1J^62^), as well as the three best mutants (VAF, YAF, and MSF), were subjected to an additional round of free enzyme MD simulations. Systems were prepared in an analogous way to protein-substrate complexes described in the previous section with three major differences: (i) a lack of substrate in the system, (ii) neutral (resting) N-terminal serine protonation state (H2N-Ser-OH)^29,58^ and (iii) hydrogen atom masses were repartitioned^63^ to enable 4 fs time-steps during MD simulations.

Parameters for this neutral protonation state were obtained from our previous study.^29^ The minimization was performed with the *pmemd* module of Amber18,^60^ while equilibration and production runs were performed with the *pmemd.CUDA* module of the same package, using analogous settings as for the protein-substrate complexes. All stages were performed in triplicate, and the production simulation was 500 ns long, with the trajectory generated by saving coordinates every 20 ps. The stability of the systems was inspected in terms of RMSD (Figure S13). Crucial distances between catalytically relevant atoms were measured using the *cpptraj* module of AmberTools18. Then, PCA was applied to perform dimensionality reduction of this distances network, as implemented in Python scikit-learn library.^64^ First, two principal components (PC1 and PC2) were clustered using hdbscan^65^ with *min_cluster_size* of 500 and *cluster_selection_epsilon* of 0.001. From these results, members featuring the strongest affiliation to a particular cluster (*probability_* = 1.0) were used to calculate average distances that characterize the cluster representatives, which were further transformed to the original PC space using the PCA model.

#### Characterization of binding site properties

*Cpptraj* from AmberTools18 was used to generate an ensemble of every tenth snapshot from free enzyme trajectories (2500 snapshots per trajectory) in PDB file format for further geometrical analysis. First, a pocket search was performed using mdpocket^33,66^ run (mode 1) with default settings for wild-type and the three shortlisted mutant structures. Then, grid points corresponding to the binding pocket cavity of interest were selected, and the following run (mode 2) was performed to calculate the pocket descriptors. Additionally, mdpocket-generated information on the most conserved residues that form particular pockets was used to select the starting point for tunnel detection using the CAVER software.^47^ Those covered the center of mass of the residues Val138α, Val56β and Trp154β for mutant VAF; Tyr52β, Val56β and Trp154β for mutants YAF and MSF; Met142α, Ser67β and Ile177β for ecPGA wild-type; and Leu146α, Leu53β, and Trp162β for paPvdQ. Tunnel calculations were performed using a probe radius of 0.7 Å. The clustering threshold was 3.5 Å, and one representative was selected for each path. Profiles to visualize binding cavities were generated based on filtered CAVER paths, and we considered only those that corresponded to frames with cavities meeting the following criteria; (i) minimum depth of the cavity greater than 5 Å, (ii) the narrowest point at the entrance to the cavity is greater than 1.4 Å, and (iii) the maximum distance between the desired and actual starting point for tunnel calculations and end point of the tunnel below 2.0 Å.

All plots were generated using matplotlib^67^ or seaborn Python libraries. Figures containing structural representations were generated using PyMOL 2.0.1.

### Experimental

#### Chemicals

The standard of C10-HSL (molecular weight (MW) of 255.4) was obtained from Merck AG (Darmstadt, Germany). The remaining AHLs, namely C06-HSL (MW of 199.2), C06-3O-HSL (MW of 213.2), C08-HSL (MW of 227.3), C08-3O-HSL (MW of 241.3), and C12-3O-HSL (MW of 297.4) were purchased from Cayman Chemical (Michigan, USA) and prepared as 10 mM solution dissolved in DMSO and stored in aliquots in -20 °C. The chemicals and ingredients used in the microbiological experiments were obtained from Sigma-Aldrich (St. Louis, MO, USA). The antibiotics such as ampicillin, amoxicillin, and kanamycin were purchased Sigma-Aldrich (Missouri, USA). The other chemicals and ingredients used in the analysis were of high analytical grade and were obtained from Merck (Darmstadt, Germany).

#### Bacterial strains and growth conditions

*E. coli* BL21(DE3), *E. coli* TOP10 (for routine cloning) and the control strain RE3(pKA18),^66^ were cultivated in Luria-Bertani (LB) medium (in g/L: tryptone 10, yeast extract 5, NaCl 10, pH 7.2) supplemented with kanamycin (35 µg/mL) for 24 h at 37°C, 180 rpm. *Chromobacterium violaceum* CV026 (CECT 5999), the AHL biosensor strain, was used to screen quorum sensing inhibitory activity and was cultured aerobically in LB medium supplemented with kanamycin (25 µg/mL) for 24 h at 28°C, 180 rpm. To produce mutated ecPGAs, recombinant strains were cultivated in 50 mL of mineral MCHGly medium for 24 h at 28°C, 180 rpm, and protein expression was induced with 1 mM isopropyl alcohol β-D-thiogalactopyranoside (IPTG).^67^ The AIM2YT Broth Base medium (2YT) enriched with trace elements (Formedium Ltd., Norfolk, UK) was used for melting point temperature determination. To prepare solid media for screening of recombinant strains and short-term strain maintenance, LB media were supplemented with agar (15 g/L).

#### Construction of expression system and mutagenesis

The *ecpga* gene was initially cloned in the pET-26b vector using restriction cloning. In brief, the *ecpga* gene (GenBank: X04114.1) was amplified by PCR using Q5 polymerase (NEB, USA) from our previous plasmid pKA18^68^ with ecPGA cloning primers bearing *Nde*I and *Xho*I site overlaps. The cloning primer for His-tagged PGA variant was designed without stop codon sequence. The purified (NucleoSpin® Gel and PCR Clean-up, Macherey-Nagel, Germany) and digested PCR product by *Nde*I and *Xho*I restriction enzymes (NEB, USA) was ligated using T4 DNA ligase (NEB, USA) into the pET-26b vector cleaved with the same enzymes. The construct was directly transformed in electrocompetent *E. coli* TOP10 cells. Single colonies were screened by colony PCR and verified by sequencing with sequencing primers. Mutagenesis steps were performed by implementing the restriction-free (RF) cloning method.^69^ Initially, mega-primers were amplified by PCR reactions (KAPA HiFi DNA Polymerase, Roche, Switzerland) with specific primers bearing mutations and the reverse primer overlapping the integration sites in the destination vector. Gel-purified mega-primers (NucleoSpin®) were incorporated in the destination plasmid pET-26-PGA by second whole-plasmid amplification PCR. The parental vector was digested by the *Dpn*I enzyme (NEB, USA), and the ligation mixture was directly electroporated into *E. coli* TOP10 cells. The successful incorporation of the mega-primers into the destination vector was verified by colony PCR and sequencing. All primers are listed in Table S12. The constructed plasmids bearing the mutated *pga* gene were purified from *E. coli* TOP10 cells and transformed to the *E. coli* BL21(DE3).

#### Enzyme purification and hydrolytic assay

The three chosen mutants were purified as previously described by Kutzbach and Rauenbusch.^70^ The activity of one unit (U) was defined as the amount of ecPGA cleaving 1 µmol of the corresponding AHL per minute in 0.05 M sodium phosphate buffer containing 2 % (w/v) AHL at 35°C and pH 7.0.

#### Biosensors assay of ecPGA for AHLs

To screen QQ enzyme activity, cells grown in MCHGly medium with 1 mM IPTG were separated by centrifugation (8000 g, 5 min, 4 °C), washed by with 0.05 M Na phosphate buffer (pH 8.0) and sonicated (using a Sonicator 3000, Misonix, at 15-20W). The cell-free extract obtained after centrifugation (8 500 x g, 5 min, 4°C) was used in screening biosensor assays with the *C. violaceum* strains CV026 and 300 µM C06-HSL, C08-HSL and C08-3O-HSL according to Last et al.,^71^, i.e., 40 µL of cell-free extract was mixed with 300 µM AHL in LB medium in an Eppendorf tube, and the mixture was incubated for 1 h at 37 °C and 100 rpm. After the biocatalytic reaction, 160 µL of diluted *C. violaceum* CV026 (in the ratio of 1:50) from an overnight culture grown in LB medium supplemented with kanamycin (25 µg/mL) was added to the reaction mixture.

#### Effect of temperature on the activity of AHL degradation

The optimal temperature for the activity of individual ecPGA variants was determined by conducting cleavage experiments as described in detail in Surpeta *et al*.^39^ for all the AHLs signaling molecules investigated herein.

#### Enzyme kinetics

Kinetic characterization of AHL degradation by ecPGA and its mutant variants was carried out in 0.05 M phosphate buffer at pH 7.0 at the optimal temperature for ecPGA (35 °C), as described previously.^39^ Monitoring of reactant concentration was performed at 210 nm using an HPLC apparatus, and samples were prepared for HPLC analyses as follows. First, the aliquot reaction was adjusted to pH 2 to terminate the reaction, followed by evaporation to dryness at 35°C. The resulting residues were dissolved in 0.2 mL of acetonitrile. An aliquot of 20 μL of the resulting solution was loaded onto an analytical RP-C18 HPLC column (250 x 4.6, 5 μm particle size, Hypersil ODS). The elution profile proceeded under the following conditions: i) isocratic elution in methanol-water (50:50, v/v) for 10 minutes, ii) linear gradient from 50 to 90 % methanol when in water for over 15 minutes, iii) isocratic elution for 25 minutes. The flow rate for HPLC was set at 0.4 mL/min. The retention times of the substrates of interest are listed in Table S13. The relationship between the initial reaction rate and the substrate concentration (1 - 1000 µM) was determined for each substrate in three independent experiments. The kinetic parameters *KM* and Vmax were calculated using Hans-Volf plots and an ANOVA calculator.

#### Activity assay of ecPGA to hydrolyze penG

The ecPGA activity assay using the harvested cells was performed according to Sobotková et al.^66^

#### VAF, YAF, and MSF mutants’ capability to hydrolase PenG

This was investigated spectrophotometrically in hydrolytic reactions with whole cells harvested from the stationary growth phase in MCHGly medium with 1 mM IPTG. The washed cells with phosphate buffer were used in the reaction with penicillin G (2% final concentration) at 37 °C, according to Balasingham et al.,^72^. The activity of one unit (U) is defined as the amount of the enzyme producing 1 μmol of 6-aminopenicillanic acid/min. The specific activity (SA) is expressed in U per g of cell dry weight.

#### Kinetically controlled synthesis of ampicillin and amoxicillin

Purified VAF, YAF, and MSF mutant variants were used in the kinetically controlled synthesis of semi-synthetic beta-lactam antibiotics ampicillin and amoxicillin. The syntheses of antibiotics were performed in 0.05 M potassium phosphate buffer adjusted to pH 7.0 and tempered to 25 °C. The reaction solution contained a 15 mM activated acyl donor (AD, D-phenylglycine amide) and 25 mM nucleophile (N, 6-APA or 7-ADCA). The addition of purified VAF, YAF, and MSF mutant variants started the synthesis. The performance of the synthetic reaction has been monitored using HPLC apparatus and Tessek SGX C18 HPLC column (250 x 4, 5 μm particle size, Tessek, Czech Republic). The mobile phase (0.01 M sodium phosphate buffer and methanol) differed for substrates: pH 3.0 and 10 % MetOH for amoxicillin and pH 5.6 and 30 % MetOH for ampicillin. The retention times of involved reactants and products are listed in Table S14.

#### His-tagged PGA variants production, purification, and melting analysis

*E. coli* BL21(DE3) cultures, transformed with their respective plasmids, were cultivated in 200 mL of 2YT media enriched with trace elements (TE) at 30 °C on an orbital incubator (250 rpm). Once the optical density (OD600) reached 0.6, heterologous protein expression was induced by adding 0.5 mM IPTG, then lowering the culture temperature to 16 °C, and incubation continued overnight. Subsequently, the cells were pelleted by centrifugation (8 000 x g, 5 min, 4 °C), resuspended in Tris buffer (50 mM Tris–HCl, pH 8.0, 100 mM NaCl), and sonicated (using a Sonicator 3000, Misonix, at 15-20 W). The sonicated lysate was treated with 5 U/mL of Benzonase® Nuclease and centrifuged (40 000 x g, 30 min, 4°C). The supernatant was loaded onto a gravity flow Ni-NTA column (2 ml, Econo-Pac® Chromatography Columns) using a Tris buffer. The column was washed with the same buffer containing 20 mM imidazole and eluted with an elution buffer (50 mM Tris–HCl, 100 mM NaCl, 250 mM imidazole, pH 8.0). The eluted fractions were subjected to analysis via SDS-PAGE, and the fractions with the highest concentrations of mutant PGA proteins were concentrated into PBS using Amicon Ultra-4 Centrifugal Filters with a 10 kDa molecular weight cutoff (Merck Millipore). This process resulted in a protein with a purity exceeding 70 % and a concentration of approximately 1 mg/mL. Protein melting curves were obtained by differential scanning fluorimetry of changes in tryptophan fluorescence at 330 and 350 nm over a temperature range of 20 °C to 95 °C, with increments of 1 °C per minute using a Prometheus NT.48 instrument (NanoTemper Technologies).

## Data Availability

The simulation and biochemical data generated in this study are available as *Supplementary Information Figures and Tables* comprising detailed selection criteria and visualization of residues selected for mutagenesis; properties of double- and triple-point mutants; characterization of docked poses; per-complex reactive stabilization scores with leave-one-out and outlier filtering and corresponding statistical significance as compared to wild-type enzyme for complexes with AHLs and PenG; thermostability of designed VAF, YAF and MSF variants; ampicillin and amoxicillin synthesis for ecPGA and VAF, YAF and MSF variants; correlation between experimental results and computational predictions; multiplots related to all replicates of MD simulations for protein-PenG complexes including RMSD of the protein backbone atoms, RMSD of the substrate and reactive stabilization information; RMSD time evolution of free enzyme simulations; in-depth characterization of PCA states; time-evolution of enzymes acyl-binding site openings; profiles of wild-type ecPGA and paPvdQ enzymes’ binding pockets; distribution of distances between center of mass for mutated residues; distribution of distances between center of mass for gating residues and mutated position 177β; crucial average distances representing components of reactive stabilization; frequency of the 3-oxo hydrogen bonding; graphical representation of reoccurring reactive stabilization after hydrogen bonding event of catalytic machinery with 3-oxo group; list of primers; retention times of substrates from HPLC experiments; *Supplementary Information File 2* with multiplots related to all replicates of MD simulations for protein-AHL complexes including RMSD of the protein backbone atoms, RMSD of the substrate and reactive stabilization information; *Supplementary Information File 3* with ecPGA variants activity as a function of substrate concentration for conversion of AHLs; *Supplementary Information File 4* with multiplots related to all replicates of MD simulations for protein complexes with 3-oxo-AHLs including RMSD of the protein backbone atoms, RMSD of substrate, reactive stabilization information and hydrogen bond between substrate 3-oxo group and oxyanion hole stabilizing residue. The data underlying this study are available from Zenodo repository at https://doi.org/10.5281/zenodo.7980949.

## Acknowledgements

This work was supported by the National Science Centre, Poland (Grant Numbers 2021/41/N/NZ2/01365 and 2017/26/E/NZ1/00548), Institutional Research project RVO61388971 from the Institute of Microbiology of the CAS, the Czech Foundation grant 21-17044S, and by the Ministry of Education, Youth and Sports of the Czech Republic grant Talking microbes - understanding microbial interactions within One Health framework (CZ.02.01.01/00/22_008/0004597) and with the project of National Institute of Virology and Bacteriology, Programme EXCELES, funded by the European Union, Next Generation EU (LX22NPO5103). B.S. is a recipient of a scholarship provided by POWER project POWR.03.02.00-00-I022/16. The computations were performed at the Poznan Supercomputing and Networking Center.

## AUTHOR INFORMATION

These authors contributed equally: Michal Grulich, Bartlomiej Surpeta

## Contributions

M.G. designed and coordinated the experimental work, performed detailed characterization of all the ecPGA designed variants (YAF, VAF and MSF), and analyzed and interpreted data. B.S. designed ecPGA variants and performed all computational work, analyzed and interpreted data. H.M. and A.P. executed all of the bacterial flasks and fed-batch cultivation in a bioreactor to produce enzymes, H.M. additionally performed PenG hydrolysis assays with PGA variants. J.Z. designed the set of used primers for the ecPGA variants and determined their thermal stability with differential scanning fluorimetry. J.B. conceived and coordinated the project, designed computational analyses, analyzed and interpreted data.

The manuscript was written through the contributions of all authors. All authors have approved the final version of the manuscript.

