## Supplementary Information Figures and Tables for "Engineering dynamic gates in binding pocket of penicillin G acylase to selectively degrade bacterial signaling molecules"

‡ These authors contributed equally.

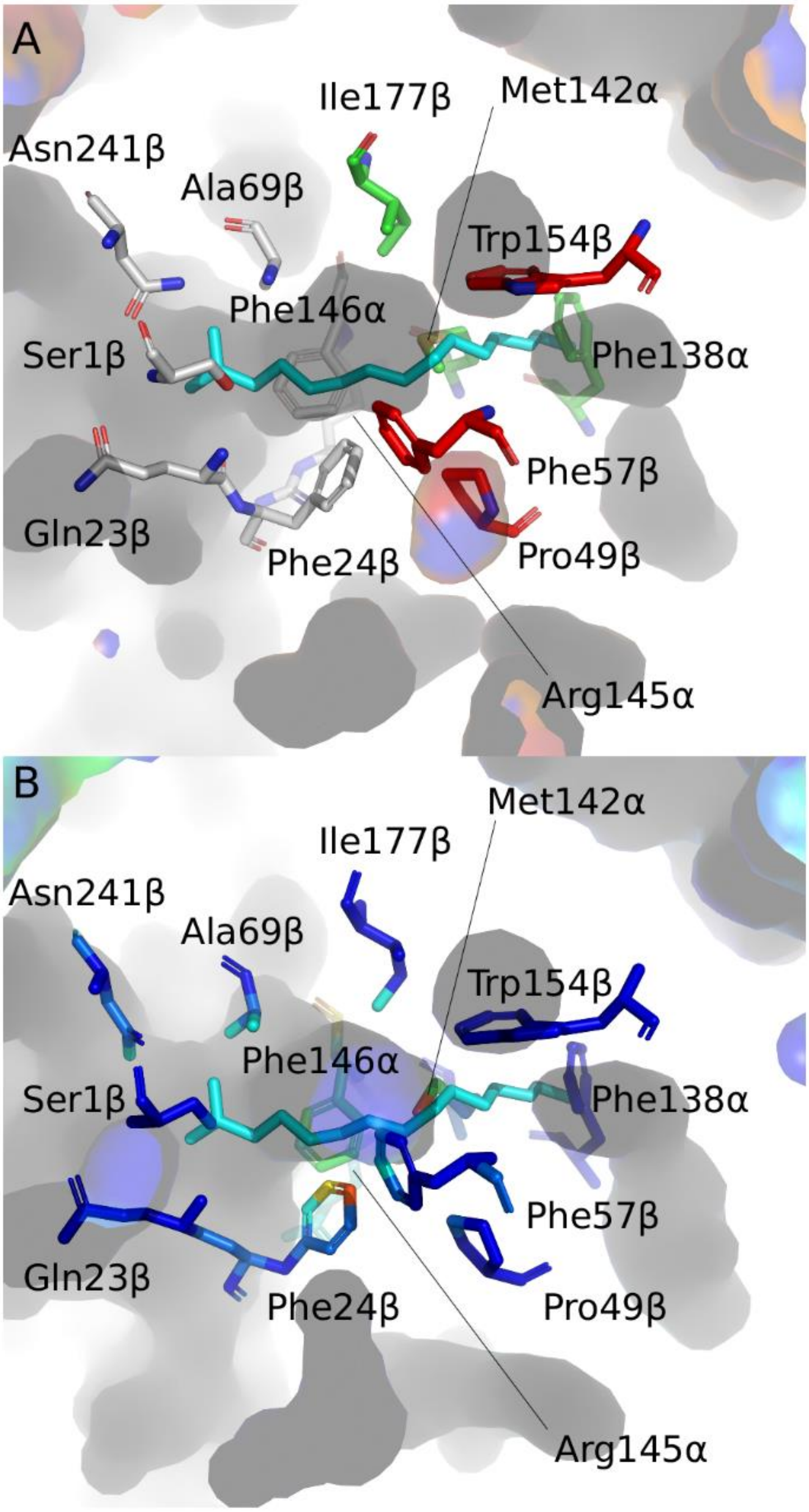


**Figure S1. Selection of residues for the mutagenesis.** A. General composition of the ecPGA binding pocket (grey sticks - functional or gating residues, red sticks - residues with low mutability score based on sequence consensus analysis performed by HotSpot Wizard 3.0 analysis, green sticks – residues selected for modification). B. Residues colored by the contribution in forming the acyl-binding pocket based on the free enzyme molecular dynamics of ecPGA wild-type protein, analyzed with mdpocket (color-coding - the warmer the more conserved – more frequently contributing to the pocket formation). paPvdQ transition stage analog is shown as cyan sticks superimposed to ecPGA structure.

**Table S1. Double-point mutants' properties.** First row indicate the filters applied for filtering applied to achieve best candidates for the next round. Row shaded in gray represents wild-type properties used to design the filters.

| Total_No_Frames | Avg_BR* | StDev | Max_BR* | Avg_Len* | StDev.1 | Avg_throug | StDev.2 | Mutation** | FoldX_diff*** | FoldX_std |
|---|---|---|---|---|---|---|---|---|---|---|
| >WT (2/3) | >WT | | >WT | >WT | | >WT | | | Sum < 4.0 | |
| 2 | 1.05 | 0.02 | 1.07 | 10.43 | 1.64 | 0.58 | 0.06 | wt,ecPGA | - | - |
| 30 | 1.69 | 0.1 | 1.83 | 10.39 | 2.17 | 0.76 | 0.04 | MA142G,IB177F | 3.21 | 0.08 |
| 30 | 1.36 | 0.13 | 1.57 | 11.21 | 2 | 0.73 | 0.04 | MA142A,IB177F | 2.44 | 0.06 |
| 8 | 0.97 | 0.03 | 1.02 | 13.41 | 0.95 | 0.37 | 0.03 | MA142L,IB177F | 0.47 | 0.41 |
| 4 | 1.03 | 0.09 | 1.14 | 13.53 | 1.23 | 0.44 | 0.1 | MA142F,IB177F | 4.32 | 4.31 |
| 9 | 0.96 | 0.03 | 1.01 | 13.27 | 1.69 | 0.45 | 0.12 | MA142W,IB177F | 9.84 | 1.16 |
| 2 | 0.95 | 0.03 | 0.98 | 5.33 | 0.02 | 0.76 | 0.01 | MA142K,IB177F | 3.71 | 0.47 |
| 7 | 0.98 | 0.1 | 1.2 | 13.64 | 0.58 | 0.56 | 0.05 | MA142Q,IB177F | 2 | 0.34 |
| 11 | 0.99 | 0.05 | 1.07 | 13.32 | 0.85 | 0.57 | 0.04 | MA142E,IB177F | 1.47 | 0.4 |
| 30 | 1.24 | 0.12 | 1.47 | 11.35 | 2.5 | 0.7 | 0.07 | MA142S,IB177F | 2.37 | 0.09 |
| 30 | 1.56 | 0.08 | 1.71 | 11.13 | 2.01 | 0.74 | 0.04 | MA142P,IB177F | 7.83 | 0.6 |
| 16 | 1.03 | 0.1 | 1.16 | 14.7 | 8.82 | 0.61 | 0.18 | MA142V,IB177F | 2.59 | 0.48 |
| 10 | 1 | 0.08 | 1.17 | 13.13 | 0.56 | 0.52 | 0.13 | MA142I,IB177F | 2.41 | 0.51 |
| 9 | 0.92 | 0.02 | 0.95 | 13.14 | 0.84 | 0.37 | 0.05 | MA142C,IB177F | 2.22 | 0.06 |
| - | - | - | - | - | - | - | - | MA142Y,IB177F | 7.91 | 4.84 |
| 4 | 0.94 | 0.02 | 0.96 | 12.59 | 0.75 | 0.41 | 0.05 | MA142H,IB177F | 5.33 | 2.08 |
| 1 | 0.93 | 0 | 0.93 | 13.4 | 0 | 0.36 | 0 | MA142R,IB177F | 3.73 | 0.7 |
| - | - | - | - | - | - | - | - | MA142N,IB177F | 2.13 | 0.56 |
| 1 | 0.96 | 0 | 0.96 | 12.3 | 0 | 0.43 | 0 | MA142D,IB177F | 2.62 | 0.61 |
| 21 | 1.1 | 0.13 | 1.27 | 16.53 | 12.69 | 0.6 | 0.2 | MA142T,IB177F | 2.53 | 0.22 |
| 30 | 1.67 | 0.11 | 1.82 | 10.63 | 2.19 | 0.76 | 0.03 | IB177F,MA142G | 3.2 | 0.09 |
| 30 | 1.41 | 0.1 | 1.6 | 10.99 | 1.81 | 0.74 | 0.03 | IB177F,MA142A | 2.39 | 0.07 |
| 1 | 0.95 | 0 | 0.95 | 13.35 | 0 | 0.35 | 0 | IB177F,MA142L | -0.35 | 0.39 |
| 2 | 0.92 | 0 | 0.92 | 12.32 | 1.72 | 0.52 | 0.02 | IB177F,MA142F | 3.37 | 1.77 |
| 2 | 0.96 | 0.04 | 1 | 11.3 | 1.21 | 0.54 | 0.15 | IB177F,MA142W | 9.22 | 1.28 |
| - | - | - | - | - | - | - | - | IB177F,MA142K | 3.71 | 0.47 |
| 6 | 1.01 | 0.06 | 1.12 | 13.33 | 0.89 | 0.56 | 0.07 | IB177F,MA142Q | 2.24 | 0.52 |
| 8 | 1 | 0.06 | 1.11 | 12.84 | 0.36 | 0.59 | 0.04 | IB177F,MA142E | 1.43 | 0.57 |
| 30 | 1.23 | 0.13 | 1.49 | 11.42 | 2.75 | 0.7 | 0.08 | IB177F,MA142S | 2.38 | 0.08 |
| 30 | 1.58 | 0.09 | 1.77 | 10.95 | 2.02 | 0.74 | 0.04 | IB177F,MA142P | 7.41 | 0.52 |
| 19 | 1.05 | 0.1 | 1.22 | 14.45 | 8.29 | 0.64 | 0.16 | IB177F,MA142V | 2.84 | 0.48 |
| 13 | 1.07 | 0.08 | 1.21 | 12.43 | 2.44 | 0.63 | 0.07 | IB177F,MA142I | 2.73 | 0.53 |
| 3 | 0.93 | 0.01 | 0.94 | 13.08 | 0.43 | 0.35 | 0.01 | IB177F,MA142C | 2.22 | 0.1 |
| 2 | 0.94 | 0.02 | 0.96 | 13.28 | 0.7 | 0.35 | 0.01 | IB177F,MA142Y | 5.23 | 2.91 |
| - | - | - | - | - | - | - | - | IB177F,MA142H | 6.62 | 2.38 |
| - | - | - | - | - | - | - | - | IB177F,MA142R | 3.72 | 0.6 |
| - | - | - | - | - | - | - | - | IB177F,MA142N | 2.28 | 0.54 |
| - | - | - | - | - | - | - | - | IB177F,MA142D | 2.86 | 0.63 |
| 17 | 1.12 | 0.12 | 1.27 | 11.49 | 2.7 | 0.7 | 0.05 | IB177F,MA142T | 2.46 | 0.23 |

* in Å;

** MA142G,IB177F corresponds to mutation Met142αGly & Ile177βPhe, remaining mutations have analogous naming convention;

*** in kcal/mol

Table S2. Selected double-point mutants for the next stage of engineering.

| Total_No_Frames | Avg_BR* | StDev | Max_BR* | Avg_Len* | StDev.1 | Avg_throug | StDev.3 | Mutation** | FoldX_diff*** | FoldX_std |
|---|---|---|---|---|---|---|---|---|---|---|
| 30 | 1.36 | 0.13 | 1.57 | 11.21 | 2.00 | 0.73 | 0.04 | MA142A,IB177F | 2.44 | 0.06 |
| 30 | 1.24 | 0.12 | 1.47 | 11.35 | 2.50 | 0.70 | 0.07 | MA142S,IB177F | 2.37 | 0.09 |
| 30 | 1.41 | 0.10 | 1.60 | 10.99 | 1.81 | 0.74 | 0.03 | IB177F,MA142A | 2.39 | 0.07 |
| 30 | 1.23 | 0.13 | 1.49 | 11.42 | 2.75 | 0.70 | 0.08 | IB177F,MA142S | 2.38 | 0.08 |

* in Å;
** MA142G,IB177F corresponds to mutation Met142αGly & Ile177βPhe, remaining mutations have analogous naming convention;
*** in kcal/mol

**Table S3. Triple-point mutants' properties.** First row indicate the filters applied for filtering applied to achieve best candidates for the extensive molecular modelling. Row shaded in gray represents wild-type properties used to design the filters.

| Total_No_Frames | Avg_BR* | StDev | Max_BR* | Avg_Len* | StDev.1 | Avg_throug | StDev.3 | Mutation** | FoldX_diff*** | FoldX_std |
|---|---|---|---|---|---|---|---|---|---|---|
| >= 95 % | >= 1.2 | | >=1.5 | >= 12 | | | | | sum < 6.0 | |
| 2 | 1.05 | 0.02 | 1.07 | 10.43 | 1.64 | 0.58 | 0.06 | ecPGA,wt, | - | - |
| 60 | 1.32 | 0.12 | 1.48 | 16.90 | 3.64 | 0.62 | 0.07 | FA138G,MA142A,IB177F | 8.08 | 0.11 |
| 60 | 1.36 | 0.11 | 1.56 | 16.50 | 3.46 | 0.62 | 0.07 | FA138A,MA142A,IB177F | 6.34 | 0.12 |
| 60 | 1.30 | 0.18 | 1.58 | 13.54 | 2.19 | 0.66 | 0.07 | FA138L,MA142A,IB177F | 4.07 | 0.17 |
| 60 | 1.31 | 0.22 | 1.64 | 13.20 | 2.78 | 0.66 | 0.09 | FA138M,MA142A,IB177F | 3.65 | 0.12 |
| 56 | 1.34 | 0.12 | 1.57 | 11.56 | 2.31 | 0.72 | 0.06 | FA138W,MA142A,IB177F | 4.88 | 0.13 |
| 60 | 1.39 | 0.13 | 1.64 | 13.11 | 3.20 | 0.68 | 0.08 | FA138K,MA142A,IB177F | 6.35 | 0.17 |
| 60 | 1.37 | 0.14 | 1.61 | 13.22 | 2.90 | 0.68 | 0.07 | FA138Q,MA142A,IB177F | 6.41 | 0.24 |
| 60 | 1.39 | 0.25 | 1.69 | 11.91 | 3.00 | 0.69 | 0.11 | FA138E,MA142A,IB177F | 7.39 | 0.10 |
| 60 | 1.39 | 0.13 | 1.59 | 15.37 | 3.35 | 0.63 | 0.07 | FA138S,MA142A,IB177F | 7.48 | 0.20 |
| 60 | 1.40 | 0.13 | 1.58 | 15.73 | 3.13 | 0.63 | 0.06 | FA138P,MA142A,IB177F | 6.50 | 0.10 |
| 60 | 1.34 | 0.09 | 1.57 | 15.13 | 2.11 | 0.62 | 0.05 | FA138V,MA142A,IB177F | 5.30 | 0.14 |
| 60 | 1.35 | 0.13 | 1.59 | 14.25 | 2.47 | 0.64 | 0.07 | FA138I,MA142A,IB177F | 5.42 | 0.87 |
| 60 | 1.33 | 0.10 | 1.54 | 14.89 | 2.17 | 0.62 | 0.05 | FA138C,MA142A,IB177F | 6.87 | 0.16 |
| 57 | 1.36 | 0.14 | 1.59 | 12.06 | 5.34 | 0.72 | 0.11 | FA138Y,MA142A,IB177F | 3.55 | 0.09 |
| 60 | 1.47 | 0.13 | 1.65 | 11.56 | 3.85 | 0.73 | 0.08 | FA138H,MA142A,IB177F | 4.65 | 0.10 |
| 60 | 1.51 | 0.18 | 1.81 | 10.86 | 2.43 | 0.74 | 0.06 | FA138R,MA142A,IB177F | 7.99 | 0.57 |
| 60 | 1.33 | 0.14 | 1.57 | 13.17 | 3.14 | 0.67 | 0.09 | FA138N,MA142A,IB177F | 6.17 | 0.11 |
| 60 | 1.34 | 0.16 | 1.63 | 13.36 | 2.72 | 0.65 | 0.08 | FA138D,MA142A,IB177F | 7.96 | 0.15 |
| 60 | 1.41 | 0.11 | 1.60 | 15.98 | 2.49 | 0.62 | 0.05 | FA138T,MA142A,IB177F | 6.86 | 0.24 |
| 60 | 1.30 | 0.11 | 1.44 | 15.88 | 5.02 | 0.63 | 0.09 | FA138G,MA142S,IB177F | 7.90 | 0.08 |
| 60 | 1.32 | 0.10 | 1.45 | 15.70 | 4.02 | 0.63 | 0.08 | FA138A,MA142S,IB177F | 6.27 | 0.09 |
| 60 | 1.21 | 0.19 | 1.54 | 14.58 | 2.57 | 0.60 | 0.09 | FA138L,MA142S,IB177F | 3.93 | 0.11 |
| 60 | 1.26 | 0.23 | 1.57 | 14.06 | 3.90 | 0.62 | 0.13 | FA138M,MA142S,IB177F | 3.57 | 0.13 |
| 52 | 1.20 | 0.14 | 1.43 | 11.65 | 3.11 | 0.69 | 0.09 | FA138W,MA142S,IB177F | 4.91 | 0.17 |
| 60 | 1.29 | 0.18 | 1.59 | 13.11 | 3.80 | 0.66 | 0.11 | FA138K,MA142S,IB177F | 6.02 | 0.25 |
| 60 | 1.36 | 0.13 | 1.60 | 13.64 | 3.61 | 0.66 | 0.09 | FA138Q,MA142S,IB177F | 6.10 | 0.23 |
| 60 | 1.18 | 0.23 | 1.62 | 14.27 | 4.35 | 0.59 | 0.14 | FA138E,MA142S,IB177F | 7.00 | 0.28 |
| 60 | 1.35 | 0.09 | 1.47 | 15.10 | 3.82 | 0.63 | 0.09 | FA138S,MA142S,IB177F | 7.32 | 0.19 |
| 60 | 1.37 | 0.10 | 1.55 | 15.30 | 4.28 | 0.63 | 0.09 | FA138P,MA142S,IB177F | 6.33 | 0.18 |
| 60 | 1.36 | 0.12 | 1.55 | 14.67 | 2.99 | 0.62 | 0.08 | FA138V,MA142S,IB177F | 5.12 | 0.15 |
| 60 | 1.37 | 0.18 | 1.61 | 14.12 | 3.21 | 0.64 | 0.09 | FA138I,MA142S,IB177F | 5.46 | 0.89 |
| 60 | 1.35 | 0.11 | 1.46 | 14.54 | 3.08 | 0.63 | 0.09 | FA138C,MA142S,IB177F | 6.66 | 0.14 |
| 60 | 1.22 | 0.13 | 1.46 | 11.37 | 2.74 | 0.70 | 0.08 | FA138Y,MA142S,IB177F | 3.51 | 0.09 |
| 60 | 1.19 | 0.17 | 1.53 | 13.57 | 6.20 | 0.64 | 0.15 | FA138H,MA142S,IB177F | 4.86 | 0.15 |
| 58 | 1.28 | 0.18 | 1.64 | 11.82 | 3.25 | 0.69 | 0.10 | FA138R,MA142S,IB177F | 7.69 | 0.76 |
| 60 | 1.28 | 0.14 | 1.51 | 14.22 | 3.55 | 0.63 | 0.10 | FA138N,MA142S,IB177F | 6.09 | 0.10 |
| 60 | 1.32 | 0.13 | 1.54 | 14.30 | 3.85 | 0.62 | 0.11 | FA138D,MA142S,IB177F | 7.87 | 0.08 |
| 60 | 1.37 | 0.10 | 1.54 | 15.39 | 3.34 | 0.63 | 0.07 | FA138T,MA142S,IB177F | 6.81 | 0.34 |

* in Å;

** FA138G,MA142G,IB177F corresponds to mutation Phe138αGly & Met142αGly & Ile177βPhe, remaining mutations have analogous naming convention;

*** in kcal/mol

**Table S4. Selected triple-point mutants for the molecular docking experiments.** Row shaded in gray represent wild-type properties. Rows shaded in green and red represent the mutants with highest destabilization effect. Only one of these was promoted for further stage – mutant VAF (green) – based on its better effect on the pocket depth. Mutant VSF (red) was discarded.

| Total_No_Frames | Avg_BR* | StDev | Max_BR* | Avg_Len* | StDev.1 | Avg_throug | StDev.3 | Mutation** | FoldX_diff*** | FoldX_std |
|---|---|---|---|---|---|---|---|---|---|---|
| **2** | **1.05** | **0.02** | **1.07** | **10.43** | **1.64** | **0.58** | **0.06** | **ecPGA, wt** | | |
| 60 | 1.30 | 0.18 | 1.58 | 13.54 | 2.19 | 0.66 | 0.07 | FA138L,MA142A,IB177F (LAF) | 4.07 | 0.17 |
| 60 | 1.31 | 0.22 | 1.64 | 13.20 | 2.78 | 0.66 | 0.09 | FA138M,MA142A,IB177F (MAF) | 3.65 | 0.12 |
| 60 | 1.34 | 0.09 | 1.57 | 15.13 | 2.11 | 0.62 | 0.05 | FA138V,MA142A,IB177F, (VAF) | 5.30 | 0.14 |
| 57 | 1.36 | 0.14 | 1.59 | 12.06 | 5.34 | 0.72 | 0.11 | FA138Y,MA142A,IB177F (YAF) | 3.55 | 0.09 |
| 60 | 1.21 | 0.19 | 1.54 | 14.58 | 2.57 | 0.60 | 0.09 | FA138L,MA142S,IB177F (LSF) | 3.93 | 0.11 |
| 60 | 1.26 | 0.23 | 1.57 | 14.06 | 3.90 | 0.62 | 0.13 | FA138M,MA142S,IB177F (MSF) | 3.57 | 0.13 |
| 60 | 1.36 | 0.12 | 1.55 | 14.67 | 2.99 | 0.62 | 0.08 | FA138V,MA142S,IB177F (VSF) | 5.12 | 0.15 |

* in Å;
** FA138L,MA142A,IB177F (LAF) corresponds to mutation Phe138αLeu & Met142αAla & Ile177βPhe, remaining mutations have analogous naming convention;
*** in kcal/mol

Table S5. Representative complexes selected from molecular docking experiments for molecular dynamics simulations stage.

| | | | Distances [Å] | | | |
|---|---|---|---|---|---|---|
| Variant | Substrate | Energy [kcal/mol] | Ser1β-O - HSL-C | Ala69β-NH - HSL-O | Asn241β-N δH2 – HSL-O | Gln23β-O - HSL-NH |
| LAF | C06 | -5.50 | 2.70 | 2.25 | 2.09 | 2.27 |
| | C06-3O | -5.19 | 2.72 | 2.24 | 2.18 | 2.24 |
| | C08 | -6.29 | 2.73 | 2.28 | 2.08 | 2.31 |
| | C08-30 | -5.97 | 2.72 | 2.22 | 2.17 | 2.23 |
| | C10 | -7.28 | 2.73 | 2.25 | 2.10 | 2.22 |
| | C12-3O | -7.94 | 2.71 | 2.22 | 2.16 | 2.21 |
| MAF | C06 | -5.67 | 2.69 | 2.35 | 2.12 | 2.23 |
| | C06-3O | -5.40 | 2.70 | 2.33 | 2.17 | 2.20 |
| | C08 | -6.43 | 2.70 | 2.36 | 2.13 | 2.21 |
| | C08-3O | -6.10 | 2.71 | 2.32 | 2.17 | 2.19 |
| | C10 | -7.47 | 2.71 | 2.27 | 2.05 | 2.31 |
| | C12-3O | -7.75 | 2.69 | 2.32 | 1.93 | 2.66 |
| VAF | C06 | -5.31 | 2.75 | 2.21 | 2.45 | 2.24 |
| | C06-3O | -5.02 | 2.75 | 2.27 | 2.51 | 2.20 |
| | C08 | -6.12 | 2.73 | 2.14 | 2.68 | 2.17 |
| | C08-3O | -5.83 | 2.75 | 2.33 | 2.53 | 2.18 |
| | C10 | -7.12 | 2.77 | 2.14 | 2.83 | 2.17 |
| | C12-3O | -7.50 | 2.72 | 2.13 | 2.69 | 2.20 |
| YAF | C06 | -5.52 | 2.71 | 2.25 | 2.10 | 2.25 |
| | C06-3O | -5.23 | 2.72 | 2.32 | 2.19 | 2.21 |
| | C08 | -6.21 | 2.71 | 2.22 | 2.04 | 2.29 |
| | C08-3O | -5.96 | 2.71 | 2.14 | 2.27 | 2.31 |
| | C10 | -7.29 | 2.73 | 2.14 | 2.39 | 2.21 |
| | C12-3O | -7.77 | 2.67 | 2.24 | 2.08 | 2.21 |
| LSF | C06 | -5.40 | 2.75 | 2.23 | 2.31 | 2.23 |
| | C06-3O | -5.12 | 2.74 | 2.22 | 2.27 | 2.30 |
| | C08 | -6.16 | 2.73 | 2.23 | 2.24 | 2.22 |
| | C08-3O | -5.79 | 2.74 | 2.15 | 2.35 | 2.35 |
| | C10 | -7.16 | 2.72 | 2.14 | 2.52 | 2.15 |
| | C12-3O | -7.53 | 2.71 | 2.14 | 2.44 | 2.22 |
| MSF | C06 | -5.46 | 2.75 | 2.22 | 2.21 | 2.24 |
| | C06-3O | -5.18 | 2.75 | 2.26 | 2.25 | 2.23 |
| | C08 | -6.23 | 2.75 | 2.23 | 2.09 | 2.34 |
| | C08-3O | -5.95 | 2.74 | 2.22 | 2.21 | 2.24 |
| | C10 | -7.24 | 2.80 | 2.26 | 1.98 | 2.59 |
| | C12-3O | -7.62 | 2.77 | 2.37 | 2.26 | 2.20 |
| ecPGA-wt | C06 | -5.31 | 2.74 | 2.20 | 2.19 | 2.27 |
| | C06-3O | -5.04 | 2.74 | 2.15 | 2.39 | 2.18 |
| | C08 | -6.15 | 2.76 | 2.28 | 2.11 | 2.40 |
| | C08-3O | -5.93 | 2.71 | 2.21 | 2.12 | 2.23 |
| | C10 | -6.77 | 2.72 | 2.30 | 2.19 | 2.25 |
| | C12-3O | -5.67 | 2.81 | 2.31 | 2.48 | 2.18 |

Table S6. Reactive stabilization score statistics.

| protein | substrate | RSS | RSS filtered outliers | RSS filtered outliers leave-one-out | RSS filtered outliers leave-one-out std | RSS filtered outiers leave-one-out n samples | mutant vs wt ratio | mutant to wt difference statistics | mutant to wt difference pvalue |
|---|---|---|---|---|---|---|---|---|---|
| LAF | C06 | 1.0535 | 0.7807 | 0.7791 | 0.0586 | 31 | 0.94 | -2.39 | 2.01E-02 |
| | C06-3O | 0.5011 | 0.3563 | 0.3566 | 0.0271 | 32 | 3.10 | 47.03 | 3.28E-50 |
| | C08 | 0.4703 | 0.2755 | 0.2799 | 0.0299 | 31 | 18.54 | 49.98 | 3.32E-51 |
| | C08-3O | 0.8861 | 0.6480 | 0.6492 | 0.0598 | 30 | 90.59 | 58.79 | 2.15E-53 |
| | C10 | 0.1911 | 0.1080 | 0.1094 | 0.0116 | 33 | 112.99 | 50.36 | 8.54E-51 |
| | C12-3O | 0.2695 | 0.1995 | 0.1972 | 0.0199 | 32 | 125.69 | 57.38 | 9.37E-57 |
| MAF | C06 | 1.6582 | 0.8460 | 0.8447 | 0.0755 | 32 | 1.02 | 0.90 | 3.70E-01 |
| | C06-3O | 1.1452 | 0.7542 | 0.7498 | 0.0495 | 33 | 6.52 | 70.99 | 7.73E-62 |
| | C08 | 1.2735 | 0.7913 | 0.8001 | 0.0660 | 33 | 53.01 | 67.28 | 2.17E-60 |
| | C08-3O | 1.7898 | 1.0431 | 1.0475 | 0.0863 | 29 | 146.17 | 66.07 | 1.47E-55 |
| | C10 | 0.1783 | 0.0734 | 0.0722 | 0.0090 | 34 | 74.57 | 42.55 | 4.60E-47 |
| | C12-3O | 0.0206 | 0.0086 | 0.0088 | 0.0011 | 31 | 5.60 | 35.89 | 1.18E-43 |
| VAF | C06 | 1.8418 | 1.0467 | 1.0400 | 0.1146 | 31 | 1.26 | 8.14 | 2.87E-11 |
| | C06-3O | 0.8435 | 0.8435 | 0.8480 | 0.0642 | 32 | 7.37 | 63.70 | 3.23E-58 |
| | C08 | 1.0564 | 0.3613 | 0.3586 | 0.0346 | 32 | 23.76 | 56.13 | 7.28E-55 |
| | C08-3O | 2.0925 | 1.5978 | 1.6065 | 0.1225 | 31 | 224.19 | 71.49 | 5.13E-59 |
| | C10 | 0.0021 | 0.0007 | 0.0007 | 0.0001 | 33 | 0.77 | -5.30 | 1.73E-06 |
| | C12-3O | 0.0386 | 0.0240 | 0.0241 | 0.0034 | 30 | 15.37 | 38.46 | 5.66E-45 |
| YAF | C06 | 1.0804 | 0.4971 | 0.5054 | 0.0672 | 29 | 0.61 | -15.36 | 4.15E-22 |
| | C06-3O | 1.4339 | 0.8582 | 0.8555 | 0.0567 | 32 | 7.44 | 72.62 | 1.06E-61 |
| | C08 | 0.1879 | 0.0917 | 0.0920 | 0.0104 | 31 | 6.10 | 41.30 | 2.70E-46 |
| | C08-3O | 1.2570 | 0.9182 | 0.9247 | 0.0711 | 32 | 129.04 | 70.64 | 1.82E-59 |
| | C10 | 0.2061 | 0.0229 | 0.0232 | 0.0028 | 33 | 23.99 | 42.83 | 1.09E-46 |
| | C12-3O | 0.3933 | 0.1876 | 0.1919 | 0.0280 | 31 | 122.33 | 39.67 | 2.80E-46 |
| LSF | C06 | 0.6542 | 0.2861 | 0.2841 | 0.0336 | 30 | 0.34 | -30.49 | 8.25E-38 |
| | C06-3O | 1.7213 | 1.3148 | 1.3083 | 0.1115 | 30 | 11.38 | 60.26 | 2.20E-55 |
| | C08 | 0.6190 | 0.1688 | 0.1716 | 0.0192 | 29 | 11.37 | 45.94 | 6.91E-48 |
| | C08-3O | 2.9523 | 2.9523 | 2.9321 | 0.2053 | 31 | 409.16 | 78.01 | 3.13E-61 |
| | C10 | 0.0371 | 0.0082 | 0.0084 | 0.0013 | 29 | 8.65 | 31.30 | 3.61E-37 |
| | C12-3O | 0.6126 | 0.2236 | 0.2257 | 0.0216 | 30 | 143.84 | 60.56 | 7.09E-57 |
| MSF | C06 | 2.2341 | 0.9675 | 0.9661 | 0.1262 | 28 | 1.17 | 4.92 | 7.60E-06 |
| | C06-3O | 1.0668 | 0.8081 | 0.8097 | 0.0628 | 29 | 7.04 | 61.70 | 2.73E-55 |
| | C08 | 0.5328 | 0.1759 | 0.1788 | 0.0205 | 29 | 11.85 | 45.06 | 2.09E-47 |
| | C08-3O | 1.1413 | 0.6921 | 0.6893 | 0.0571 | 29 | 96.19 | 65.40 | 2.60E-55 |
| | C10 | 0.0288 | 0.0182 | 0.0184 | 0.0023 | 29 | 18.99 | 40.81 | 2.37E-43 |
| | C12-3O | 0.7332 | 0.5059 | 0.5047 | 0.0475 | 29 | 321.68 | 61.85 | 9.59E-57 |
| ecPGA | C06 | 1.5040 | 0.8216 | 0.8257 | 0.0915 | 31 | 1.00 | 0.00 | 1.00E+00 |
| | C06-3O | 0.1527 | 0.1155 | 0.1150 | 0.0106 | 32 | 1.00 | 0.00 | 1.00E+00 |
| | C08 | 0.0407 | 0.0151 | 0.0151 | 0.0018 | 32 | 1.00 | 0.00 | 1.00E+00 |
| | C08-3O | 0.0214 | 0.0069 | 0.0072 | 0.0012 | 30 | 1.00 | 0.00 | 1.00E+00 |
| | C10 | 0.0027 | 0.0010 | 0.0010 | 0.0002 | 29 | 1.00 | 0.00 | 1.00E+00 |
| | C12-3O | 0.0031 | 0.0016 | 0.0016 | 0.0003 | 34 | 1.00 | 0.00 | 1.00E+00 |

Statistical test calculated using two-sided ttest, as implemented in python scipy library (ttest_ind_from_stats).

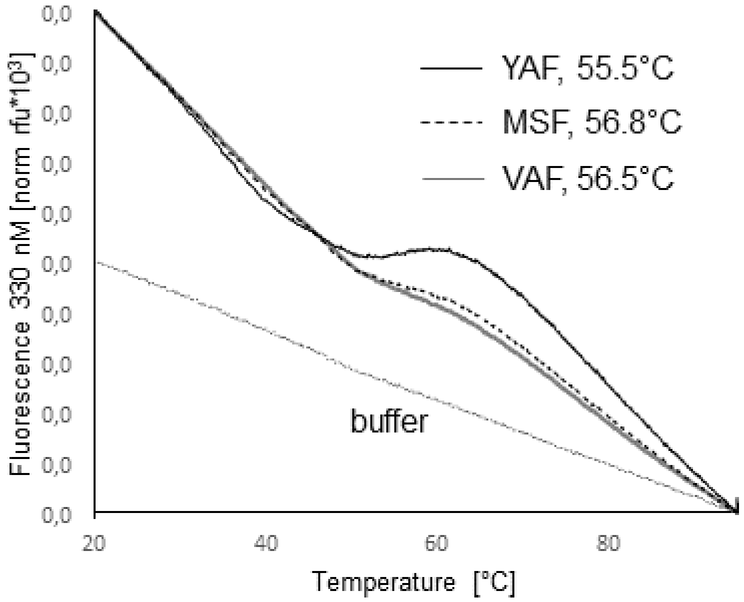


Figure S2. Thermostability of designed ecPGA variants VAF, YAF and MSF.

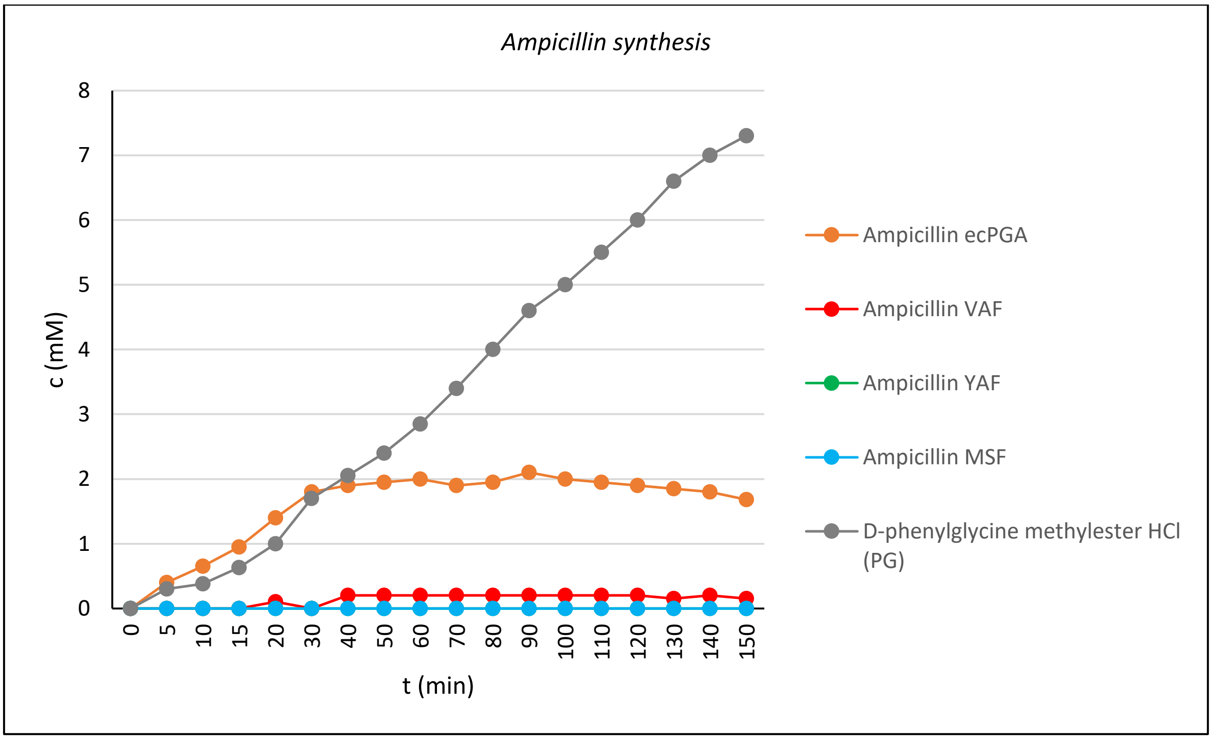


Figure S3. Ampicillin synthesis by ecPGA wild-type and VAF, YAF and MSF variants.

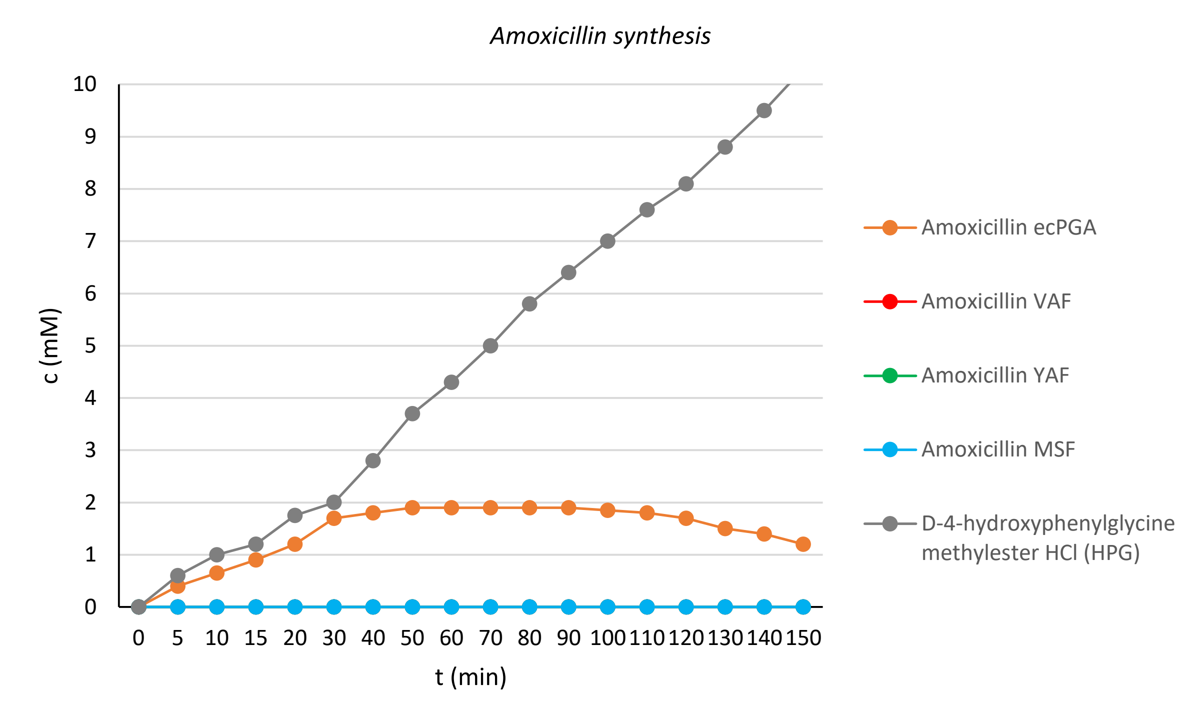


Figure S4. Amoxicillin synthesis by ecPGA wild-type and VAF, YAF and MSF variants.

Table S7. S/H ratio a $P_{smax}$ parameters for kinetically controlled syntheses of semi-synthetic β-lactam antibiotics ampicillin and amoxicillin catalyzed by wild-type ecPGA variants VAF, YAF and MSF.

| Activated acyl-donor (AD) | β-lactam nucleophile (N) | Antibiotics | $P_{Smax}$ (mM) | | | | S/H | | | |
|---|---|---|---|---|---|---|---|---|---|---|
| | | | ecPGA | VAF | YAF | MSF | ecPGA | VAF | YAF | MSF |
| D-PGA | 6-APA | Ampicillin | 1.9 | - | - | - | 0.8 | - | - | - |
| D-HPGA | 6-APA | Amoxicillin | 1.5 | - | - | - | 0.8 | - | - | - |

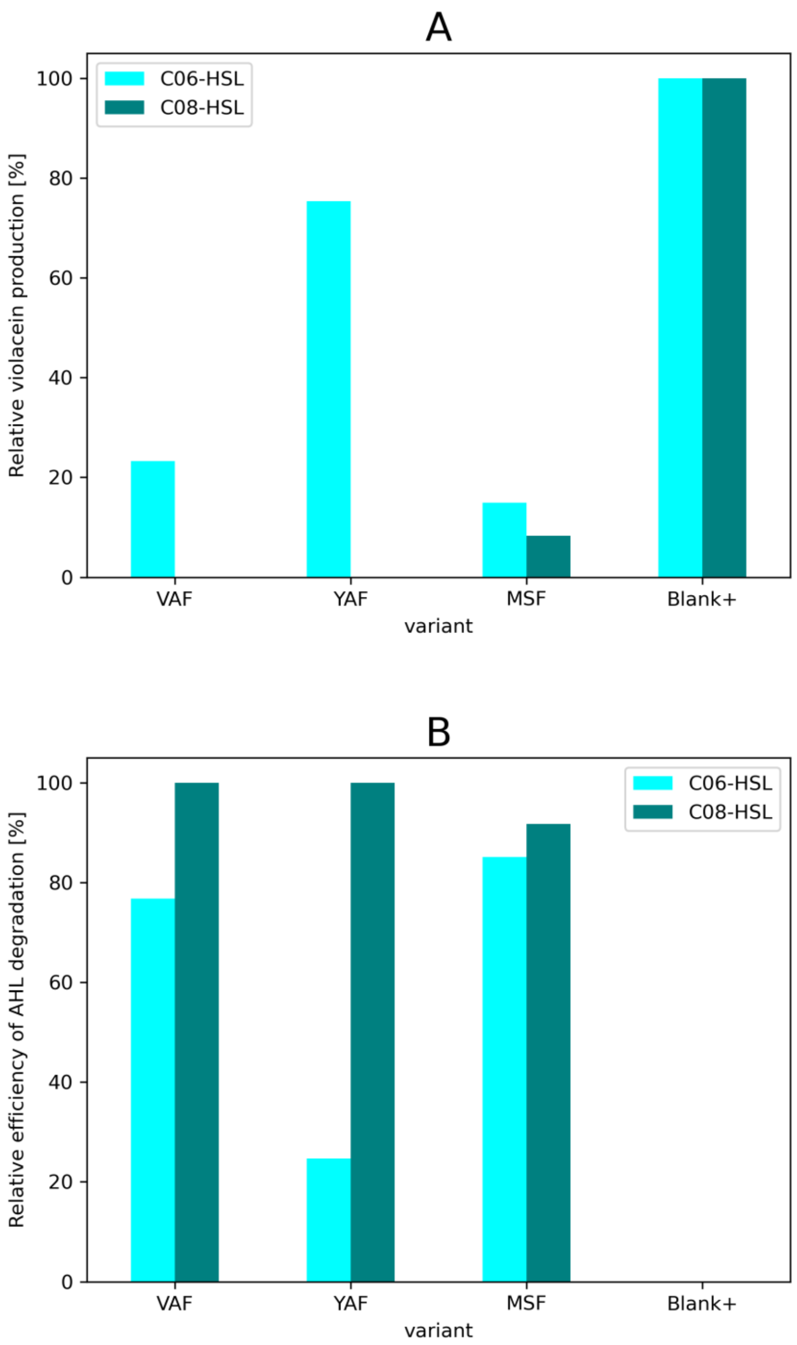


**Figure S5. Relative violacein production by *C. violaceum* after AHL degradation by crude cell lysates with ecPGA enzyme mutants (A) and relative efficiency of AHL degradation (B).** Blank+ represents production of violacein in the presence of AHL molecule.

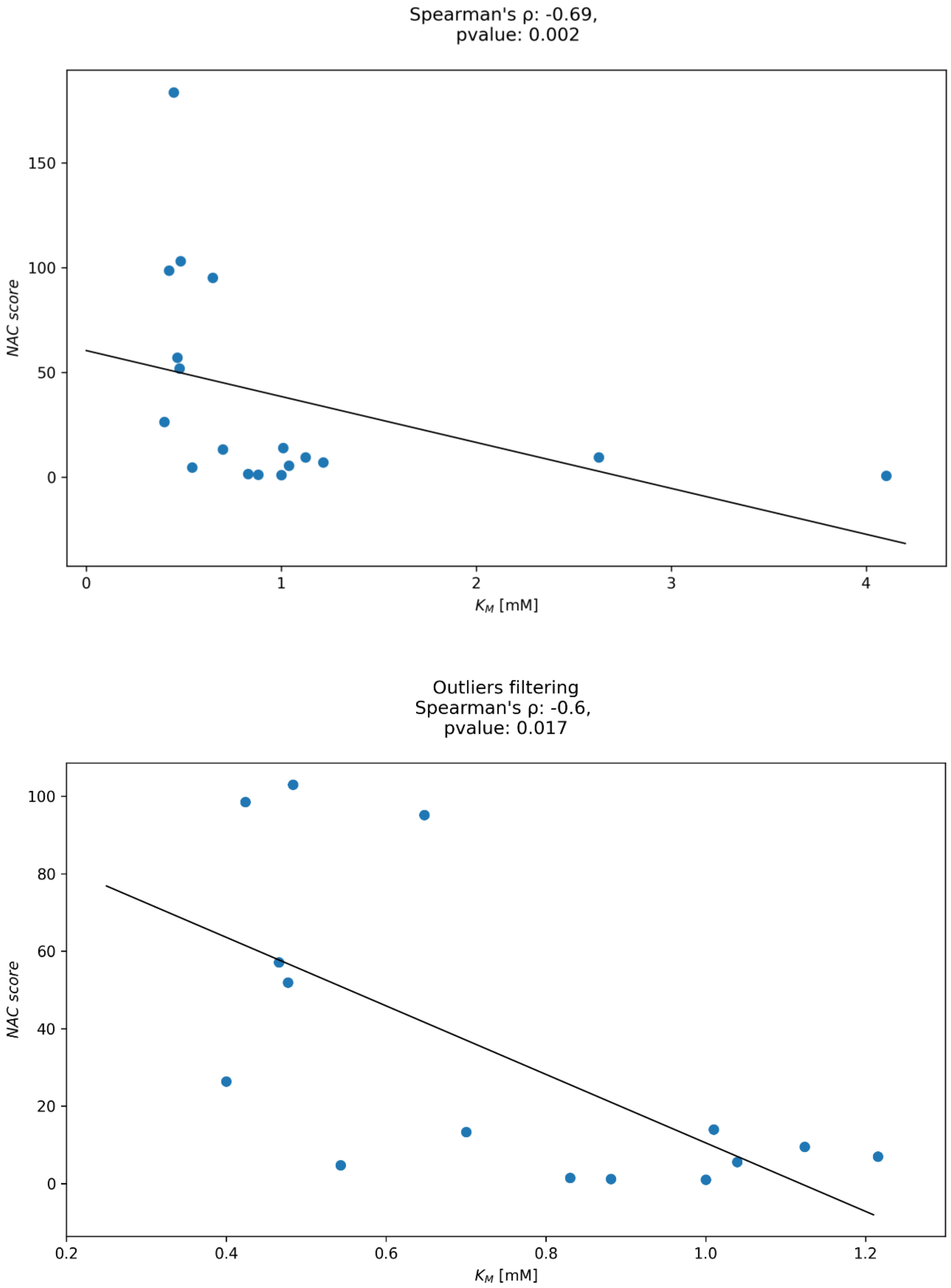


Figure S6. Spearman's correlation between experimental relative $K_M$ and computationally predicted capabilities to maintain productive stabilization relative to wild-type ecPGA (reactive stabilization score): top – full dataset, bottom: without outliers.

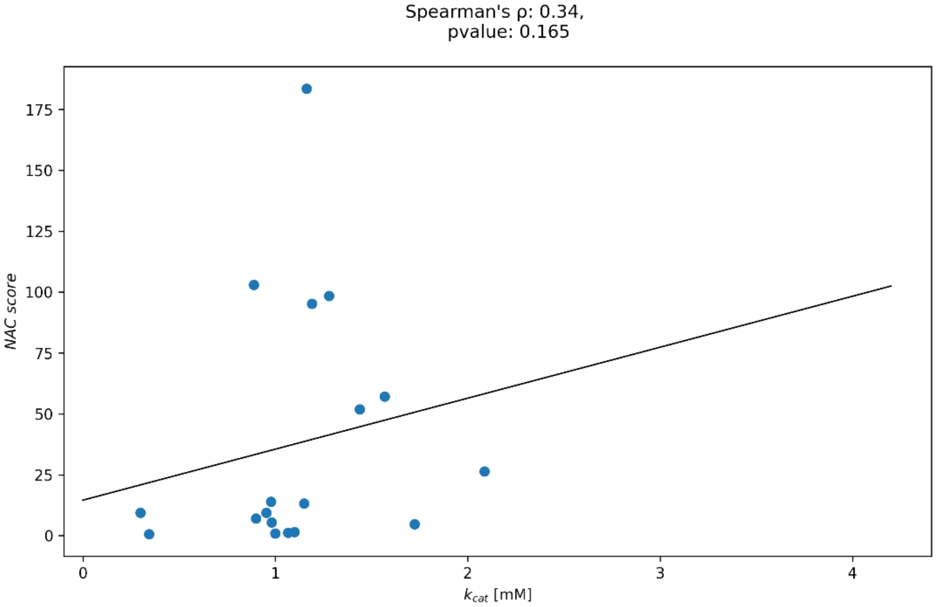


Figure S7. Spearman's correlation between experimental relative $k_{cat}$ and computationally predicted capabilities to maintain productive stabilization relative to wild-type ecPGA (reactive stabilization score).

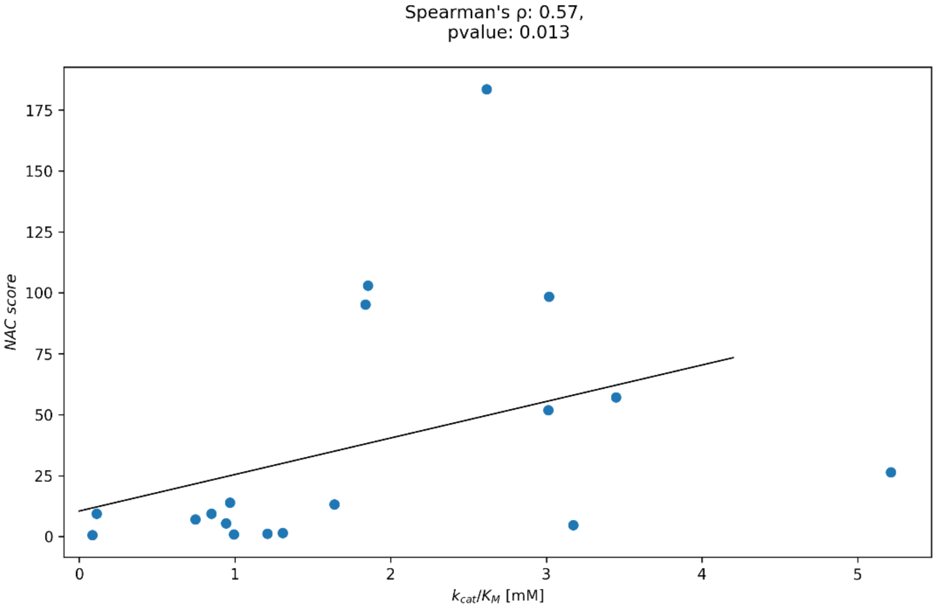


Figure S8. Spearman's correlation between experimental relative $k_{cat}/K_M$ and computationally predicted capabilities to maintain productive stabilization relative to wild-type ecPGA (reactive stabilization score).

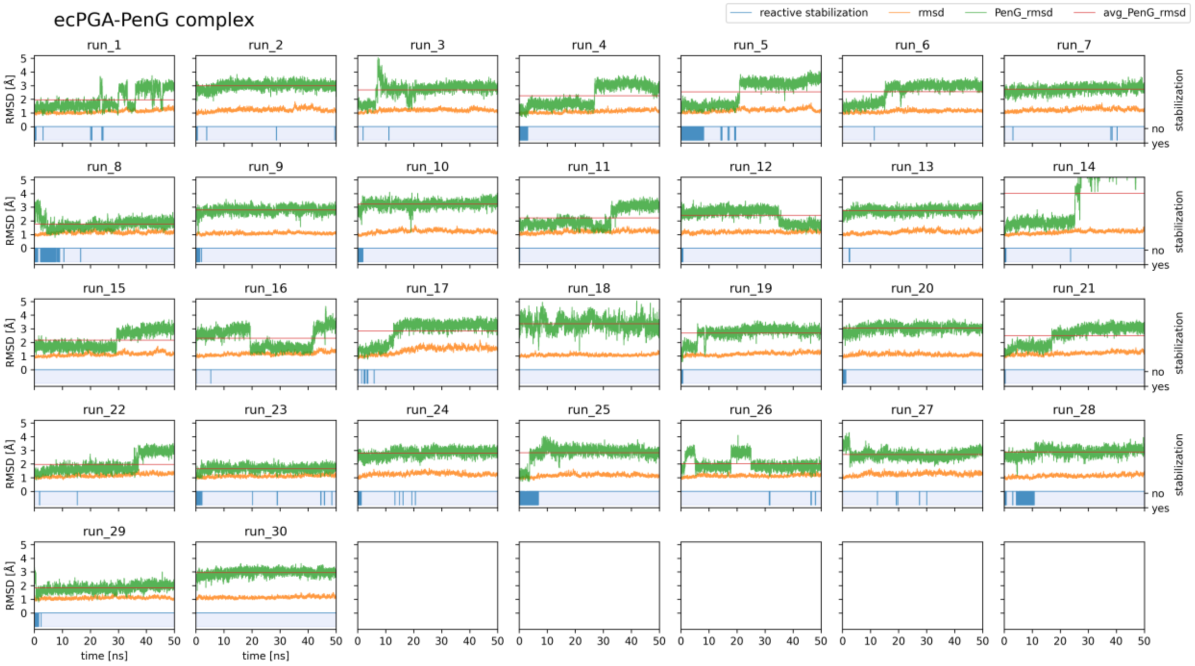


**Figure S9. Stabilization of ecPGA-PenG complex from repetitive MD simulations**. On each of particular subplots green curve corresponds to PenG heavy atoms root mean square deviation (RMSD), orange curve corresponds to protein backbone RMSD, red continuous line is an PenG average RMSD (left Y-axes). Reactive stabilization defined as properly stabilized PenG by interactions with oxyanion hole stabilizing residues Ala69β and Asn241β, and exhibiting nucleophile attack distance and angle within 3.3 Å and 75-105°, respectively, is shown as blue line on blue area of the plot (right Y-axes).

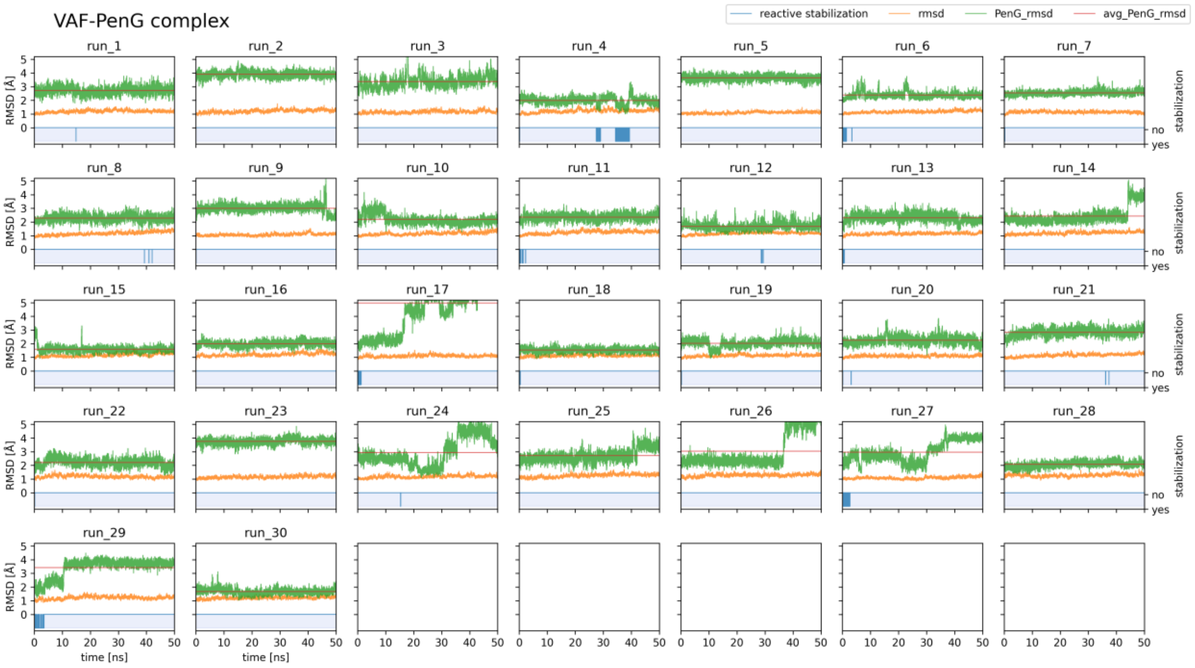


**Figure S10. Stabilization of VAF-PenG complex from repetitive MD simulations**. On each of particular subplots green curve corresponds to PenG heavy atoms root mean square deviation (RMSD), orange curve corresponds to protein backbone RMSD, red continuous line is an PenG average RMSD (left Y-axes). Reactive stabilization defined as properly stabilized PenG by interactions with oxyanion hole stabilizing residues Ala69β and Asn241β, and exhibiting nucleophile attack distance and angle within 3.3 Å and 75-105°, respectively, is shown as blue line on blue area of the plot (right Y-axes).

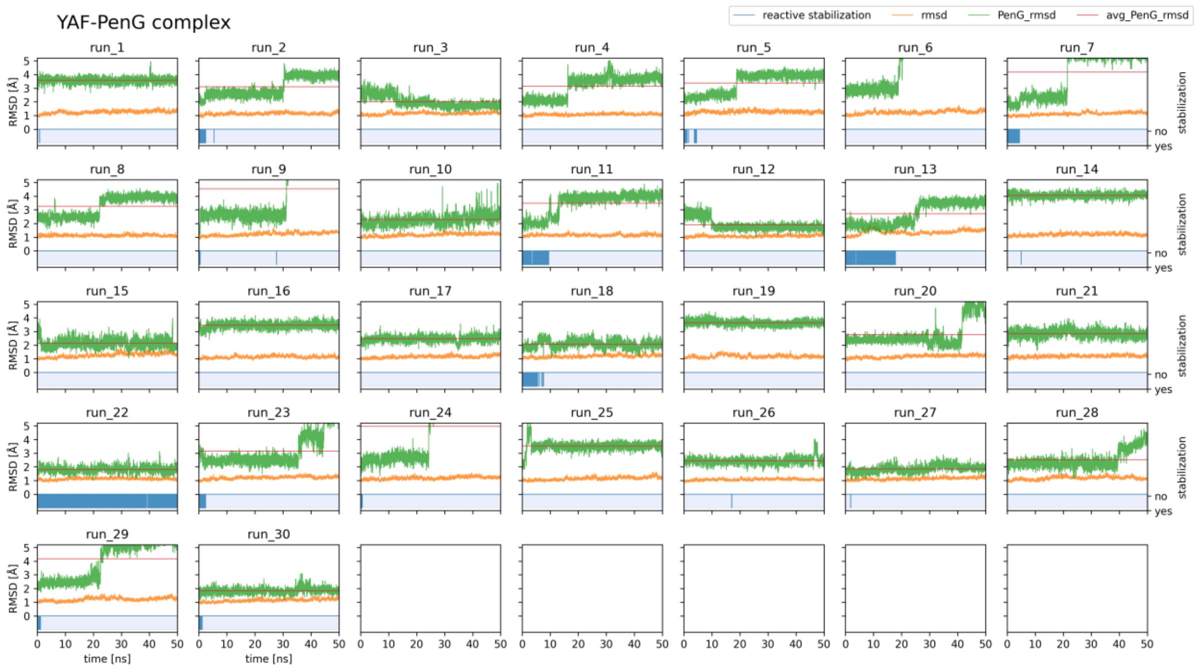


**Figure S11. Stabilization of YAF-PenG complex from repetitive MD simulations.** On each of particular subplots green curve corresponds to PenG heavy atoms root mean square deviation (RMSD), orange curve corresponds to protein backbone RMSD, red continuous line is an PenG average RMSD (left Y-axes). Reactive stabilization defined as properly stabilized PenG by interactions with oxyanion hole stabilizing residues Ala69β and Asn241β, and exhibiting nucleophile attack distance and angle within 3.3 Å and 75-105°, respectively, is shown as blue line on blue area of the plot (right Y-axes).

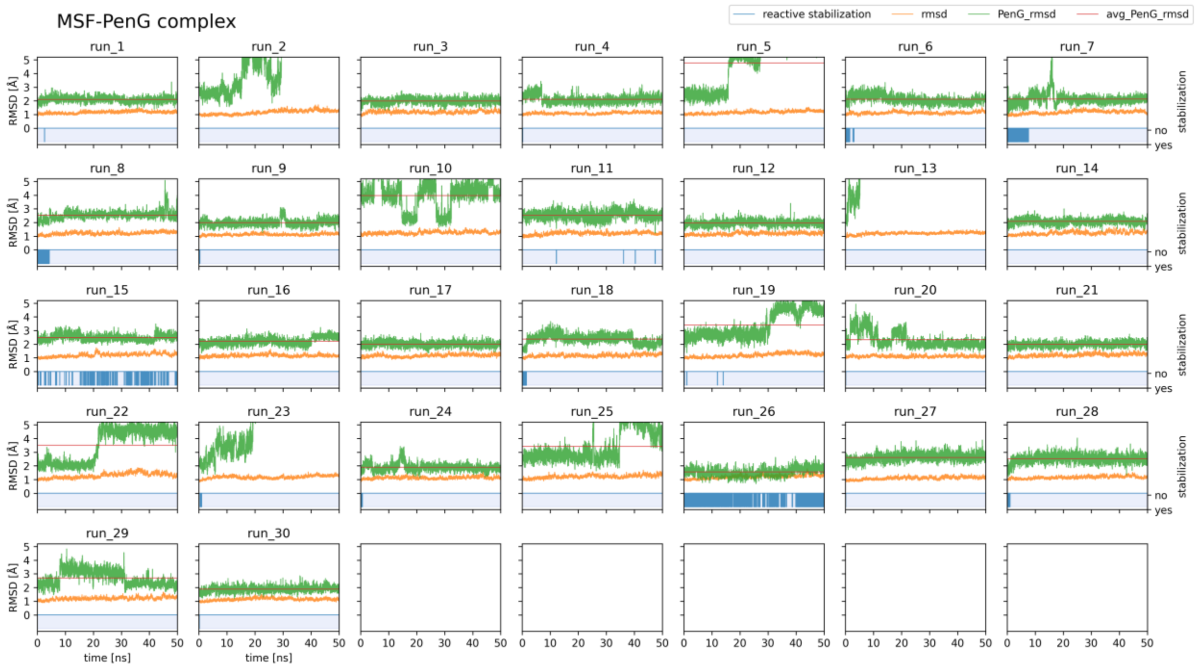


**Figure S12. Stabilization of MSF-PenG complex from repetitive MD simulations.** On each of particular subplots green curve corresponds to PenG heavy atoms root mean square deviation (RMSD), orange curve corresponds to protein backbone RMSD, red continuous line is an PenG average RMSD (left Y-axes). Reactive stabilization defined as properly stabilized PenG by interactions with oxyanion hole stabilizing residues Ala69β and Asn241β, and exhibiting nucleophile attack distance and angle within 3.3 Å and 75-105°, respectively, is shown as blue line on blue area of the plot (right Y-axes).

Table S8. Reactive stabilization score statistics for PenG complexes with ecPGA and three best mutants.

| protein | substrate | RSS | RSS filtered outliers | RSS filtered outliers leave-one-out | RSS filtered outliers leave-one-out std | RSS filtered outiers leave-one-out n samples | mutant vs wt ratio | mutant to wt difference statistics | mutant to wt difference pvalue |
|---|---|---|---|---|---|---|---|---|---|
| ecPGA | PEG | 2.1240 | 1.7428 | 1.7428 | 0.0930 | 29 | 1.00 | 0.00 | 1.00E+00 |
| VAF | PEG | 0.5432 | 0.3264 | 0.3265 | 0.0326 | 29 | 0.19 | -77.39 | 1.25E-58 |
| YAF | PEG | 2.1324 | 0.9499 | 0.9549 | 0.0892 | 28 | 0.55 | -32.61 | 1.15E-37 |
| MSF | PEG | 0.7619 | 0.4310 | 0.4318 | 0.0360 | 29 | 0.25 | -70.78 | 1.77E-56 |

Statistical test calculated using two-sided ttest, as implemented in python scipy library (ttest_ind_from_stats).

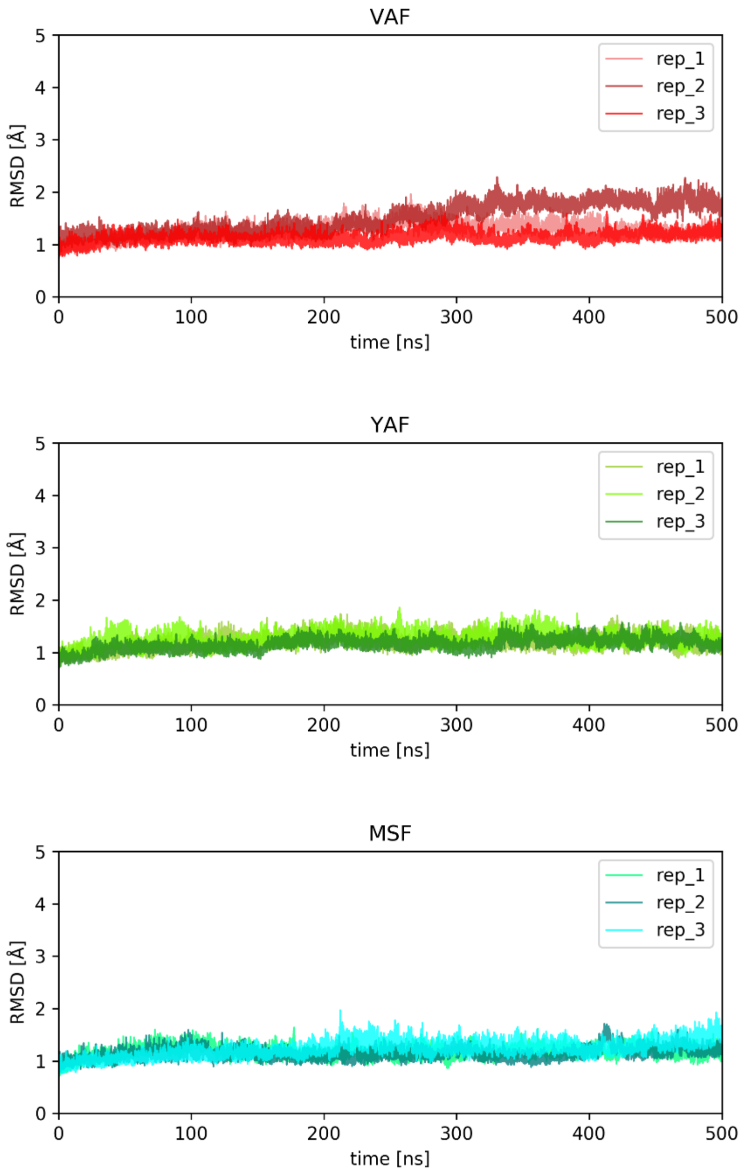


Figure S13. Free enzyme RMSD without C-terminal tail (200-207).

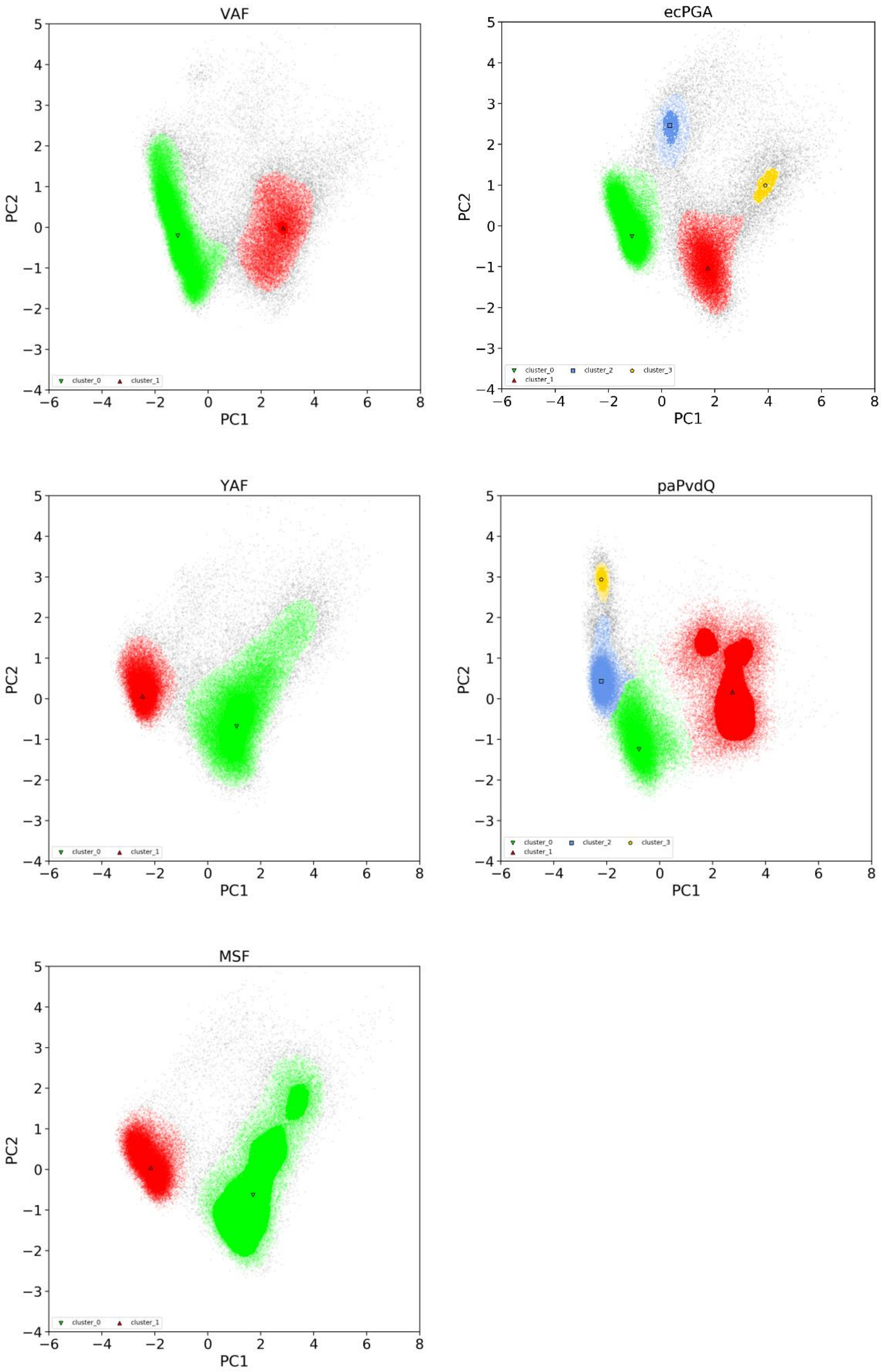


Figure S14. 1st and 2nd principal component's clusters. Black contour shapes represent the cluster representative.

**Table S9. Principal component analysis of the distances between functional atoms of catalytic machinery for shortlisted best three variants and wild-type ecPGA and paPvdQ enzymes.** Rows highlighted in bold represent clusters of state where the conformations of the residues forming the catalytic machinery are favorably pre-organized for productive binding of AHLs.

| Principle component cluster | Ser1β-NH1 – Ser1β-O | Ser1β-NH2 – Ser1β-O | Ser1β-OH – Ser1β-N | Asn241/269β-Nδ-Ala69/Val70β-N | Asn241/269β-ND - Ser1β-O | Ala69/Val70β-N - Ser1β-O | Gln23/His23β-O - Ser1β-O | Members (%) |
|---|---|---|---|---|---|---|---|---|
| | | | | ecPGA | | | | |
| 0_mean | 2.96 | 3.07 | 2.62 | 3.55 | 2.99 | 3.04 | 5.45 | 61.2 |
| 0_std | 0.51 | 0.54 | 0.27 | 0.18 | 0.15 | 0.15 | 0.38 | |
| **1_mean** | **3.35** | **3.34** | **2.72** | **4.38** | **4.55** | **4.79** | **3.93** | **23.8** |
| 1_std | 0.49 | 0.48 | 0.51 | 0.30 | 0.33 | 0.33 | 0.27 | |
| 2_mean | 3.05 | 3.25 | 2.45 | 5.86 | 4.45 | 3.01 | 6.85 | 2.7 |
| 2_std | 0.54 | 0.56 | 0.22 | 0.22 | 0.18 | 0.16 | 0.26 | |
| 3_mean | 3.27 | 3.10 | 2.81 | 6.97 | 6.05 | 5.14 | 4.28 | 0.9 |
| 3_std | 0.47 | 0.50 | 0.59 | 0.39 | 0.45 | 0.43 | 0.24 | |
| | | | | VAF | | | | |
| 0_mean | 2.98 | 3.10 | 2.59 | 3.59 | 2.98 | 3.06 | 5.72 | 67.9 |
| 0_std | 0.51 | 0.53 | 0.26 | 0.19 | 0.14 | 0.15 | 0.63 | |
| **1_mean** | **3.19** | **3.19** | **2.68** | **5.31** | **5.20** | **5.49** | **4.29** | **21.3** |
| 1_std | 0.48 | 0.53 | 0.47 | 0.77 | 0.52 | 0.58 | 0.27 | |
| | | | | YAF | | | | |
| **0_mean** | **3.38** | **3.36** | **2.51** | **4.96** | **4.89** | **5.36** | **3.93** | **59.3** |
| 0_std | 0.38 | 0.38 | 0.46 | 0.50 | 0.39 | 0.37 | 0.34 | |
| 1_mean | 3.04 | 2.94 | 2.55 | 3.66 | 2.98 | 3.06 | 5.42 | 33.8 |
| 1_std | 0.54 | 0.50 | 0.25 | 0.11 | 0.13 | 0.14 | 0.21 | |
| | | | | MSF | | | | |
| **0_mean** | **3.27** | **3.31** | **2.60** | **4.94** | **5.04** | **5.65** | **4.11** | 51.1 |
| 0_std | 0.46 | 0.44 | 0.54 | 1.14 | 0.48 | 0.39 | 0.46 | |
| 1_mean | 3.07 | 3.06 | 2.64 | 3.62 | 3.01 | 3.07 | 5.80 | 45.1 |
| 1_std | 0.57 | 0.56 | 0.38 | 0.11 | 0.17 | 0.16 | 0.61 | |
| | | | | paPvdQ | | | | |
| **0_mean** | **3.46** | **3.43** | **2.27** | **3.72** | **4.31** | **4.10** | **3.81** | **35.7** |
| 0_std | 0.21 | 0.22 | 0.27 | 0.22 | 0.22 | 0.29 | 0.19 | |
| 1_mean | 3.55 | 3.54 | 2.50 | 6.76 | 6.51 | 3.74 | 4.35 | 34.1 |
| 1_std | 0.37 | 0.39 | 0.70 | 0.31 | 0.46 | 0.51 | 0.67 | |
| 2_mean | 3.35 | 3.26 | 2.44 | 3.53 | 2.96 | 3.06 | 5.27 | 25.8 |
| 2_std | 0.45 | 0.46 | 0.24 | 0.11 | 0.12 | 0.16 | 0.12 | |
| 3_mean | 3.80 | 3.14 | 3.41 | 4.08 | 2.93 | 3.32 | 7.88 | 1.3 |
| 3_std | 0.36 | 0.38 | 0.17 | 0.14 | 0.12 | 0.26 | 0.19 | |

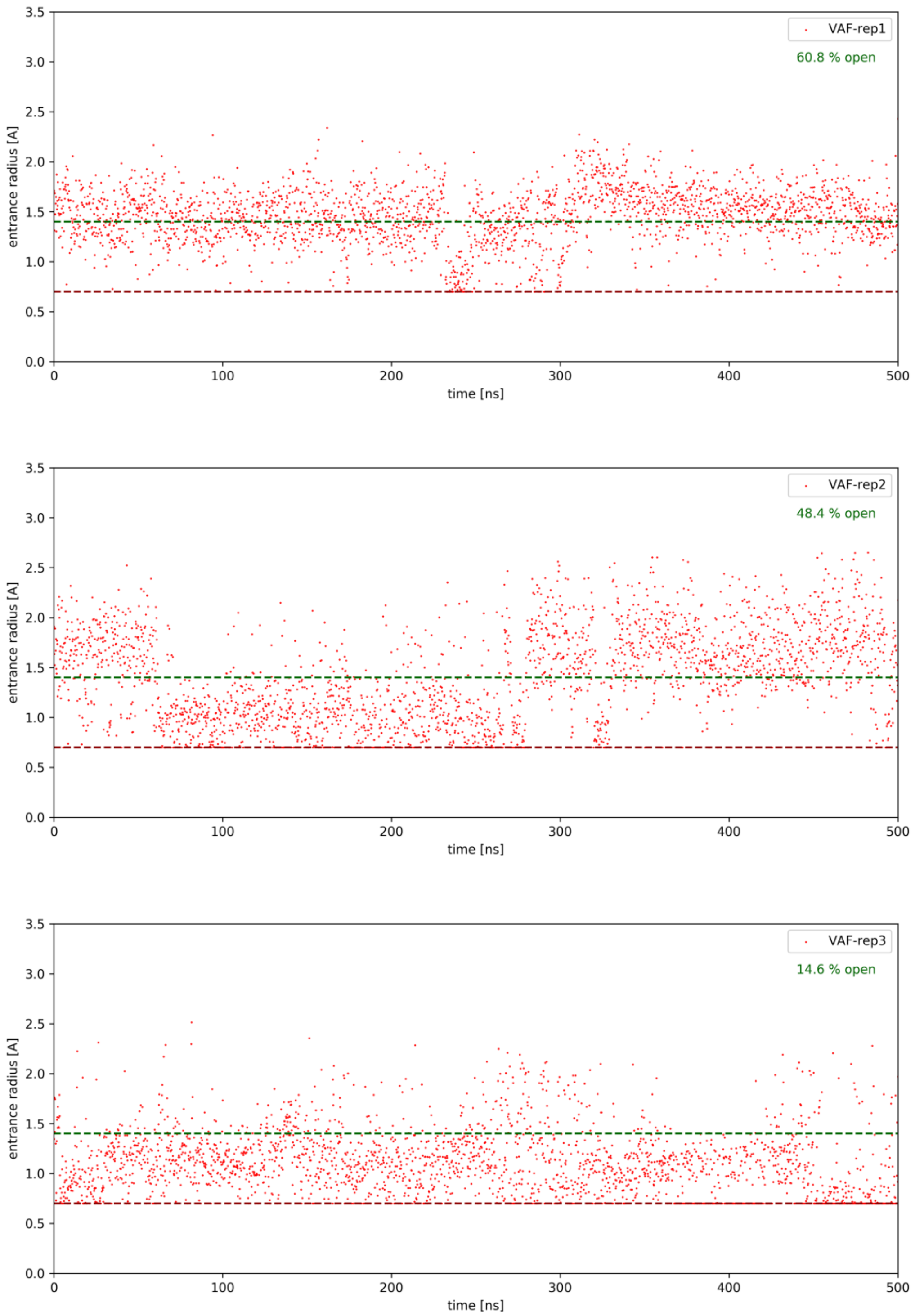


Figure S15. VAF acyl-binding site entrance time-evolution.

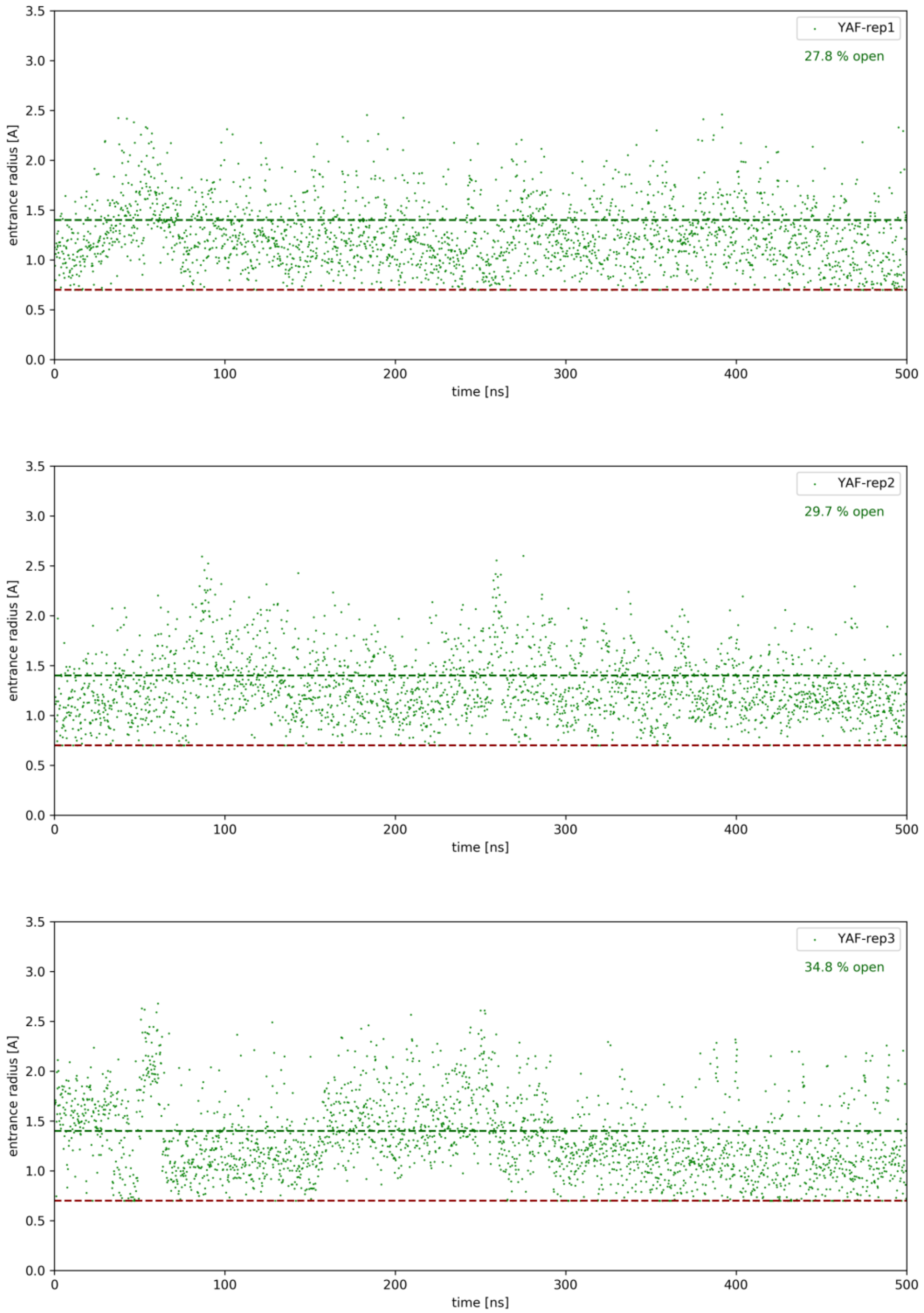


Figure S16. YAF acyl-binding site entrance time-evolution.

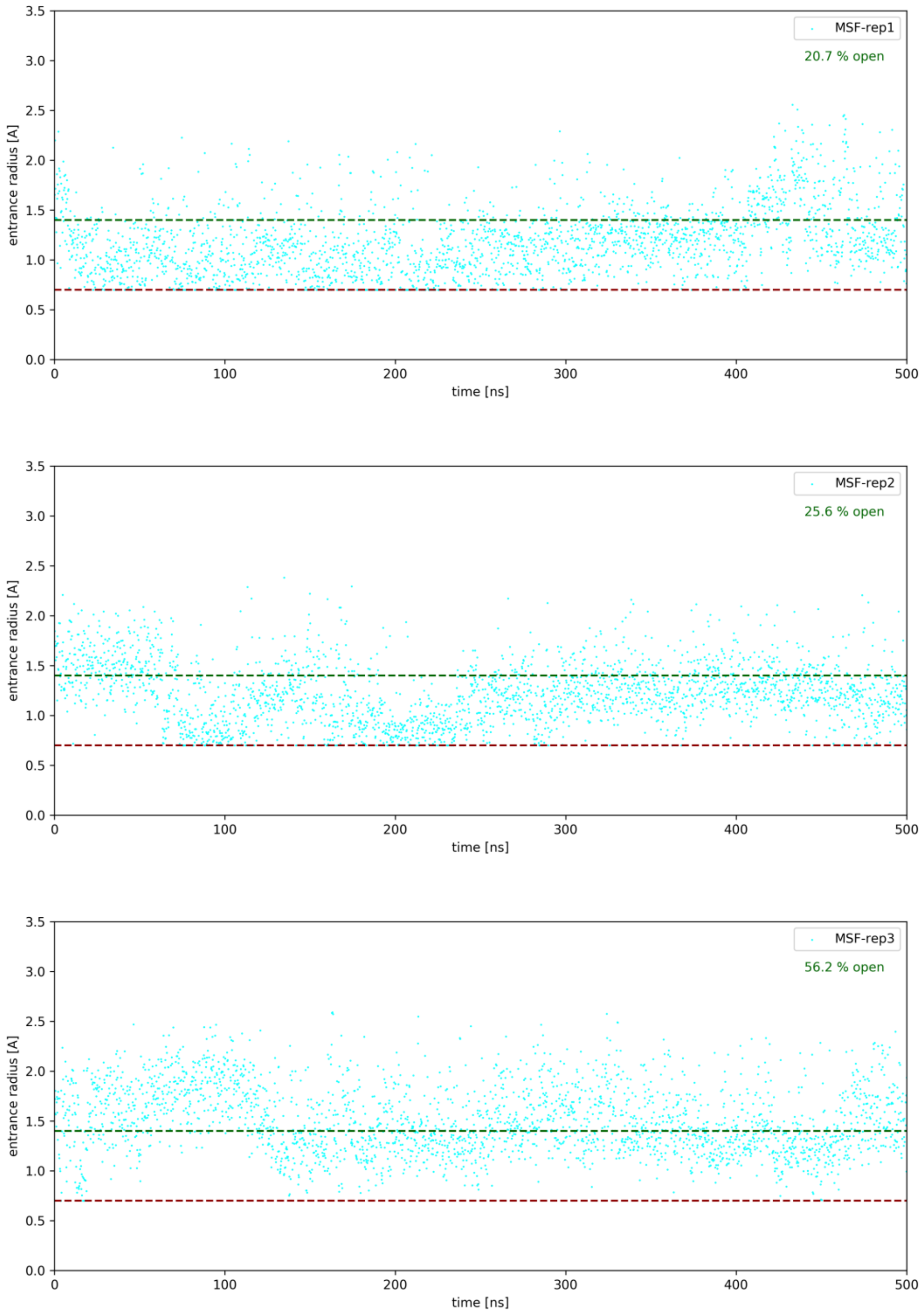


Figure S17. MSF acyl-binding site entrance time-evolution.

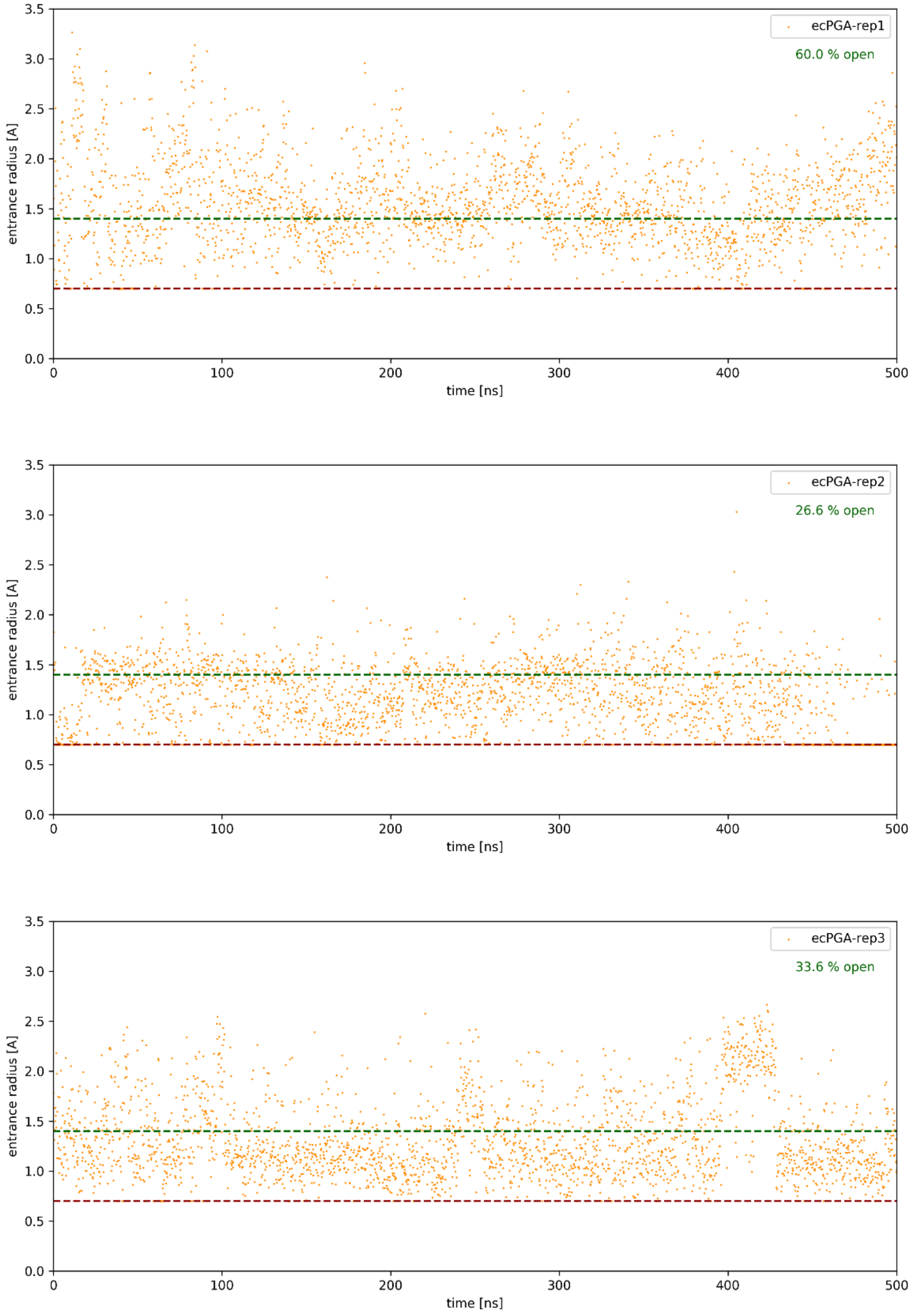


Figure S18. ecPGA acyl-binding site entrance time-evolution.

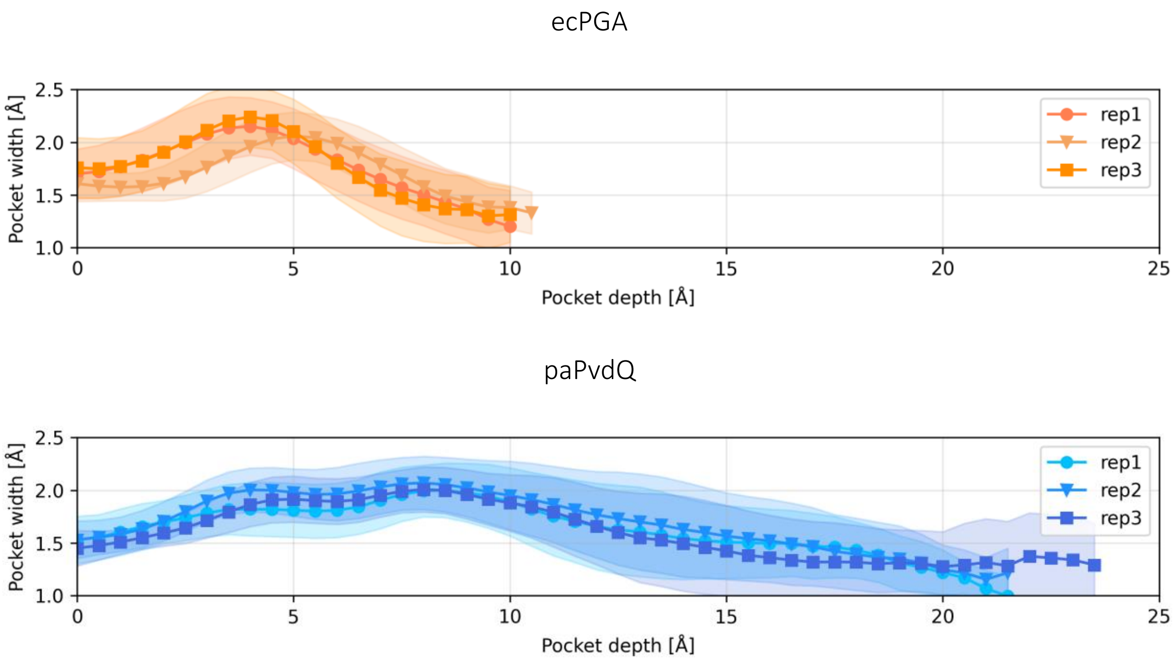


Figure S19. Profiles of pockets in wild-type ecPGA and paPvdQ.

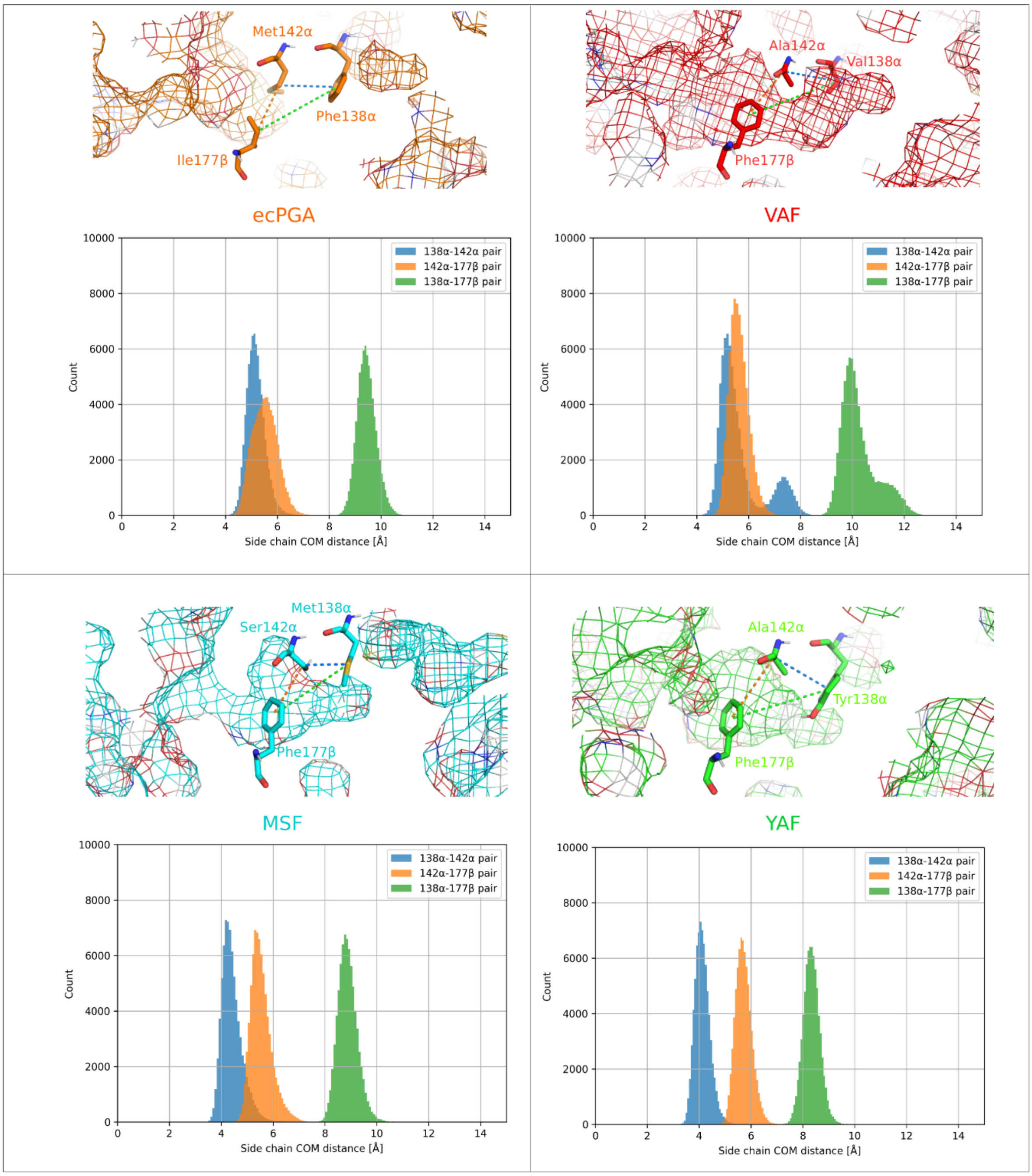


**Figure S20. Distribution of the distances between center of mass (COM) for side chains of introduced mutations.** The pocket is represented as isomesh representation, while the mutated residues are represented as sticks.

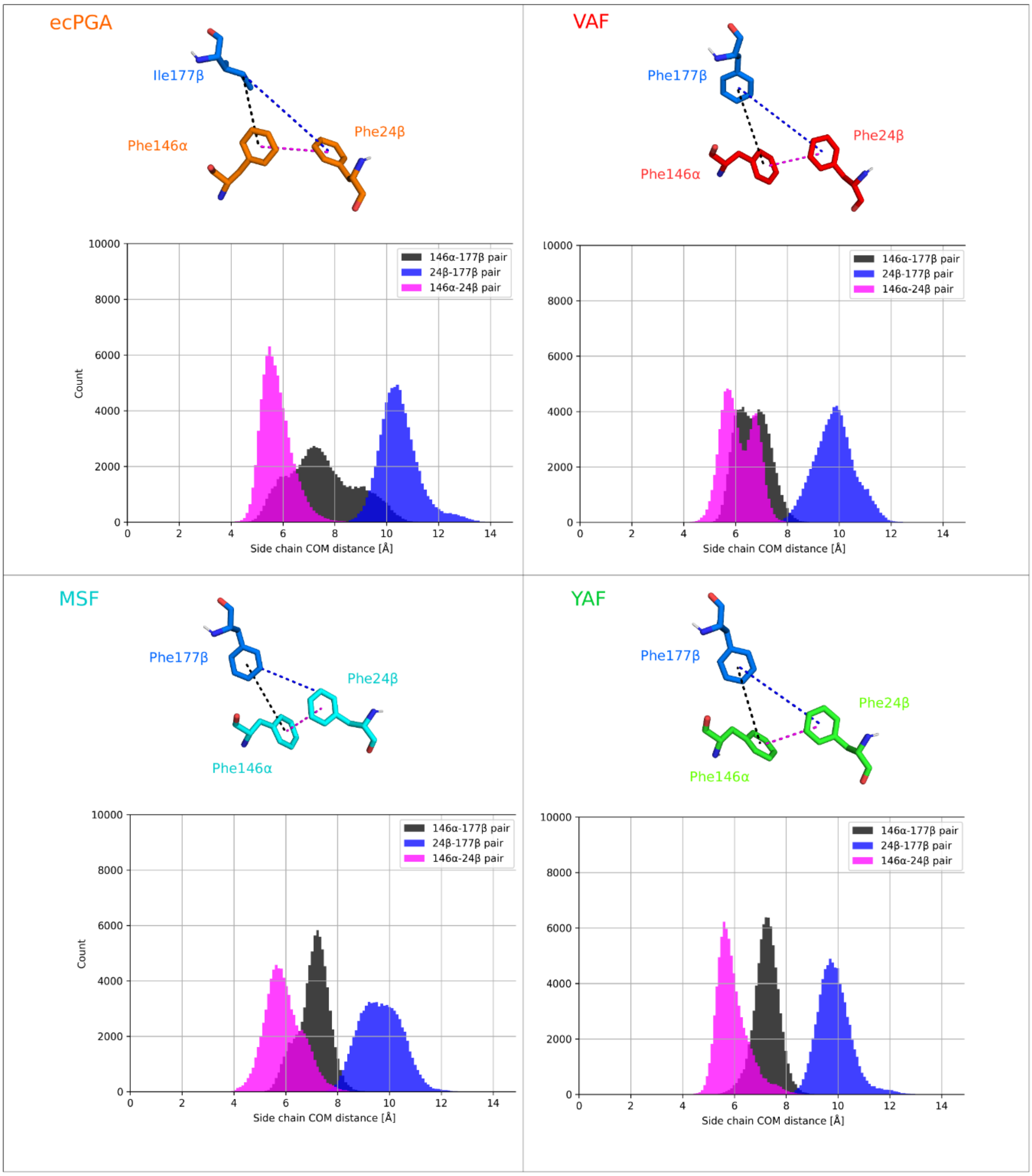


Figure S21. Distribution of the distances between center of mass (COM) for side chains of gating residues Phe146α, Phe24β and introduced Phe177β, which are represented as sticks.

Table S10. Crucial average distances representing components of reactive stabilization, averaged from the ensembles of properly stabilized substrate (reactive stabilization ensembles).

| Distance [Å] | Ala69/Val70β –NH – HSL-O | Gln23β/His23 β-O – HSL-NH | Ser1β-O – HSL-C | Asn241/269β-NH2(min) – HSL-O | Ser1β-O > HSL-C > HSL-O |
|---|---|---|---|---|---|
| VAF_C06_complex | | | | | |
| Mean | 2.2 | 3.5 | 3.1 | 2.1 | 91 |
| Std | 0.3 | 1.3 | 0.1 | 0.2 | 7 |
| VAF_C06-3O_complex | | | | | |
| Mean | 2.4 | 2.3 | 3.1 | 2.3 | 89 |
| Std | 0.3 | 0.6 | 0.1 | 0.3 | 7 |
| VAF_C08_complex | | | | | |
| Mean | 2.2 | 2.4 | 3.1 | 2.1 | 89 |
| Std | 0.3 | 0.7 | 0.1 | 0.2 | 6 |
| VAF_C08-3O_complex | | | | | |
| Mean | 2.3 | 3.0 | 3.1 | 2.1 | 89 |
| Std | 0.3 | 0.8 | 0.1 | 0.2 | 7 |
| VAF_C10_complex | | | | | |
| Mean | 2.9 | 2.6 | 3.1 | 2.1 | 83 |
| Std | 0.1 | 0.8 | 0.1 | 0.3 | 4 |
| VAF_C12-3O_complex | | | | | |
| Mean | 2.6 | 2.2 | 3.1 | 2.3 | 89 |
| Std | 0.3 | 0.4 | 0.1 | 0.4 | 7 |
| YAF_C06_complex | | | | | |
| Mean | 2.1 | 3.7 | 3.1 | 2.1 | 91 |
| Std | 0.3 | 0.9 | 0.1 | 0.2 | 7 |
| YAF_C06-3O_complex | | | | | |
| Mean | 2.3 | 2.7 | 3.1 | 2.2 | 90 |
| Std | 0.3 | 1.0 | 0.1 | 0.3 | 6 |
| YAF_C08_complex | | | | | |
| Mean | 2.3 | 2.4 | 3.1 | 2.1 | 88 |
| Std | 0.3 | 0.7 | 0.1 | 0.3 | 7 |
| YAF_C08-3O_complex | | | | | |
| Mean | 2.3 | 3.1 | 3.1 | 2.1 | 89 |
| Std | 0.3 | 0.9 | 0.1 | 0.2 | 7 |
| YAF_C10_complex | | | | | |
| Mean | 2.2 | 2.6 | 3.1 | 2.0 | 90 |
| Std | 0.3 | 0.7 | 0.1 | 0.2 | 7 |
| YAF_C12-3O_complex | | | | | |
| Mean | 2.3 | 2.2 | 3.1 | 2.2 | 89 |
| Std | 0.3 | 0.5 | 0.1 | 0.3 | 6 |
| MSF_C06_complex | | | | | |
| Mean | 2.3 | 3.2 | 3.1 | 2.1 | 92 |
| Std | 0.3 | 1.0 | 0.1 | 0.2 | 7 |
| MSF_C06-3O_complex | | | | | |
| Mean | 2.4 | 2.5 | 3.1 | 2.3 | 90 |
| Std | 0.3 | 0.8 | 0.1 | 0.3 | 7 |
| MSF_C08_complex | | | | | |
| Mean | 2.1 | 3.8 | 3.1 | 2.1 | 92 |
| Std | 0.3 | 1.1 | 0.1 | 0.2 | 7 |
| MSF_C08-3O_complex | | | | | |
| Mean | 2.3 | 2.6 | 3.1 | 2.2 | 88 |
| Std | 0.3 | 0.8 | 0.1 | 0.2 | 7 |
| MSF_C10_complex | | | | | |
| Mean | 2.6 | 2.4 | 3.1 | 2.2 | 90 |
| Std | 0.3 | 0.9 | 0.1 | 0.3 | 6 |
| MSF_C12-3O_complex | | | | | |
| Mean | 2.5 | 2.6 | 3.1 | 2.2 | 89 |
| Std | 0.3 | 0.8 | 0.1 | 0.3 | 7 |

Table S11. Frequency of the 3-oxo hydrogen bonding as a fraction of simulations that substrate average RMSD did not exceed 3 Å, which occurred in at least of 5 % of these simulations in any of the complexes.

| complex | Ala69β-NH – HSL-3-oxo | Gln23β – HSL-3-oxo | Ser1β-NH2 – HSL-3-oxo |
|---|---|---|---|
| VAF-C06-3O | 0.04 | - | - |
| VAF-C08-3O | 0.24 | - | 0.00 |
| VAF-C12-3O | 0.03 | 0.00 | - |
| YAF-C06-3O | 0.13 | - | - |
| YAF-C08-3O | 0.21 | - | 0.00 |
| YAF-C12-3O | 0.07 | - | - |
| MSF-C06-3O | 0.18 | 0.01 | 0.00 |
| MSF-C08-3O | 0.23 | 0.00 | 0.01 |
| MSF-C12-3O | 0.30 | 0.02 | 0.00 |
| ecPGA-C06-3O | 0.07 | 0.02 | 0.00 |
| ecPGA-C08-3O | 0.12 | 0.00 | 0.01 |
| ecPGA-C12-3O | - | 0.28 | 0.10 |

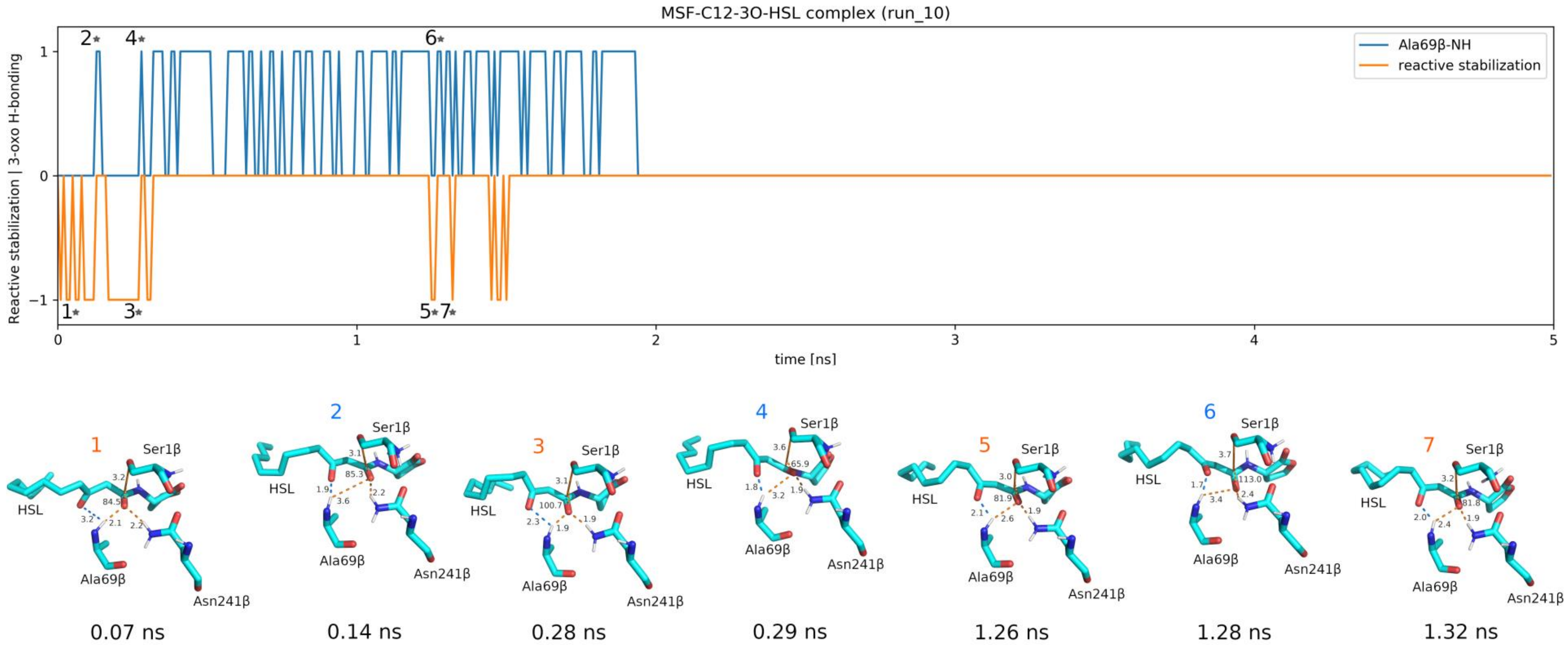


**Figure S22. Reoccurring reactive stabilization after hydrogen bonding event of catalytic machinery with 3-oxo group.** Our simulations show, that even in the mutants with relatively constrained pocket for longer substrates (C10-, C12-3O-HSL), the 3-oxo substituent improves the occurrence of the reactive stabilization and limits the complete unbinding of the substrate. As represented by the orange and blue curve, the hydrogen bonding of the 3-oxo group with Ala69β-NH enables the restoration of the reactive stabilization. States marked in the plot as * are visualized as sticks below the time-evolution plot. Orange dashed lines represent the crucial components of the reactive stabilization, while the blue dashed line corresponds to the Ala69β-NH distance to 3-oxo substituent.

Table S12. List of primers.

| ecPGA cloning primers | |
|---|---|
| pET_PGA_F | CCTCTAGAAATAATTTTGTTTAACTTTAAGAAGGAGATATA**CATATG**AAAAATAGAAATCGTATGATCGTGAAC |
| pET_PGA_R | GGATCTCAGTGGTGGTGGTGGTGGTG**CTCGAG**TTATCTCTGAACGTGCAACACTTCC |
| pET PGA R-His variant | GGATCTCAGTGGTGGTGGTGGTGGTG**CTCGAG**TCTCTGAACGTGCAACACTTCC |
| Mutagenic primers | |
| MSF Fwd | CTGGGAACCGTTTGATGTCGCGATGATAATGGTGGGCACCAGCGCAAACCGCTTCTCTGATAGC |
| YAF Fwd | CTGGGAACCGTTTGATGTCGCGATGATATATGTGGGCACCGCGGCAAACCGCTTCTCTGATAGC |
| VAF Fwd | CTGGGAACCGTTTGATGTCGCGATGATAGTGGTGGGCACCGCGGCAAACCGCTTCTCTGATAGC |
| MutPGAuni_R | GCCGTTTACATCAGCATAGTACCAGTTGAAGGTCAGTGCTTGTTTCGCTGCC |
| Sequencing primers | |
| MSF check | ATGGTGGGCACCAGC |
| YAF check | TATGTGGGCACCGCG |
| VAF check | GTGGTGGGCACCGCG |
| PGAver_R | CATCAGCATAGTACCAGTTGAA |
| PGAseq_F | GCAAATTGCTGCCCTTTCCC |
| PGAseq_R | GGCAAACAGATCTGAAGCGG |

Restriction sites in bold

Table S13. Retention times of AHL substrates obtained from HPLC measurement.

| Substrate | Retention time (min) |
|---|---|
| C06-HSL | 15.8 |
| C06-3O-HSL | 9.3 |
| C08-HSL | 28.4 |
| C08-3O-HSL | 18.2 |
| C10-HSL | 33.6 |
| C12-3O-HSL | 34.3 |

Table S14. Retention times of reactants and products in synthesis of ampicillin and amoxicillin obtained from HPLC measurement.

| Activated acyl-donor (AD) | Retention time (min) | β-lactam nucleophile | Retention time (min) | Antibiotics | Retention time (min) |
|---|---|---|---|---|---|
| D-PGA | 10.3 | 6-APA | 6.7 | Ampicillin | 11.7 |
| D-HPGA | 7.6 | 6-APA | 8.7 | Amoxicillin | 20.0 |
