## Supplementary Information File 2 for "Engineering dynamic gates in binding pocket of penicillin G acylase to selectively degrade bacterial signaling molecules"

‡ These authors contributed equally.

This supplementary file includes multiplots related to all replicates of molecular dynamics simulations for all protein-substrate complexes. On each of particular subplots green curve corresponds to N-acyl-homoserine lactone (AHL) heavy atoms root mean square deviation (RMSD), orange curve corresponds to protein backbone RMSD, red continuous line is an AHL average RMSD (left Y-axes). Reactive stabilization defined as properly stabilized AHLs by interactions with oxyanion hole stabilizing residues Ala69β and Asn241β, and exhibiting nucleophile attack distance and angle within 3.3 Å and 75-105°, respectively, is shown as blue line on blue area of the plot (right Y-axes).

Full names of presented protein variants:

LAF, Phe138αLeu & Met142αAla & Ile177βPhe ecPGA variant;

MAF, Phe138αMet & Met142αAla & Ile177βPhe ecPGA variant;

VAF, Phe138αVal & Met142αAla & Ile177βPhe ecPGA variant;

YAF, Phe138αTyr & Met142αAla & Ile177βPhe ecPGA variant;

LSF, Phe138αLeu & Met142αSer & Ile177βPhe ecPGA variant;

MSF, Phe138αMet & Met142αSer & Ile177βPhe ecPGA variant

Full names of presented AHLs:

C06-HSL, N-hexanoyl-L-homoserine lactone;

C06-3O-HSL, N-3-oxo-hexanoyl-L-homoserine lactone;

C08-HSL, N-octanoyl-L-homoserine lactone;

C08-3O-HSL, N-3-oxo-octanoyl-L-homoserine lactone;

C10-HSL, N-decanoyl-L-homoserine lactone;

C12-3O-HSL, N-3-oxo-dodecanoyl-L-homoserine lactone

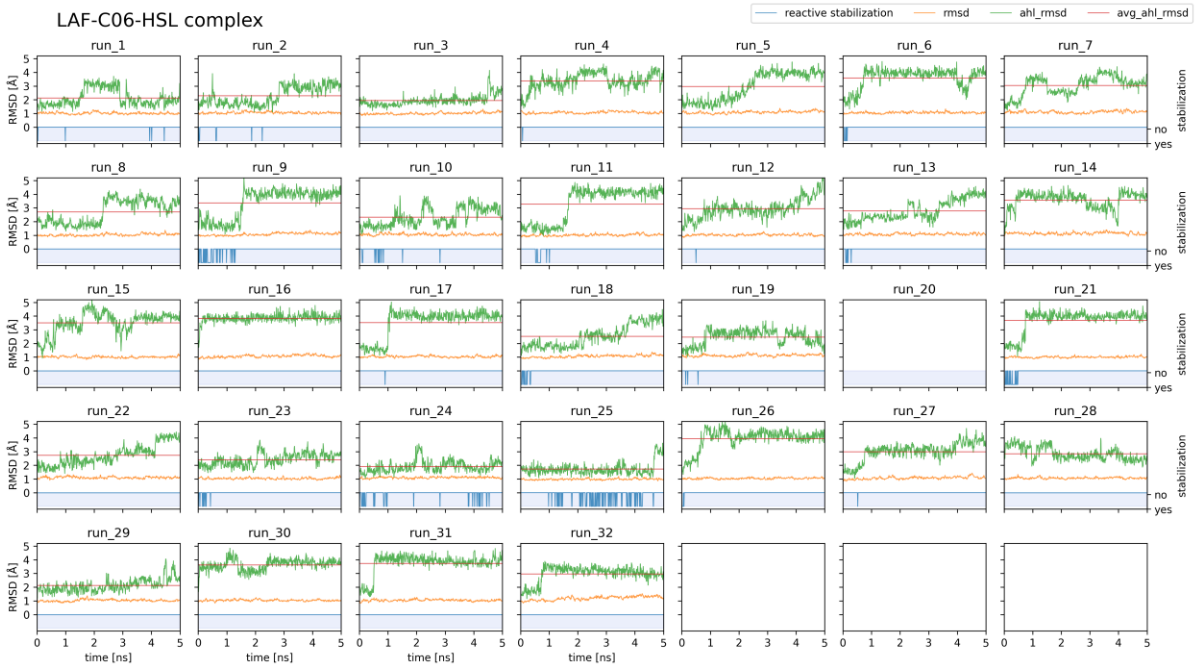
LAF-C06-HSL complex
reactive stabilization
rmsd
ahl_rmsd
avg_ahl_rmsd
run_1 run_2 run_3 run_4 run_5 run_6 run_7
run_8 run_9 run_10 run_11 run_12 run_13 run_14
run_15 run_16 run_17 run_18 run_19 run_20 run_21
run_22 run_23 run_24 run_25 run_26 run_27 run_28
run_29 run_30 run_31 run_32
RMSD [Å]
stabilization
no
yes
time [ns]

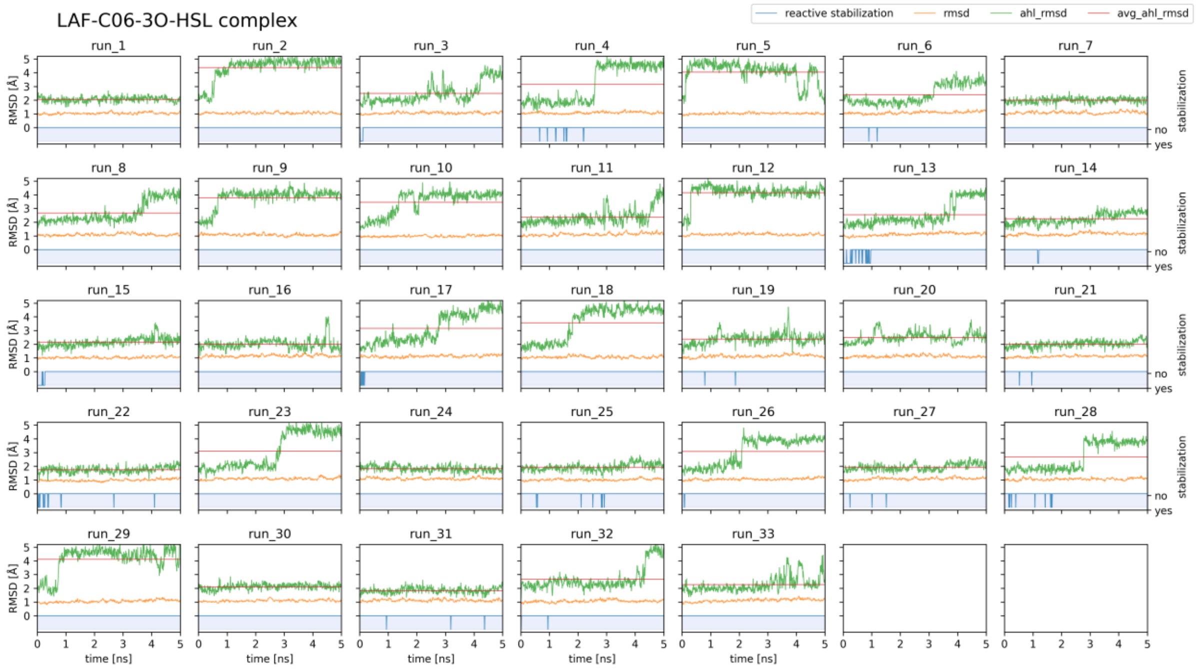
LAF-C06-3O-HSL complex
reactive stabilization
rmsd
ahl_rmsd
avg_ahl_rmsd
run_1 run_2 run_3 run_4 run_5 run_6 run_7
run_8 run_9 run_10 run_11 run_12 run_13 run_14
run_15 run_16 run_17 run_18 run_19 run_20 run_21
run_22 run_23 run_24 run_25 run_26 run_27 run_28
run_29 run_30 run_31 run_32 run_33
RMSD [Å]
stabilization
no
yes
time [ns]

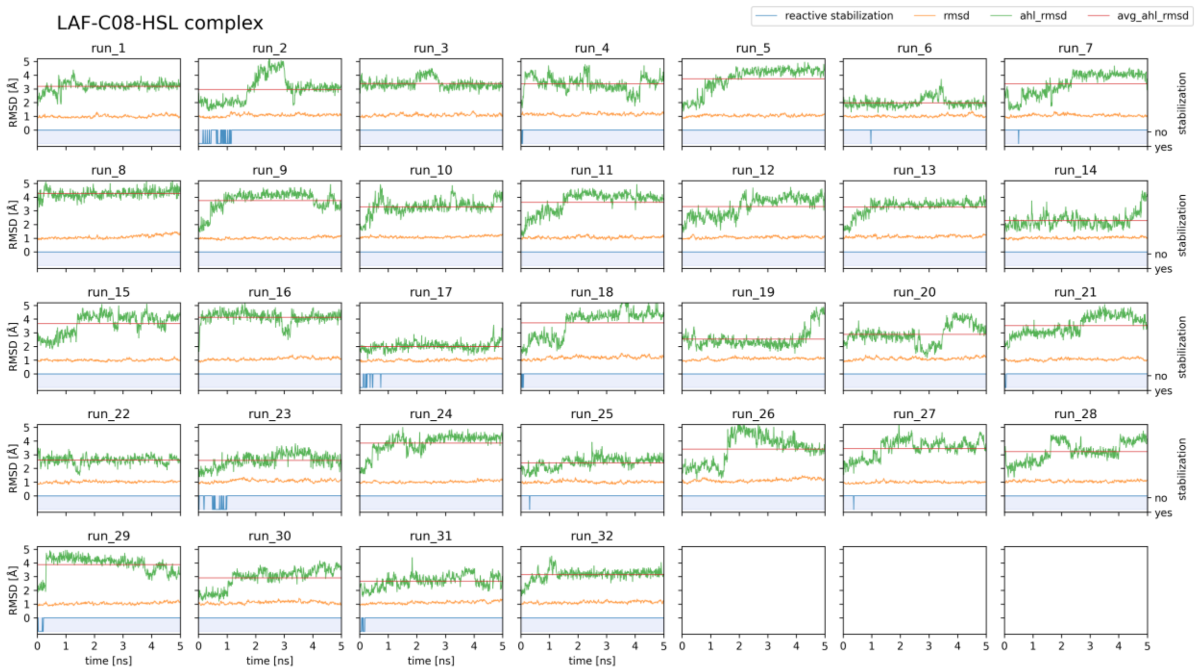
LAF-C08-HSL complex
reactive stabilization
rmsd
ahl_rmsd
avg_ahl_rmsd
run_1
run_2
run_3
run_4
run_5
run_6
run_7
run_8
run_9
run_10
run_11
run_12
run_13
run_14
run_15
run_16
run_17
run_18
run_19
run_20
run_21
run_22
run_23
run_24
run_25
run_26
run_27
run_28
run_29
run_30
run_31
run_32
RMSD [Å]
stabilization
no
yes
time [ns]

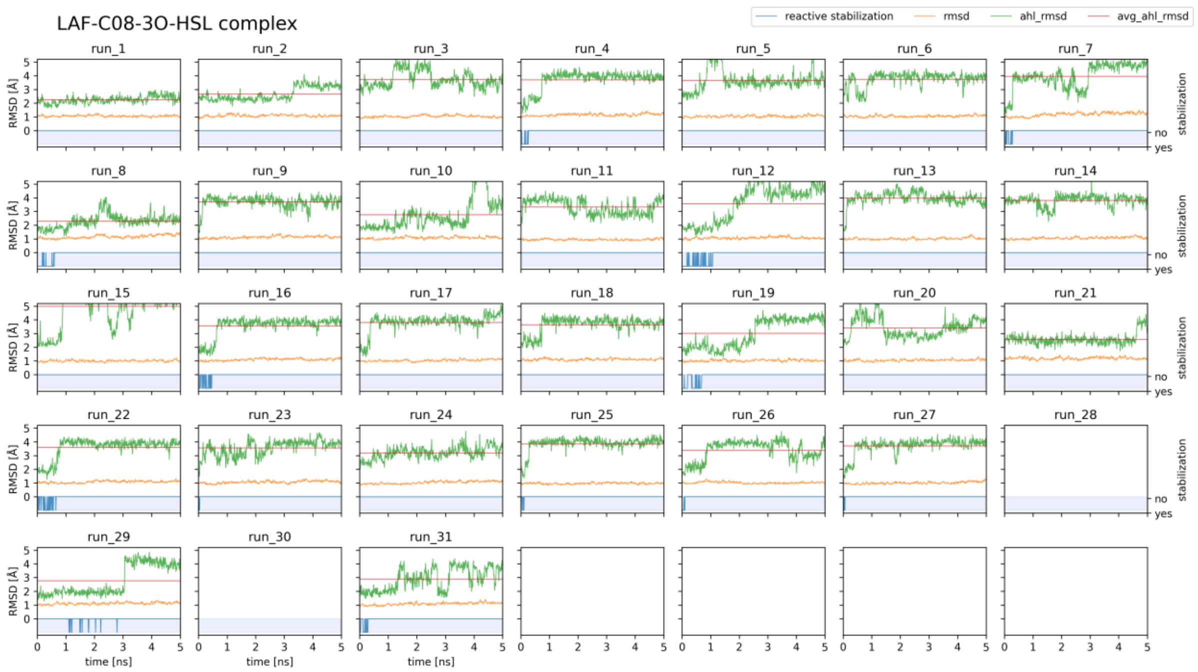
LAF-C08-3O-HSL complex
reactive stabilization
rmsd
ahl_rmsd
avg_ahl_rmsd
run_1
run_2
run_3
run_4
run_5
run_6
run_7
run_8
run_9
run_10
run_11
run_12
run_13
run_14
run_15
run_16
run_17
run_18
run_19
run_20
run_21
run_22
run_23
run_24
run_25
run_26
run_27
run_28
run_29
run_30
run_31
RMSD [Å]
stabilization
no
yes
time [ns]

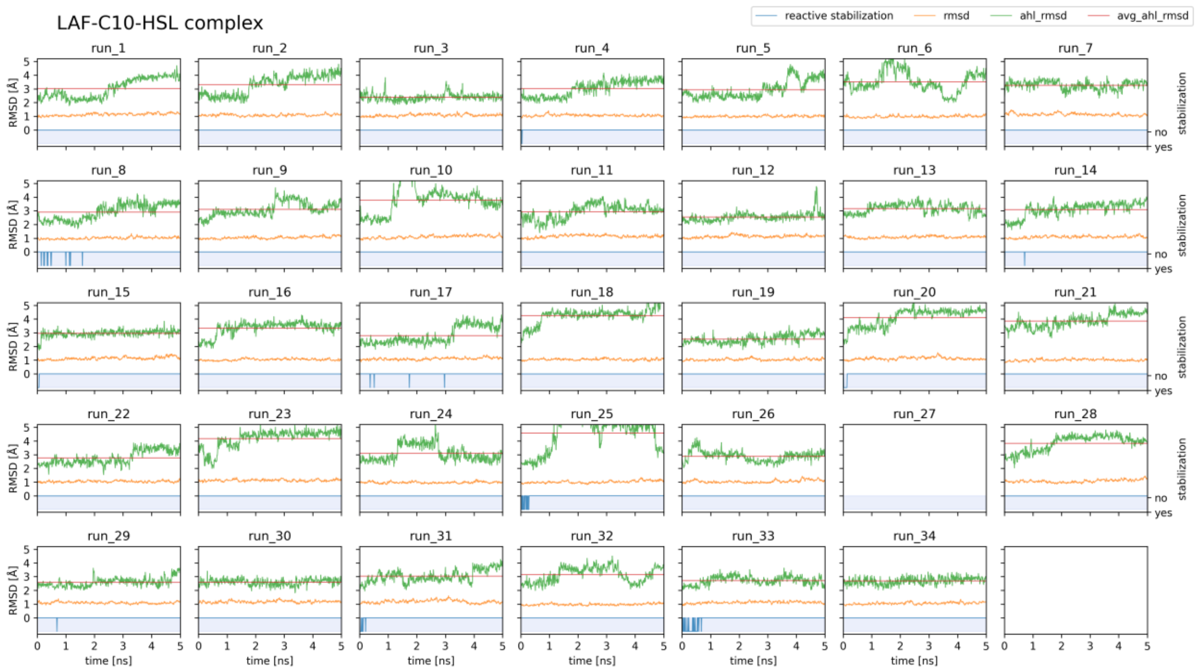
LAF-C10-HSL complex
reactive stabilization
rmsd
ahl_rmsd
avg_ahl_rmsd
run_1 run_2 run_3 run_4 run_5 run_6 run_7
run_8 run_9 run_10 run_11 run_12 run_13 run_14
run_15 run_16 run_17 run_18 run_19 run_20 run_21
run_22 run_23 run_24 run_25 run_26 run_27 run_28
run_29 run_30 run_31 run_32 run_33 run_34
RMSD [Å]
stabilization
no
yes
time [ns]

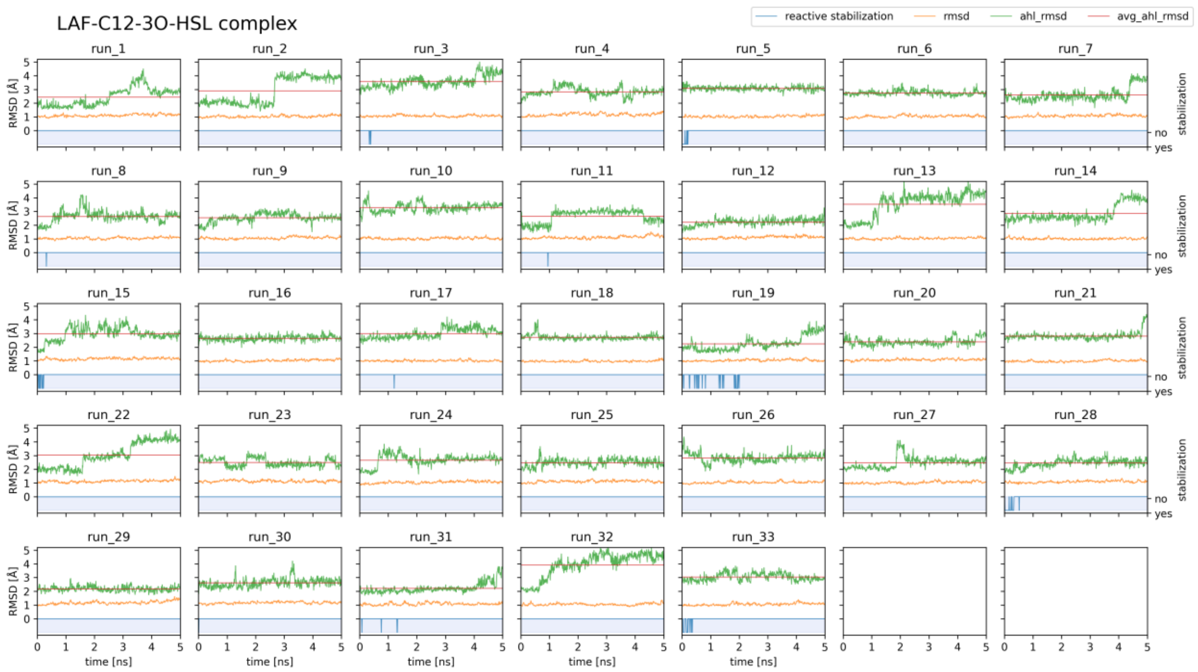
LAF-C12-3O-HSL complex
reactive stabilization
rmsd
ahl_rmsd
avg_ahl_rmsd
run_1 run_2 run_3 run_4 run_5 run_6 run_7
run_8 run_9 run_10 run_11 run_12 run_13 run_14
run_15 run_16 run_17 run_18 run_19 run_20 run_21
run_22 run_23 run_24 run_25 run_26 run_27 run_28
run_29 run_30 run_31 run_32 run_33
RMSD [Å]
stabilization
no
yes
time [ns]

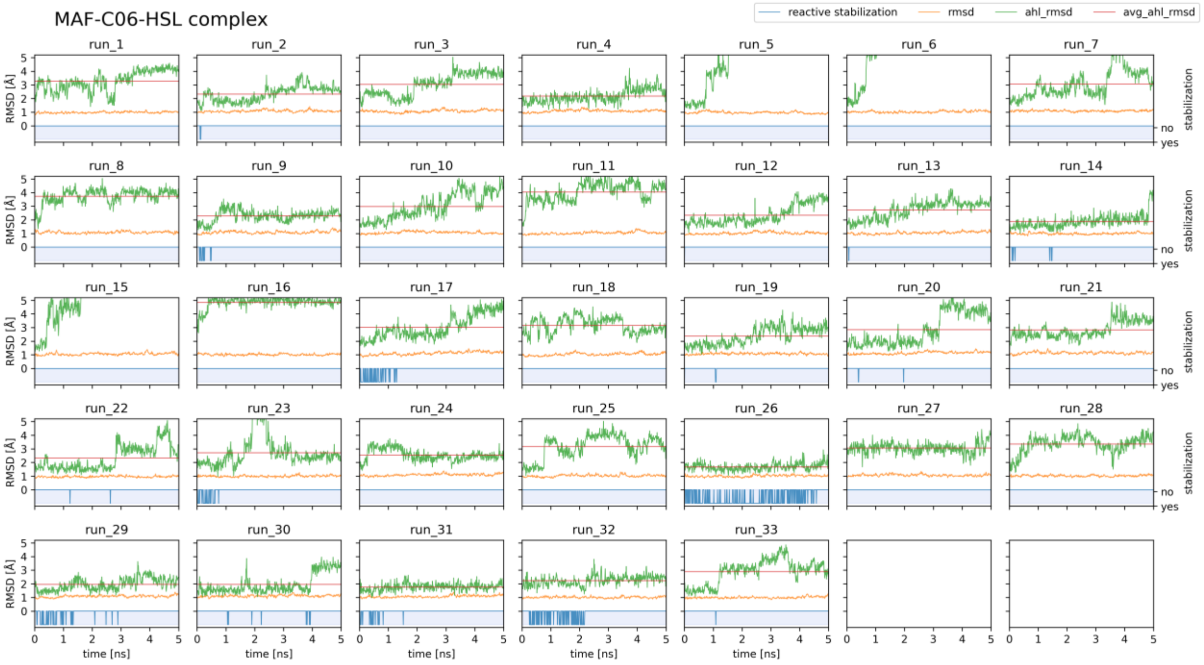
MAF-C06-HSL complex
reactive stabilization
rmsd
ahl_rmsd
avg_ahl_rmsd
run_1 run_2 run_3 run_4 run_5 run_6 run_7
run_8 run_9 run_10 run_11 run_12 run_13 run_14
run_15 run_16 run_17 run_18 run_19 run_20 run_21
run_22 run_23 run_24 run_25 run_26 run_27 run_28
run_29 run_30 run_31 run_32 run_33
RMSD [Å]
time [ns]
stabilization
no
yes

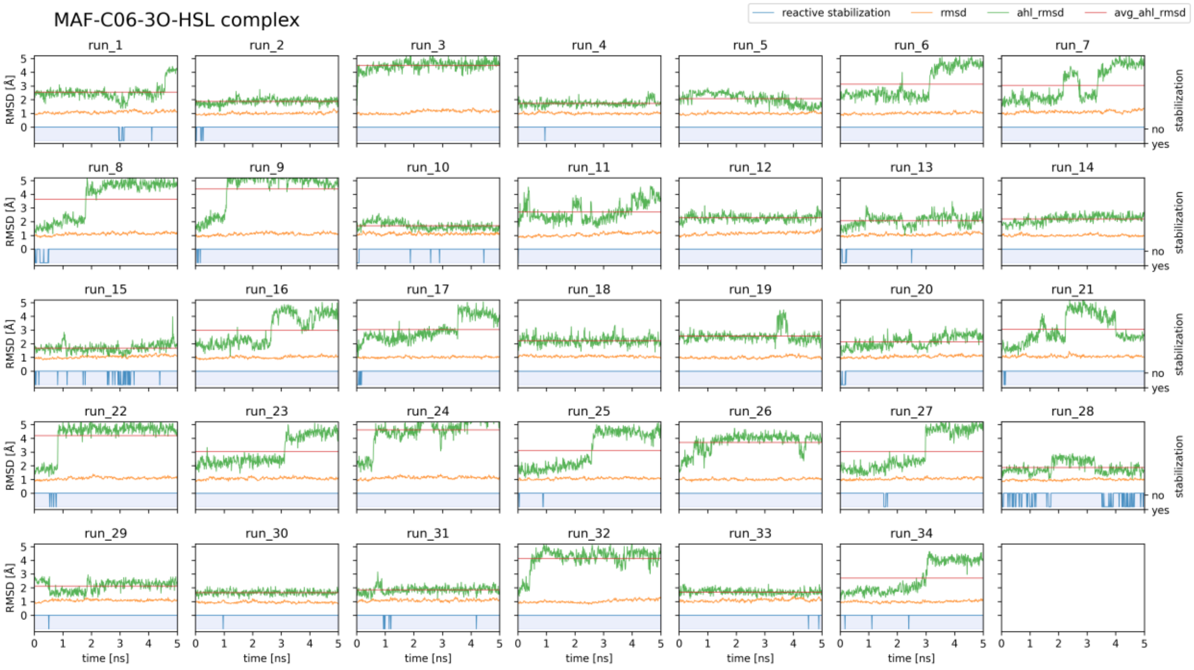
MAF-C06-3O-HSL complex
reactive stabilization
rmsd
ahl_rmsd
avg_ahl_rmsd
run_1 run_2 run_3 run_4 run_5 run_6 run_7
run_8 run_9 run_10 run_11 run_12 run_13 run_14
run_15 run_16 run_17 run_18 run_19 run_20 run_21
run_22 run_23 run_24 run_25 run_26 run_27 run_28
run_29 run_30 run_31 run_32 run_33 run_34
RMSD [Å]
time [ns]
stabilization
no
yes

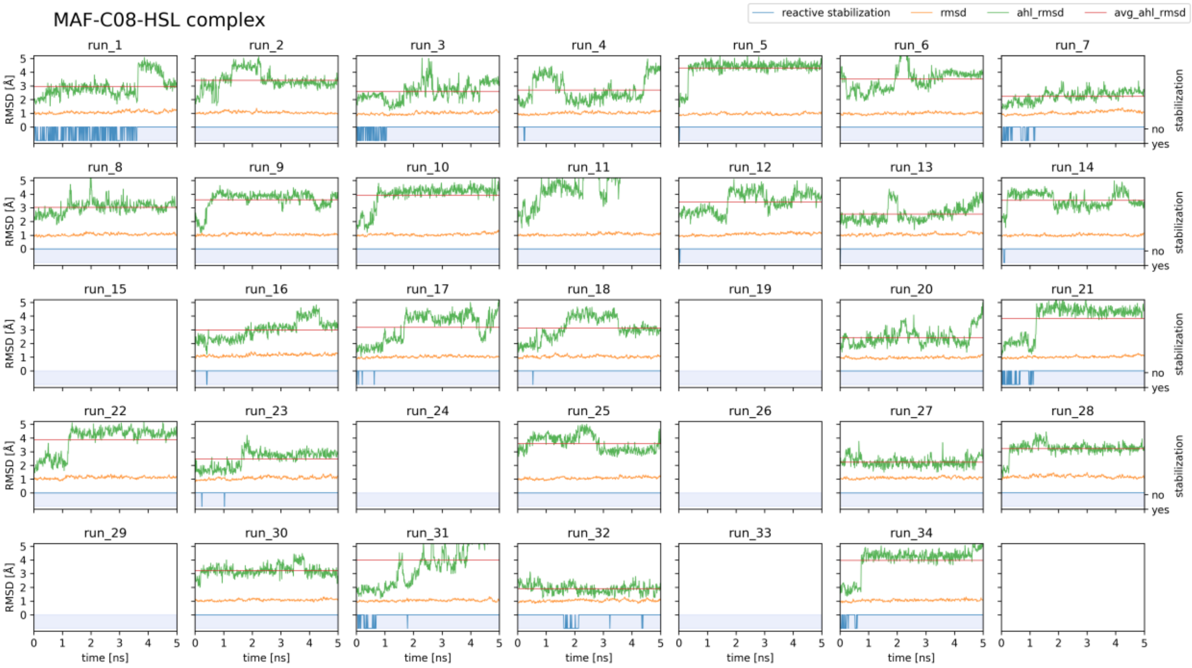
MAF-C08-HSL complex
reactive stabilization
rmsd
ahl_rmsd
avg_ahl_rmsd
RMSD [Å]
time [ns]
stabilization
no
yes

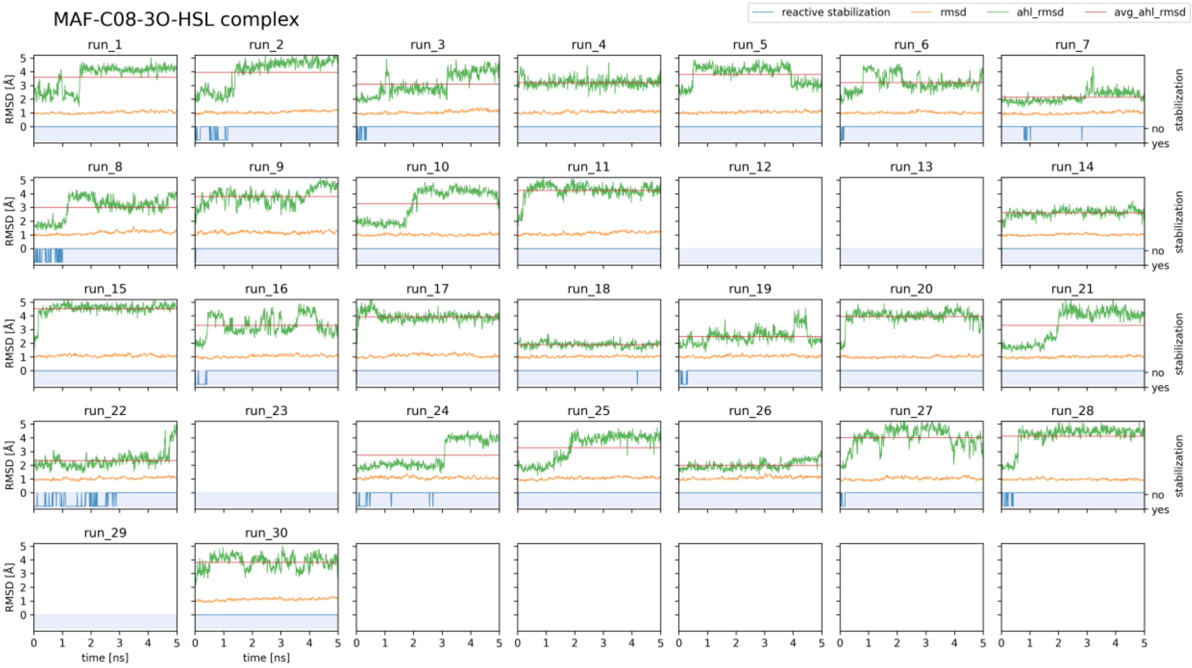
MAF-C08-3O-HSL complex
reactive stabilization
rmsd
ahl_rmsd
avg_ahl_rmsd
RMSD [Å]
time [ns]
stabilization
no
yes

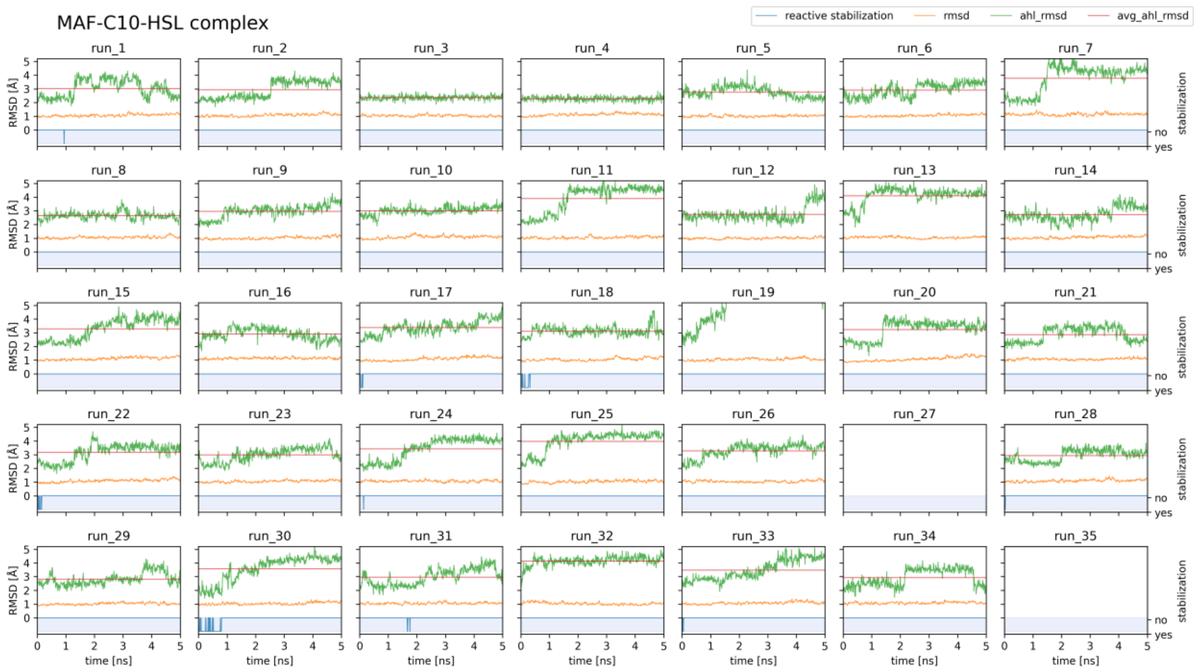
MAF-C10-HSL complex
reactive stabilization
rmsd
ahl_rmsd
avg_ahl_rmsd
RMSD [Å]
stabilization
no
yes
time [ns]

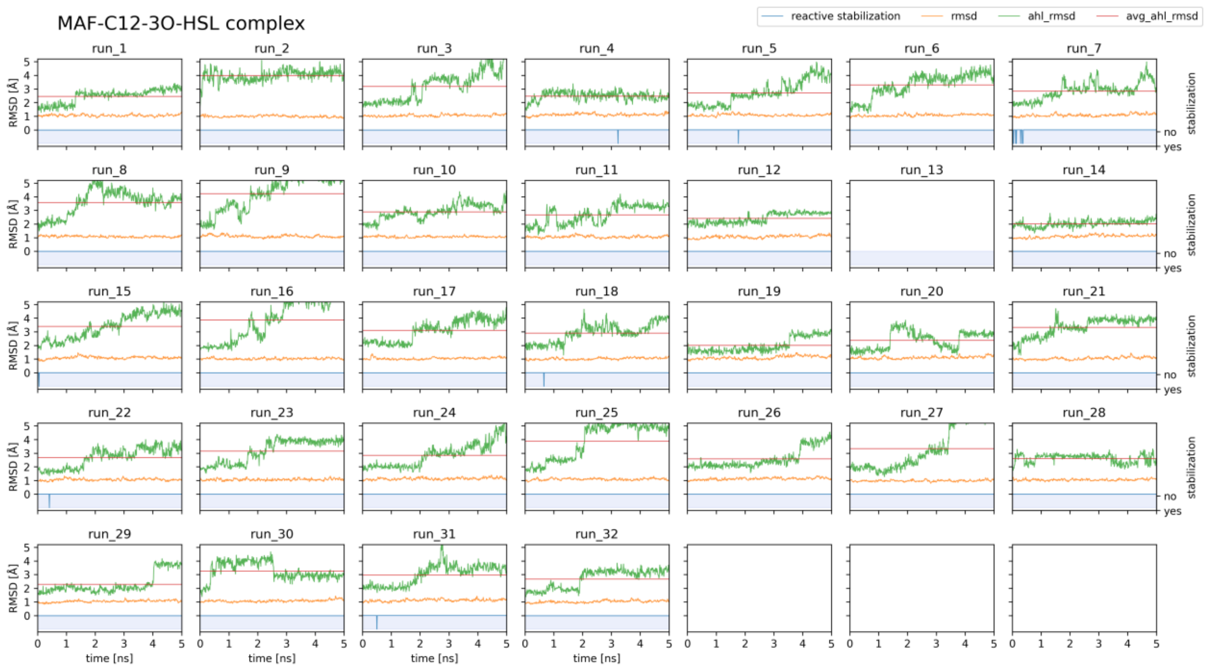
MAF-C12-3O-HSL complex
reactive stabilization
rmsd
ahl_rmsd
avg_ahl_rmsd
RMSD [Å]
stabilization
no
yes
time [ns]

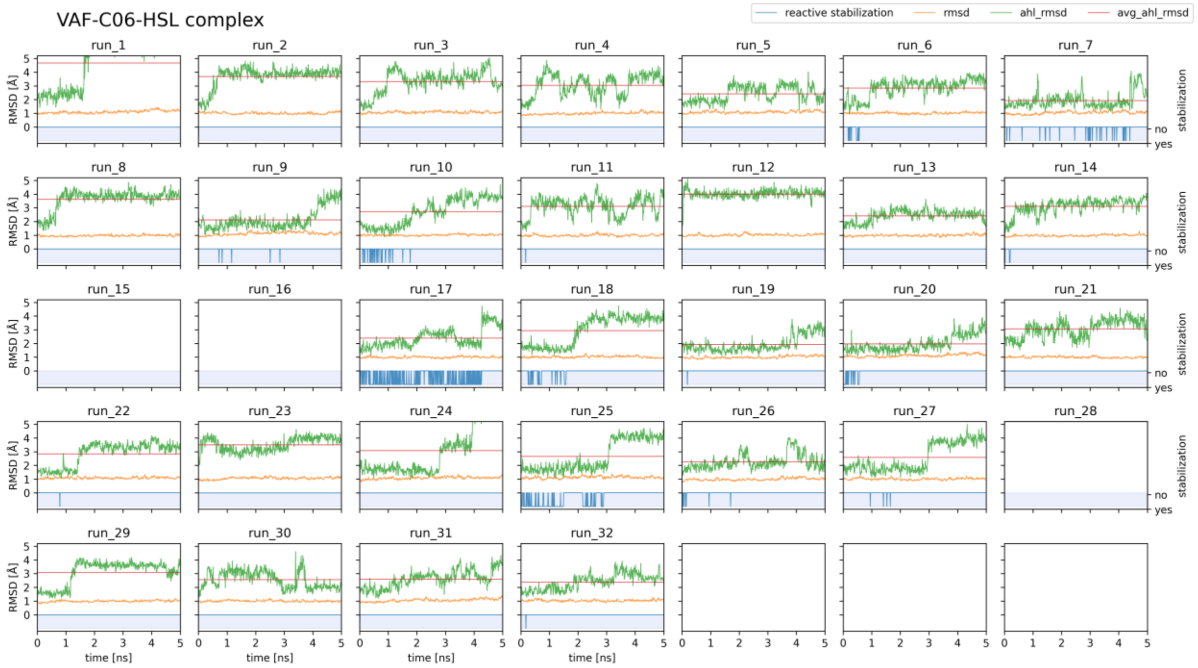
VAF-C06-HSL complex
reactive stabilization
rmsd
ahl_rmsd
avg_ahl_rmsd
run_1
run_2
run_3
run_4
run_5
run_6
run_7
run_8
run_9
run_10
run_11
run_12
run_13
run_14
run_15
run_16
run_17
run_18
run_19
run_20
run_21
run_22
run_23
run_24
run_25
run_26
run_27
run_28
run_29
run_30
run_31
run_32
RMSD [Å]
stabilization
no
yes
time [ns]

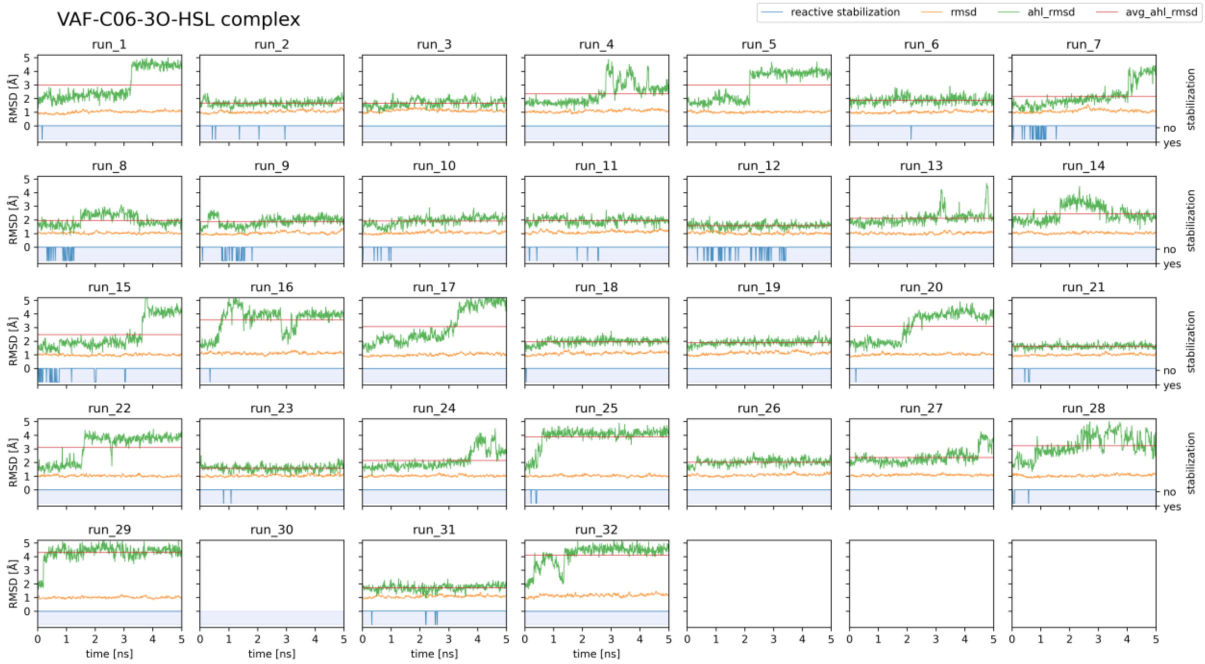
VAF-C06-3O-HSL complex
reactive stabilization
rmsd
ahl_rmsd
avg_ahl_rmsd
run_1
run_2
run_3
run_4
run_5
run_6
run_7
run_8
run_9
run_10
run_11
run_12
run_13
run_14
run_15
run_16
run_17
run_18
run_19
run_20
run_21
run_22
run_23
run_24
run_25
run_26
run_27
run_28
run_29
run_30
run_31
run_32
RMSD [Å]
stabilization
no
yes
time [ns]

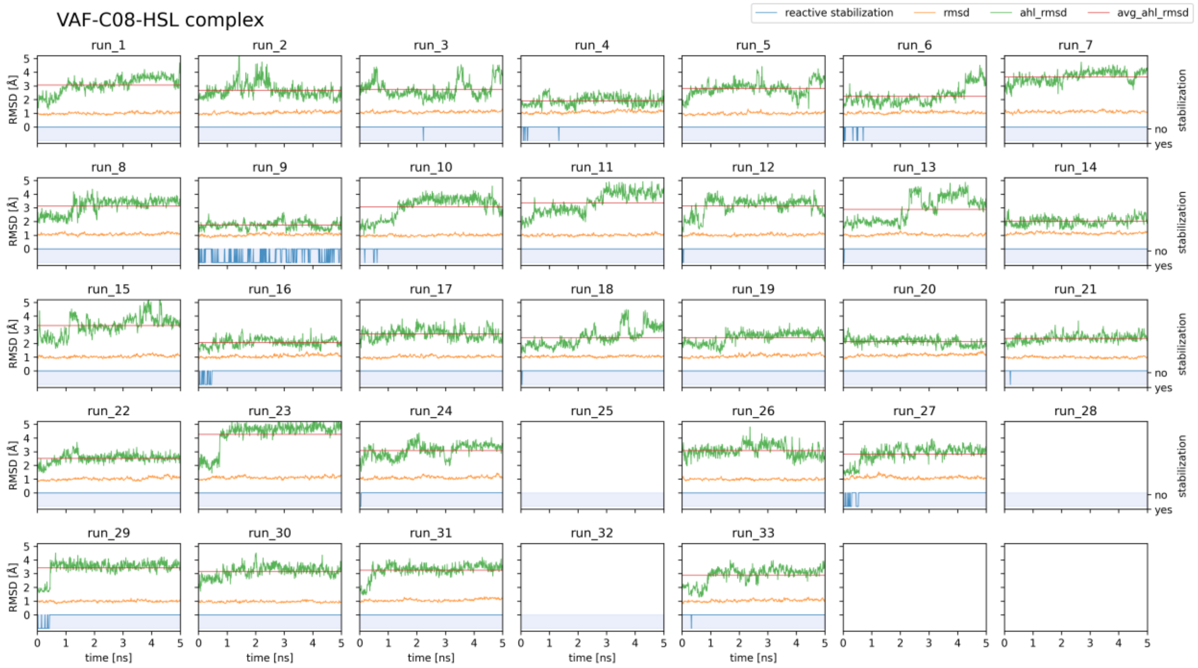
VAF-C08-HSL complex
reactive stabilization
rmsd
ahl_rmsd
avg_ahl_rmsd
run_1
run_2
run_3
run_4
run_5
run_6
run_7
run_8
run_9
run_10
run_11
run_12
run_13
run_14
run_15
run_16
run_17
run_18
run_19
run_20
run_21
run_22
run_23
run_24
run_25
run_26
run_27
run_28
run_29
run_30
run_31
run_32
run_33
RMSD [Å]
time [ns]
stabilization
no
yes

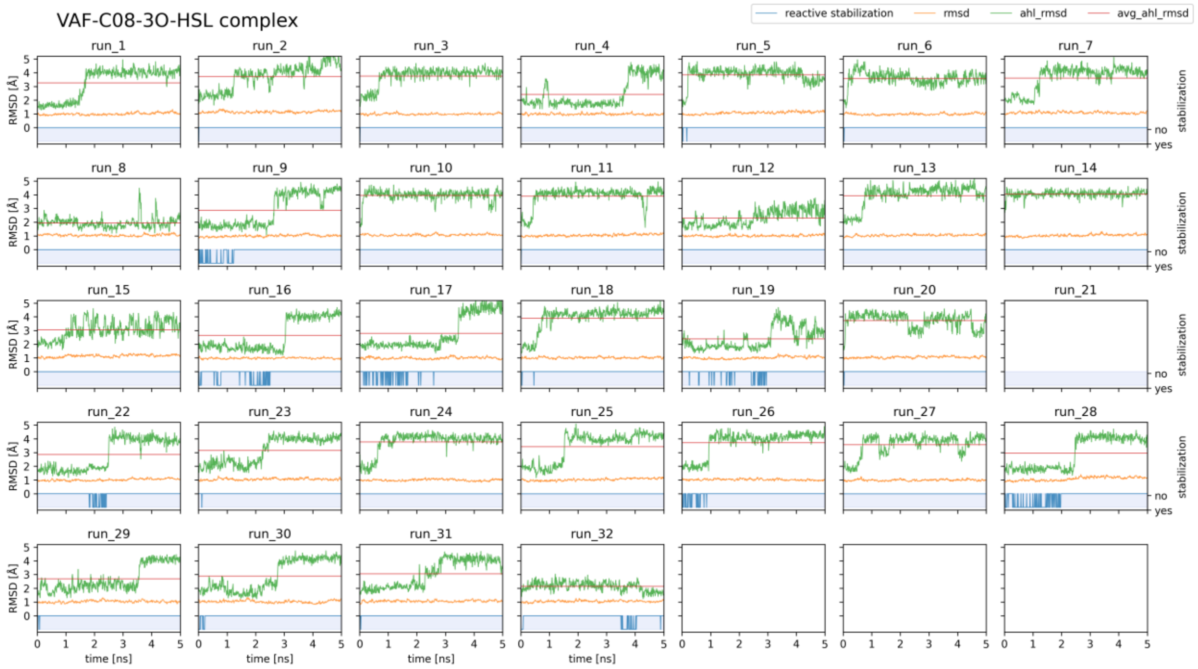
VAF-C08-3O-HSL complex
reactive stabilization
rmsd
ahl_rmsd
avg_ahl_rmsd
run_1
run_2
run_3
run_4
run_5
run_6
run_7
run_8
run_9
run_10
run_11
run_12
run_13
run_14
run_15
run_16
run_17
run_18
run_19
run_20
run_21
run_22
run_23
run_24
run_25
run_26
run_27
run_28
run_29
run_30
run_31
run_32
RMSD [Å]
time [ns]
stabilization
no
yes

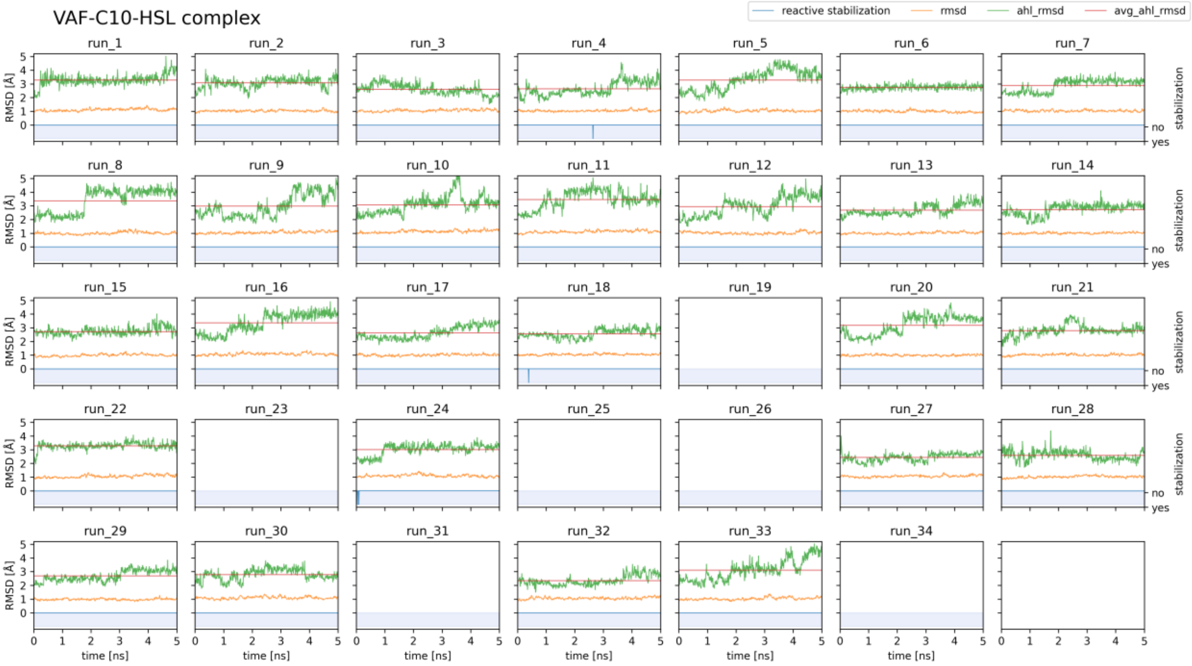
VAF-C10-HSL complex
reactive stabilization
rmsd
ahl_rmsd
avg_ahl_rmsd
RMSD [Å]
time [ns]
stabilization
no
yes

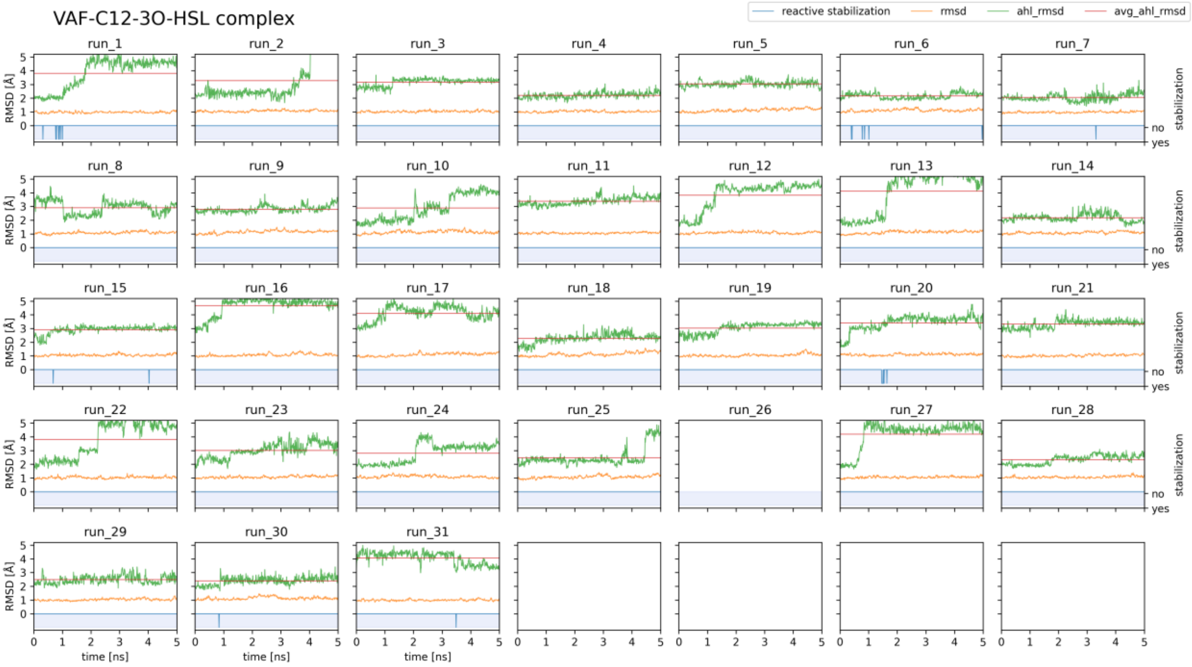
VAF-C12-3O-HSL complex
reactive stabilization
rmsd
ahl_rmsd
avg_ahl_rmsd
RMSD [Å]
time [ns]
stabilization
no
yes

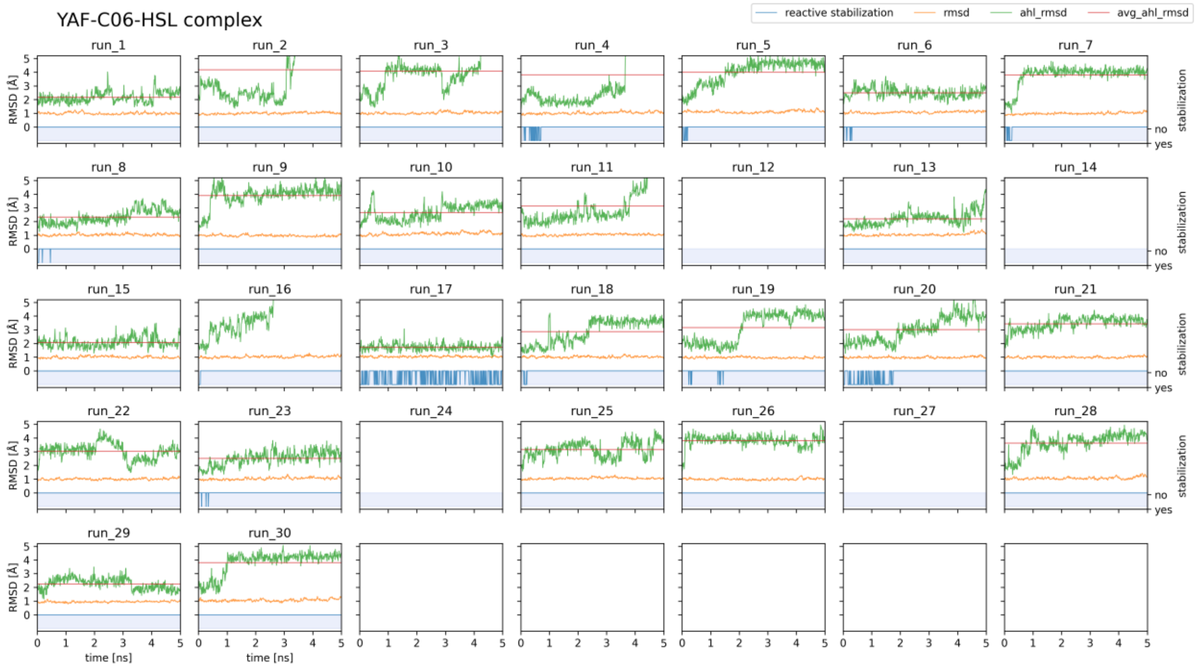
YAF-C06-HSL complex
reactive stabilization
rmsd
ahl_rmsd
avg_ahl_rmsd
run_1 run_2 run_3 run_4 run_5 run_6 run_7
run_8 run_9 run_10 run_11 run_12 run_13 run_14
run_15 run_16 run_17 run_18 run_19 run_20 run_21
run_22 run_23 run_24 run_25 run_26 run_27 run_28
run_29 run_30
RMSD [Å]
stabilization
no
yes
time [ns]

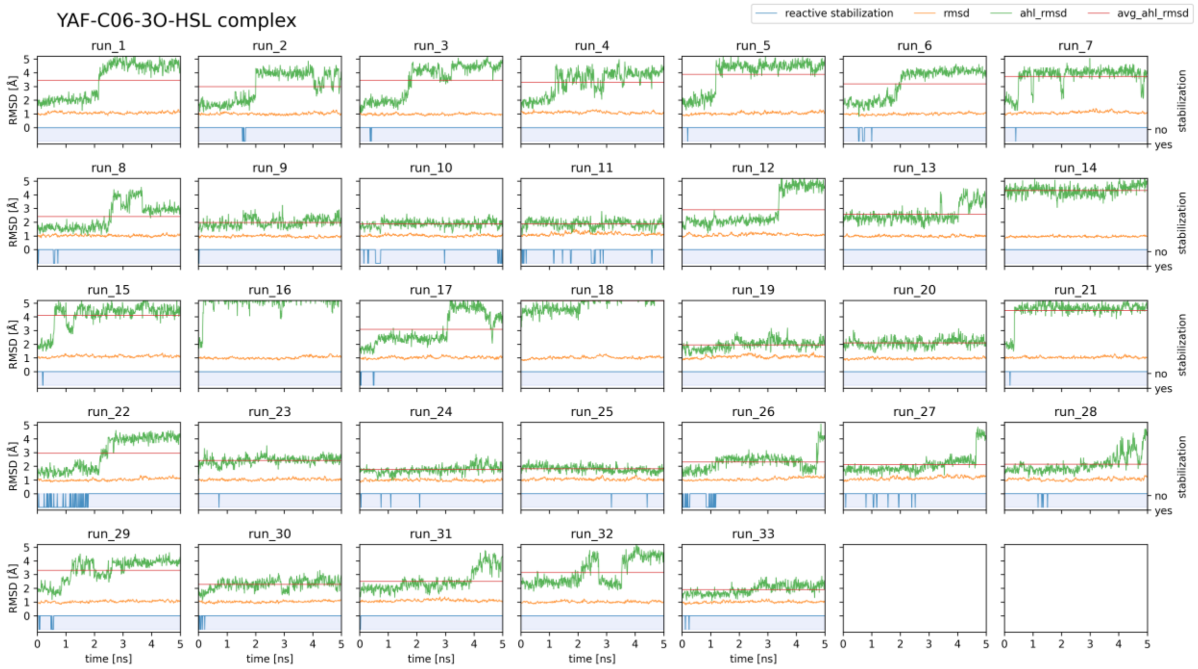
YAF-C06-3O-HSL complex
reactive stabilization
rmsd
ahl_rmsd
avg_ahl_rmsd
run_1 run_2 run_3 run_4 run_5 run_6 run_7
run_8 run_9 run_10 run_11 run_12 run_13 run_14
run_15 run_16 run_17 run_18 run_19 run_20 run_21
run_22 run_23 run_24 run_25 run_26 run_27 run_28
run_29 run_30 run_31 run_32 run_33
RMSD [Å]
stabilization
no
yes
time [ns]

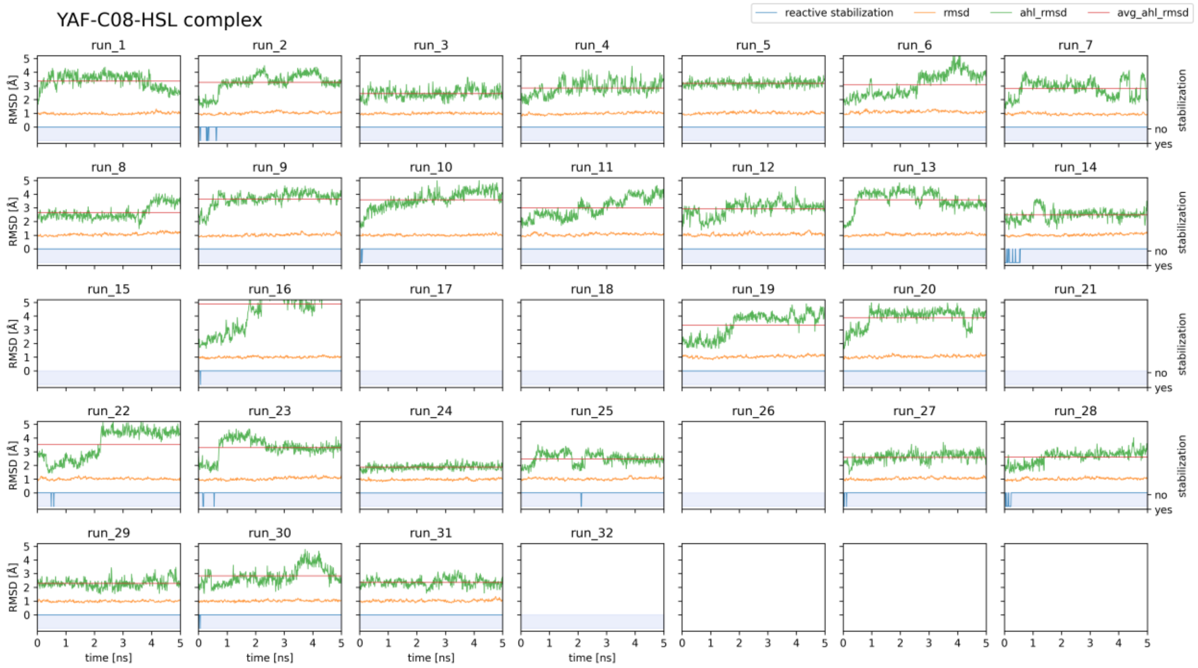
YAF-C08-HSL complex
reactive stabilization
rmsd
ahl_rmsd
avg_ahl_rmsd
RMSD [Å]
time [ns]
stabilization
no
yes

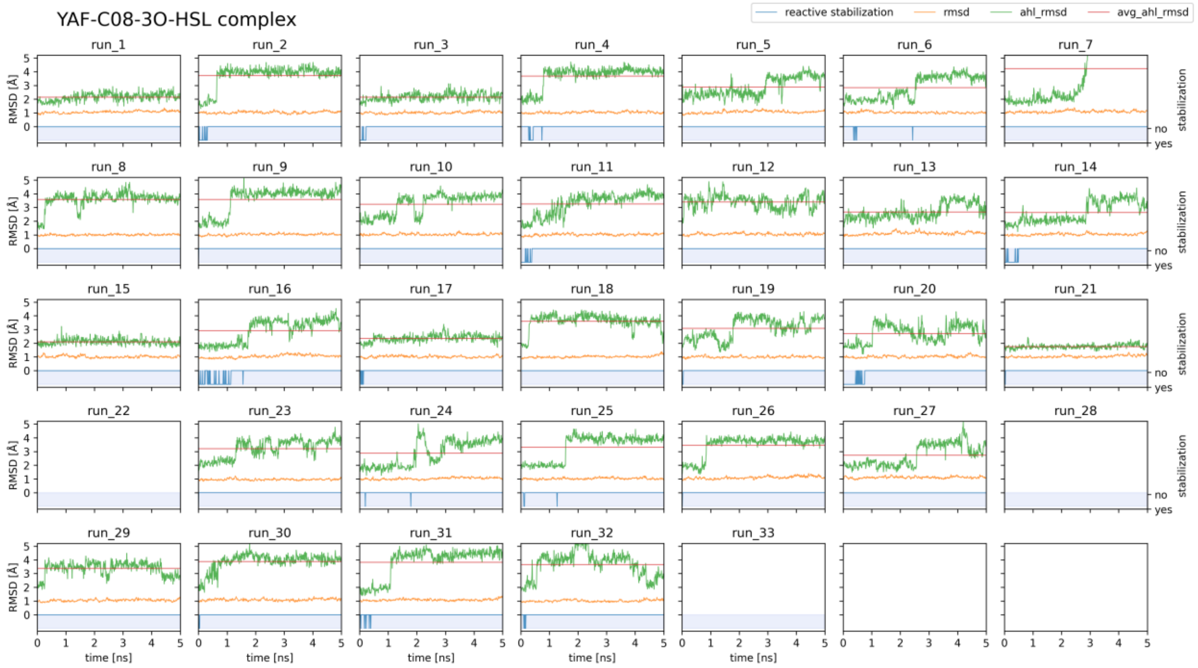
YAF-C08-3O-HSL complex
reactive stabilization
rmsd
ahl_rmsd
avg_ahl_rmsd
RMSD [Å]
time [ns]
stabilization
no
yes

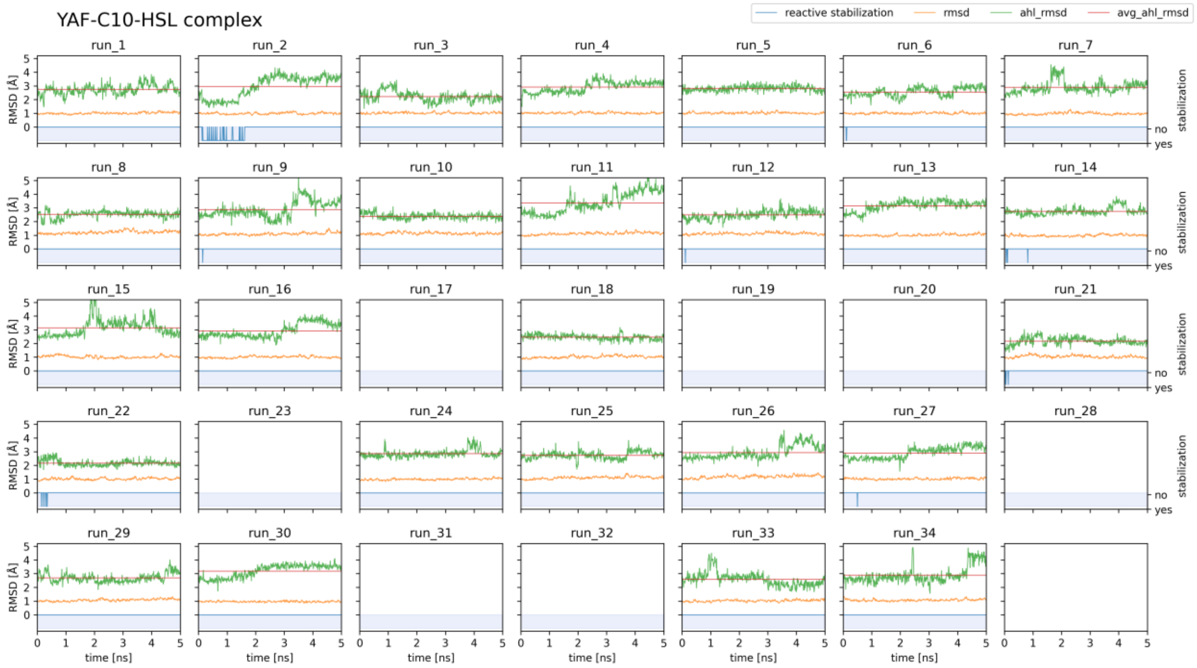
YAF-C10-HSL complex
reactive stabilization
rmsd
ahl_rmsd
avg_ahl_rmsd
RMSD [Å]
stabilization
time [ns]

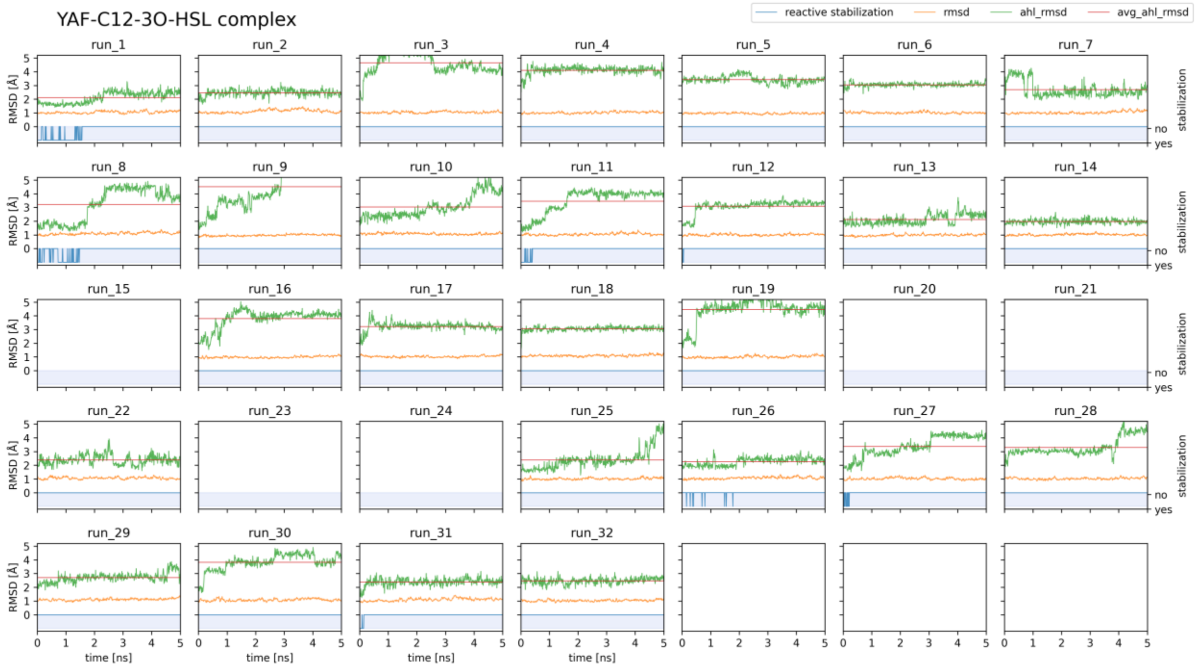
YAF-C12-3O-HSL complex
reactive stabilization
rmsd
ahl_rmsd
avg_ahl_rmsd
RMSD [Å]
stabilization
time [ns]

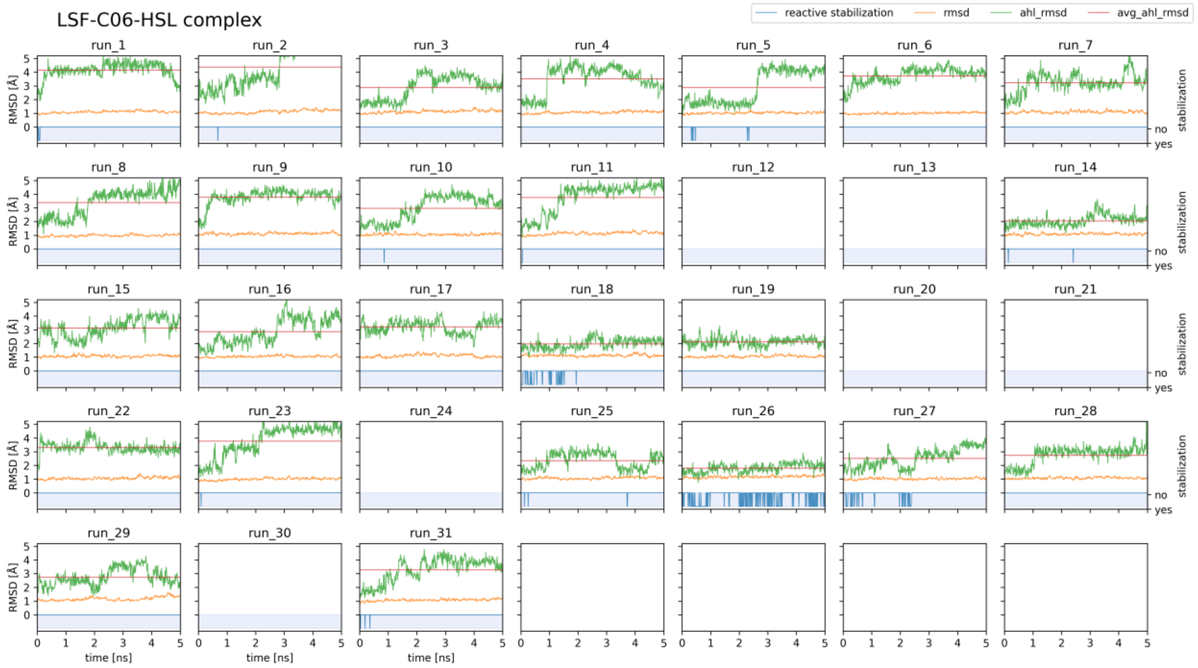
LSF-C06-HSL complex
reactive stabilization
rmsd
ahl_rmsd
avg_ahl_rmsd
run_1
run_2
run_3
run_4
run_5
run_6
run_7
run_8
run_9
run_10
run_11
run_12
run_13
run_14
run_15
run_16
run_17
run_18
run_19
run_20
run_21
run_22
run_23
run_24
run_25
run_26
run_27
run_28
run_29
run_30
run_31
RMSD [Å]
stabilization
no
yes
time [ns]

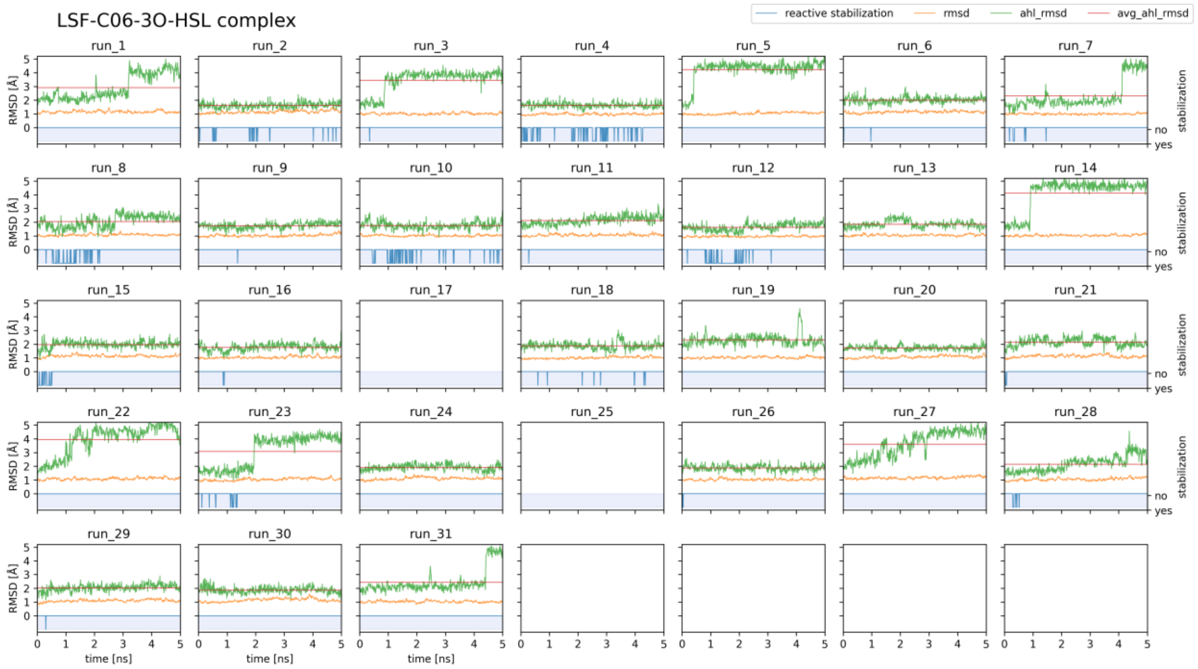
LSF-C06-3O-HSL complex
reactive stabilization
rmsd
ahl_rmsd
avg_ahl_rmsd
run_1
run_2
run_3
run_4
run_5
run_6
run_7
run_8
run_9
run_10
run_11
run_12
run_13
run_14
run_15
run_16
run_17
run_18
run_19
run_20
run_21
run_22
run_23
run_24
run_25
run_26
run_27
run_28
run_29
run_30
run_31
RMSD [Å]
stabilization
no
yes
time [ns]

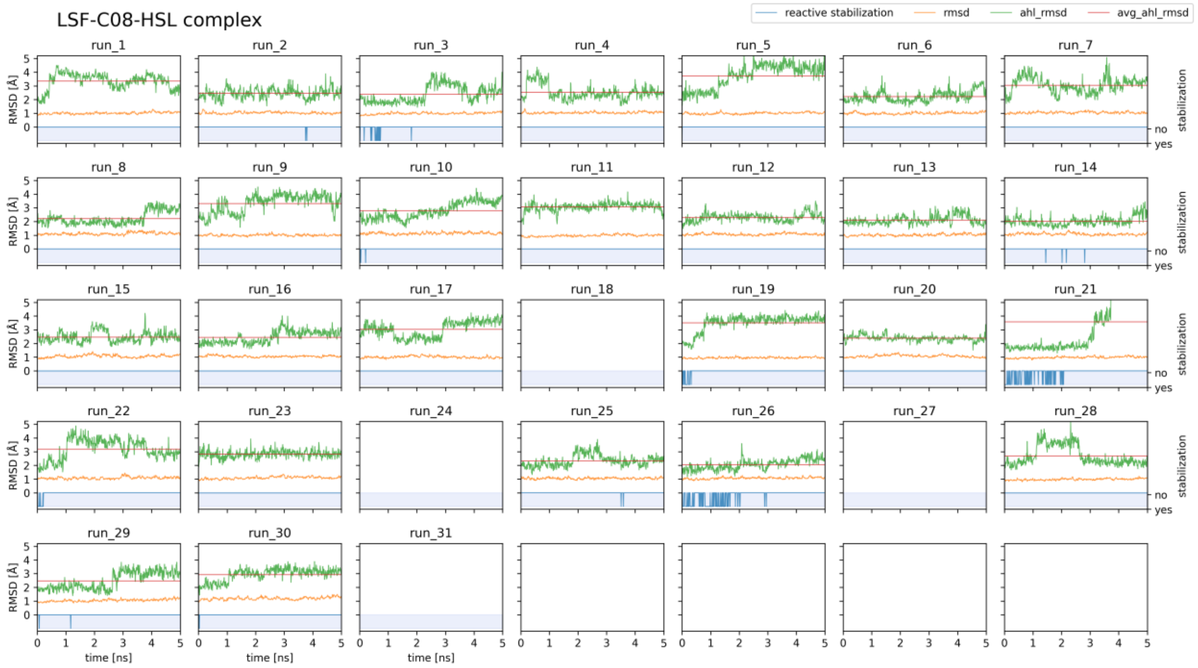
LSF-C08-HSL complex
reactive stabilization
rmsd
ahl_rmsd
avg_ahl_rmsd
run_1
run_2
run_3
run_4
run_5
run_6
run_7
run_8
run_9
run_10
run_11
run_12
run_13
run_14
run_15
run_16
run_17
run_18
run_19
run_20
run_21
run_22
run_23
run_24
run_25
run_26
run_27
run_28
run_29
run_30
run_31
RMSD [Å]
stabilization
no
yes
time [ns]

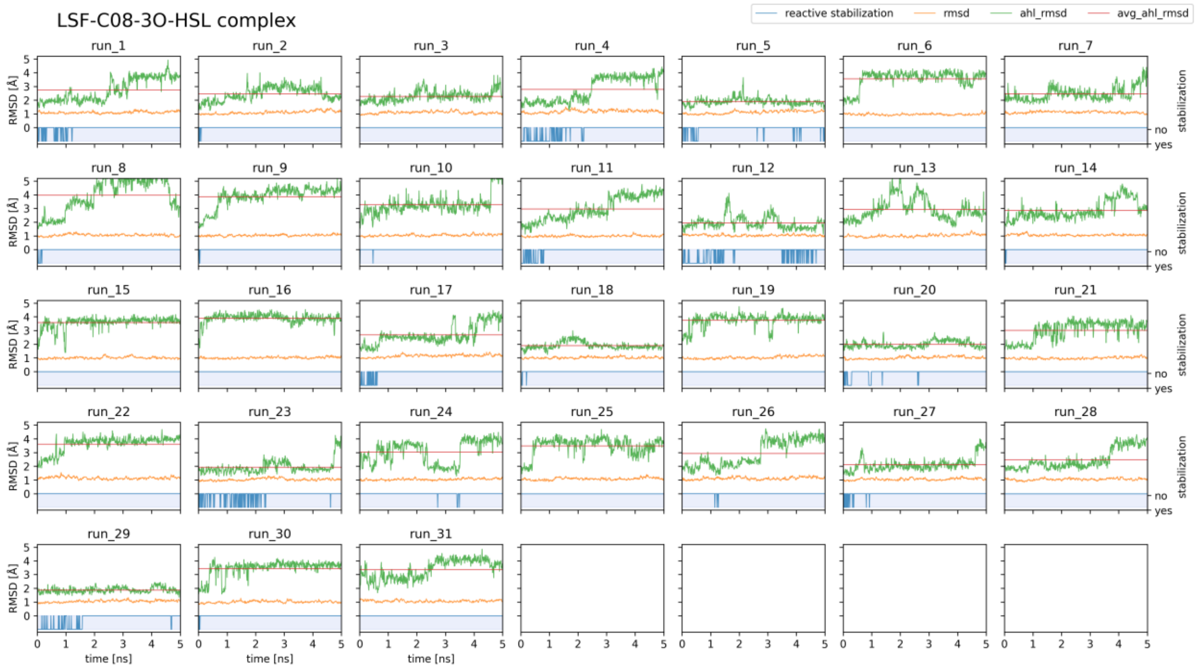
LSF-C08-3O-HSL complex
reactive stabilization
rmsd
ahl_rmsd
avg_ahl_rmsd
run_1
run_2
run_3
run_4
run_5
run_6
run_7
run_8
run_9
run_10
run_11
run_12
run_13
run_14
run_15
run_16
run_17
run_18
run_19
run_20
run_21
run_22
run_23
run_24
run_25
run_26
run_27
run_28
run_29
run_30
run_31
RMSD [Å]
stabilization
no
yes
time [ns]

LSF-C10-HSL complex

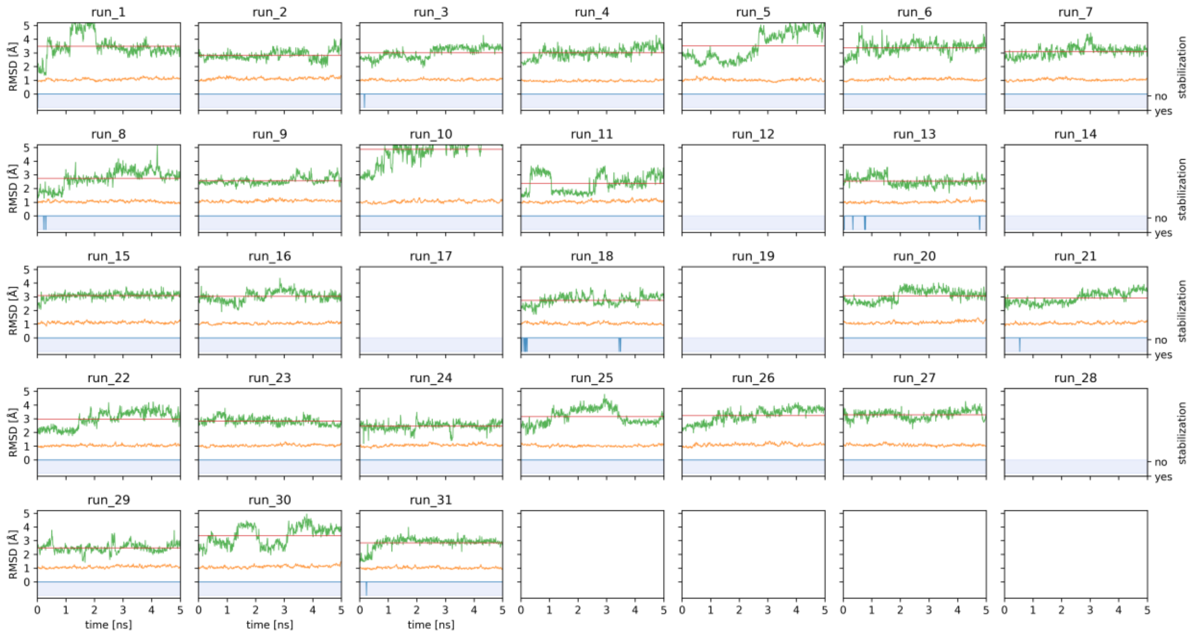


LSF-C12-3O-HSL complex

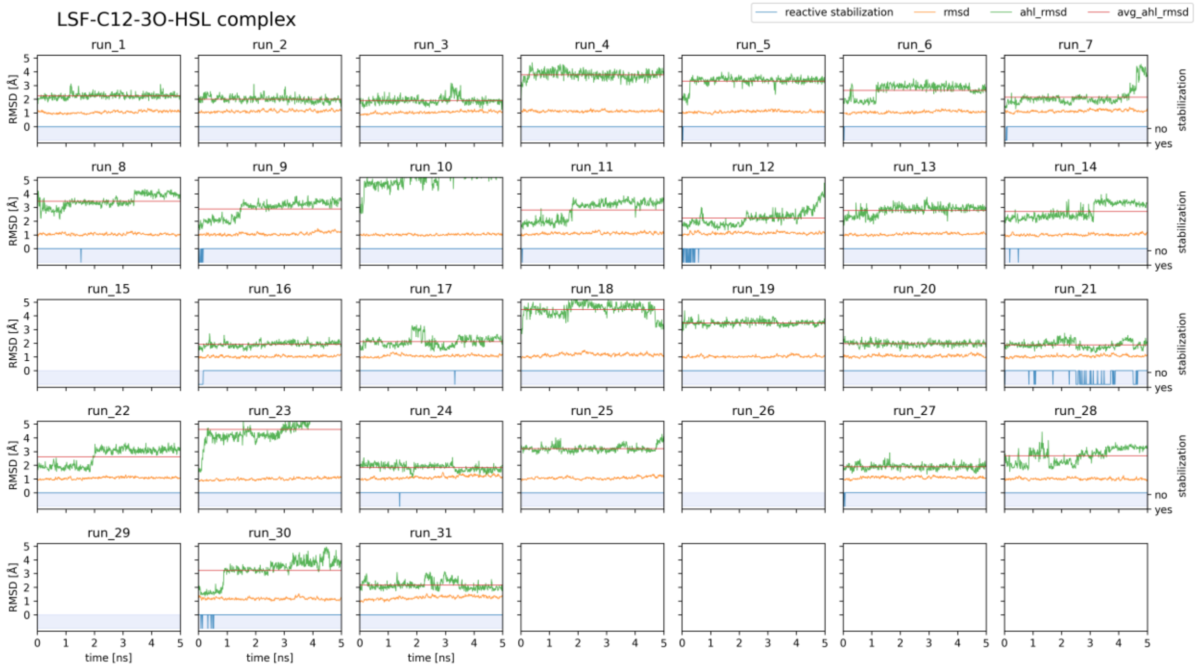

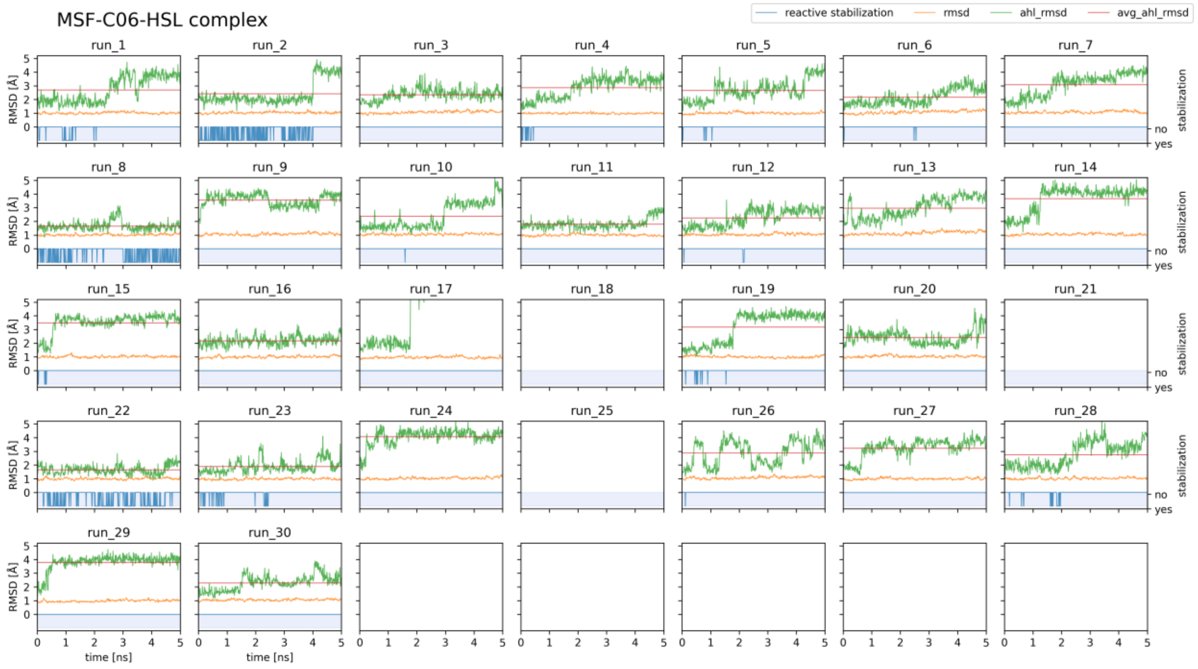
MSF-C06-HSL complex
reactive stabilization
rmsd
ahl_rmsd
avg_ahl_rmsd
run_1 run_2 run_3 run_4 run_5 run_6 run_7
run_8 run_9 run_10 run_11 run_12 run_13 run_14
run_15 run_16 run_17 run_18 run_19 run_20 run_21
run_22 run_23 run_24 run_25 run_26 run_27 run_28
run_29 run_30
RMSD [Å]
stabilization
no
yes
time [ns]

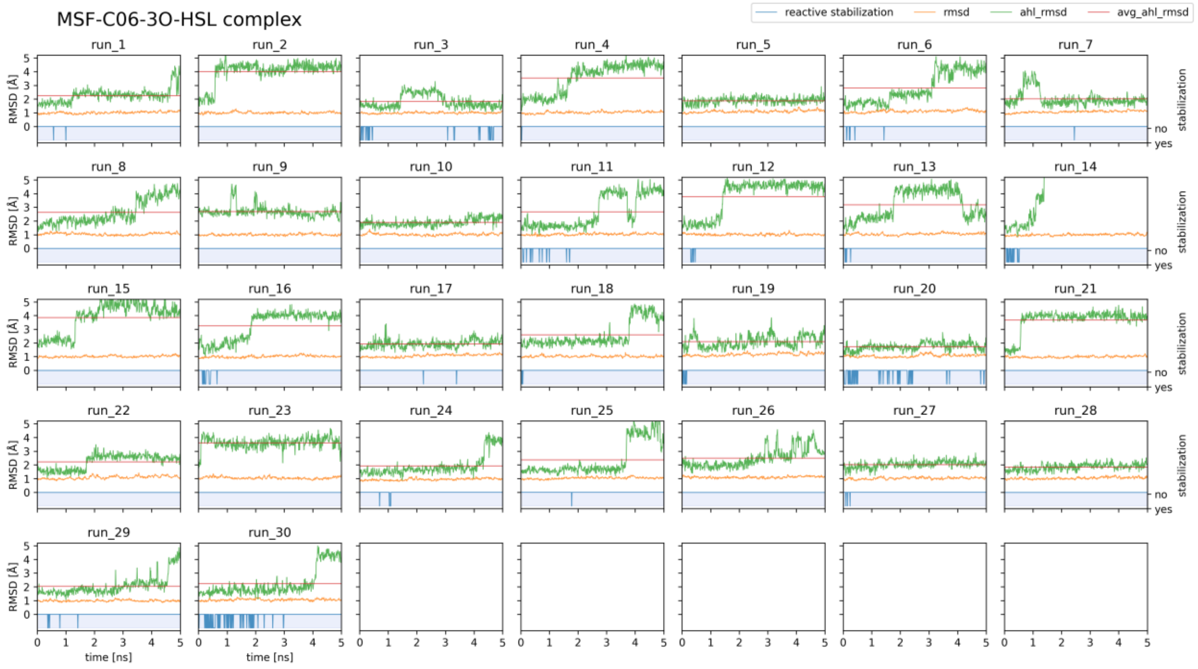
MSF-C06-3O-HSL complex
reactive stabilization
rmsd
ahl_rmsd
avg_ahl_rmsd
run_1 run_2 run_3 run_4 run_5 run_6 run_7
run_8 run_9 run_10 run_11 run_12 run_13 run_14
run_15 run_16 run_17 run_18 run_19 run_20 run_21
run_22 run_23 run_24 run_25 run_26 run_27 run_28
run_29 run_30
RMSD [Å]
stabilization
no
yes
time [ns]

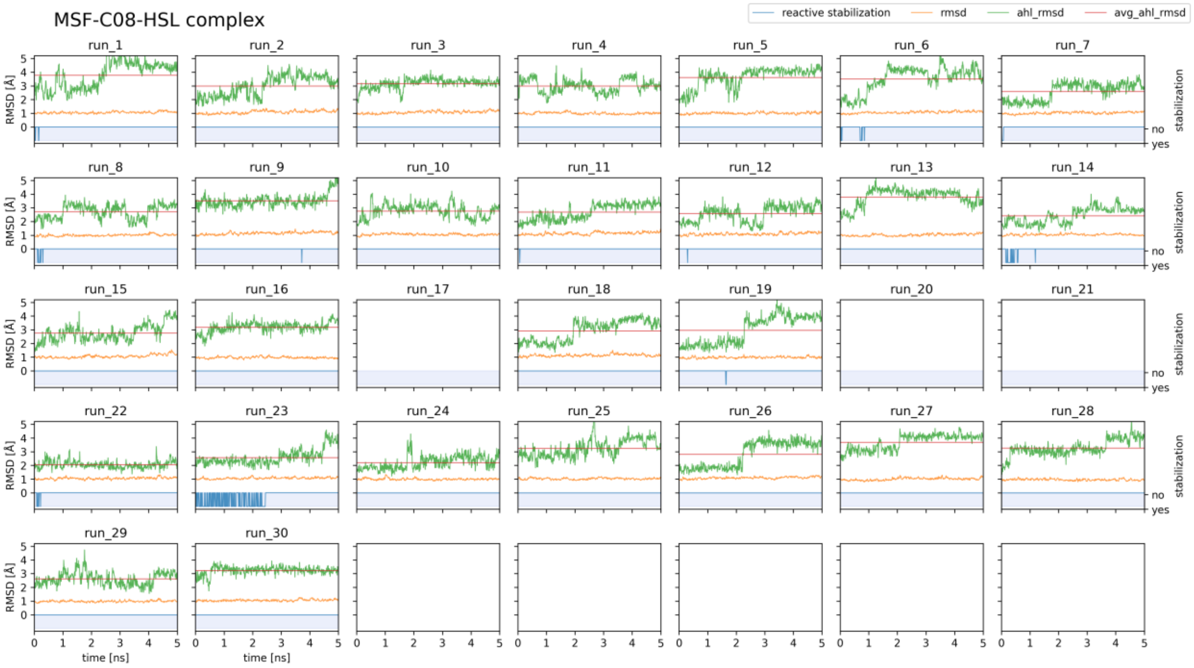
MSF-C08-HSL complex
reactive stabilization
rmsd
ahl_rmsd
avg_ahl_rmsd
RMSD [Å]
time [ns]
stabilization
no
yes

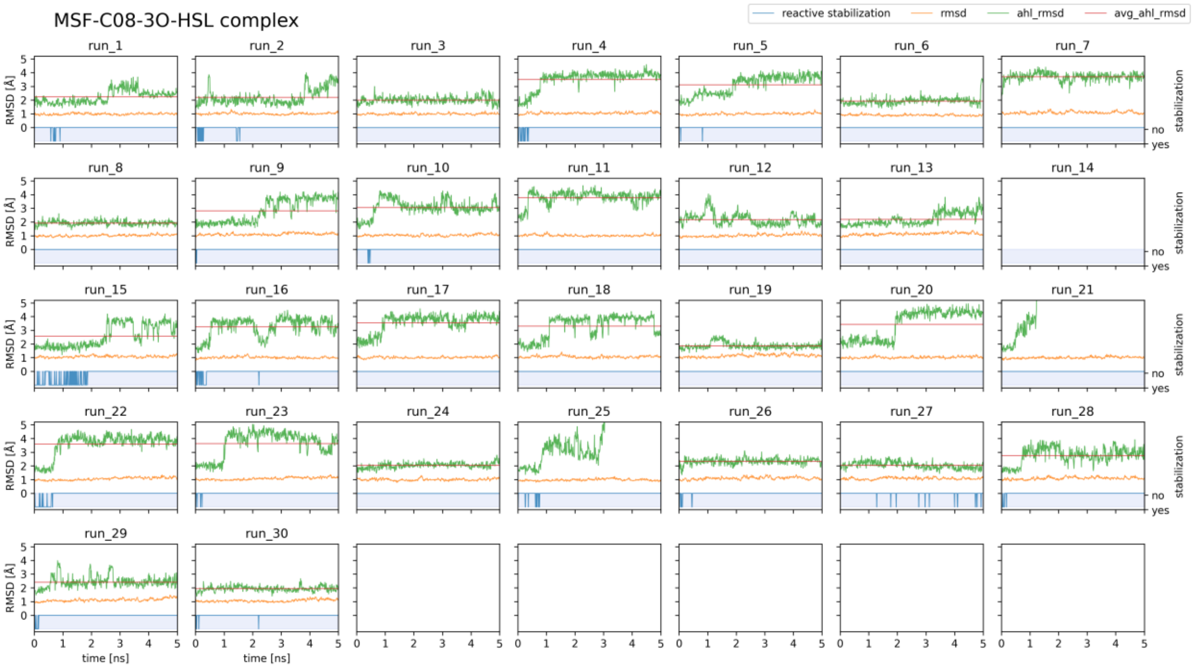
MSF-C08-3O-HSL complex
reactive stabilization
rmsd
ahl_rmsd
avg_ahl_rmsd
RMSD [Å]
time [ns]
stabilization
no
yes

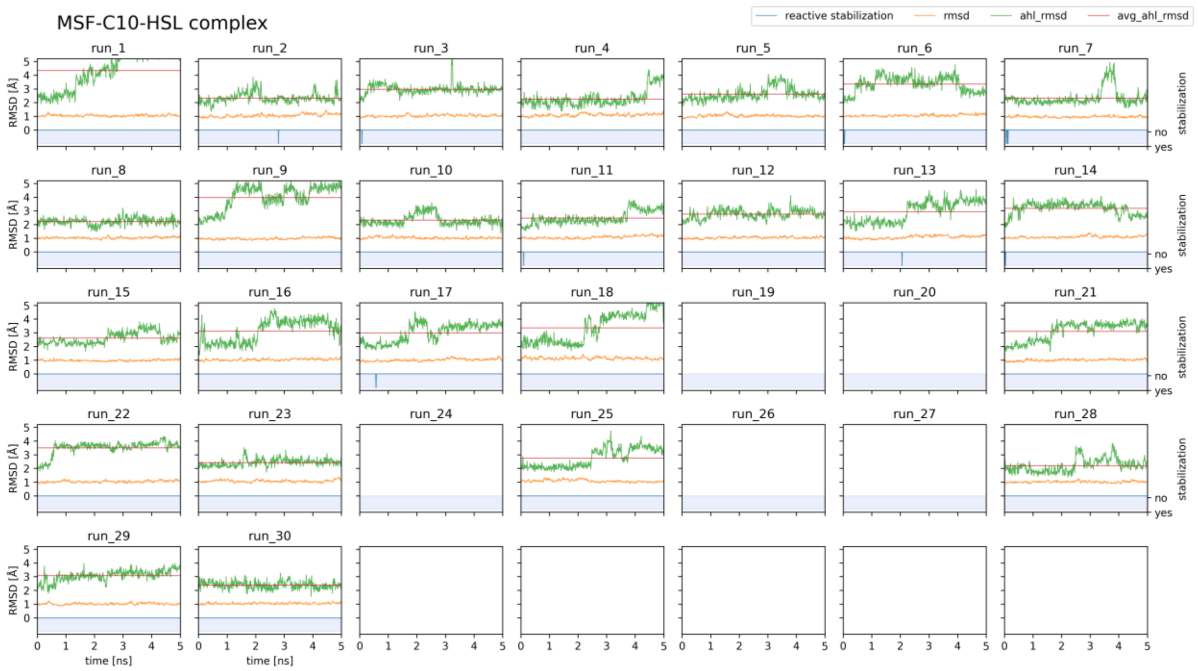
MSF-C10-HSL complex
reactive stabilization
rmsd
ahl_rmsd
avg_ahl_rmsd
run_1
run_2
run_3
run_4
run_5
run_6
run_7
run_8
run_9
run_10
run_11
run_12
run_13
run_14
run_15
run_16
run_17
run_18
run_19
run_20
run_21
run_22
run_23
run_24
run_25
run_26
run_27
run_28
run_29
run_30
RMSD [Å]
stabilization
no
yes
time [ns]

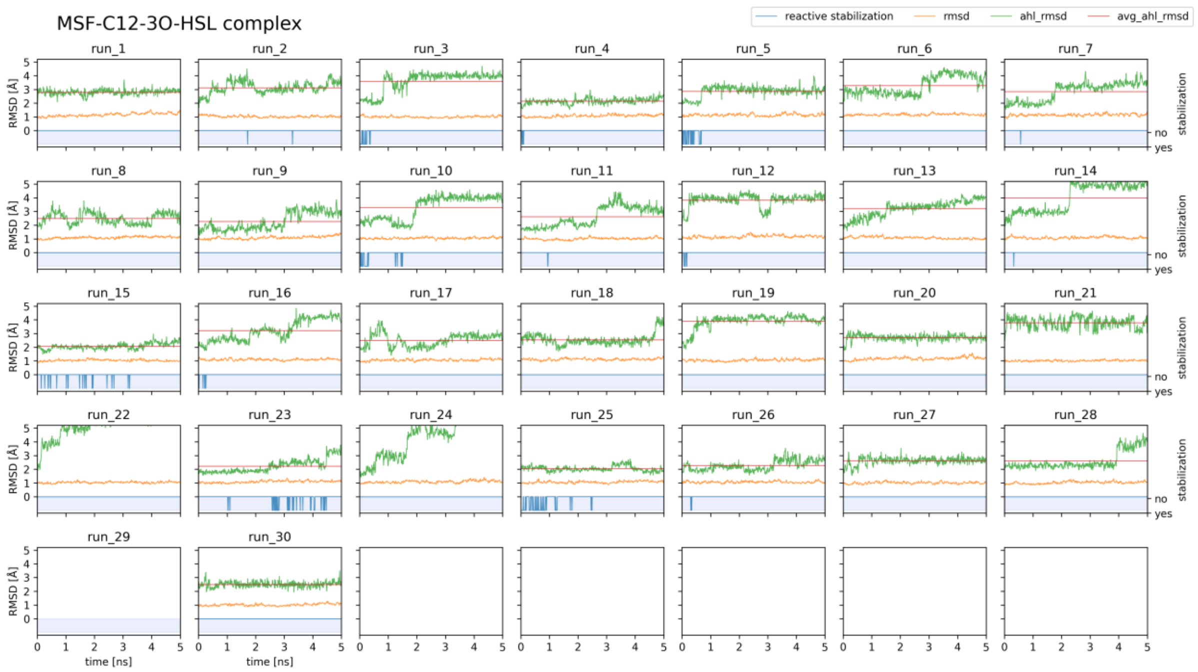
MSF-C12-3O-HSL complex
reactive stabilization
rmsd
ahl_rmsd
avg_ahl_rmsd
run_1
run_2
run_3
run_4
run_5
run_6
run_7
run_8
run_9
run_10
run_11
run_12
run_13
run_14
run_15
run_16
run_17
run_18
run_19
run_20
run_21
run_22
run_23
run_24
run_25
run_26
run_27
run_28
run_29
run_30
RMSD [Å]
stabilization
no
yes
time [ns]

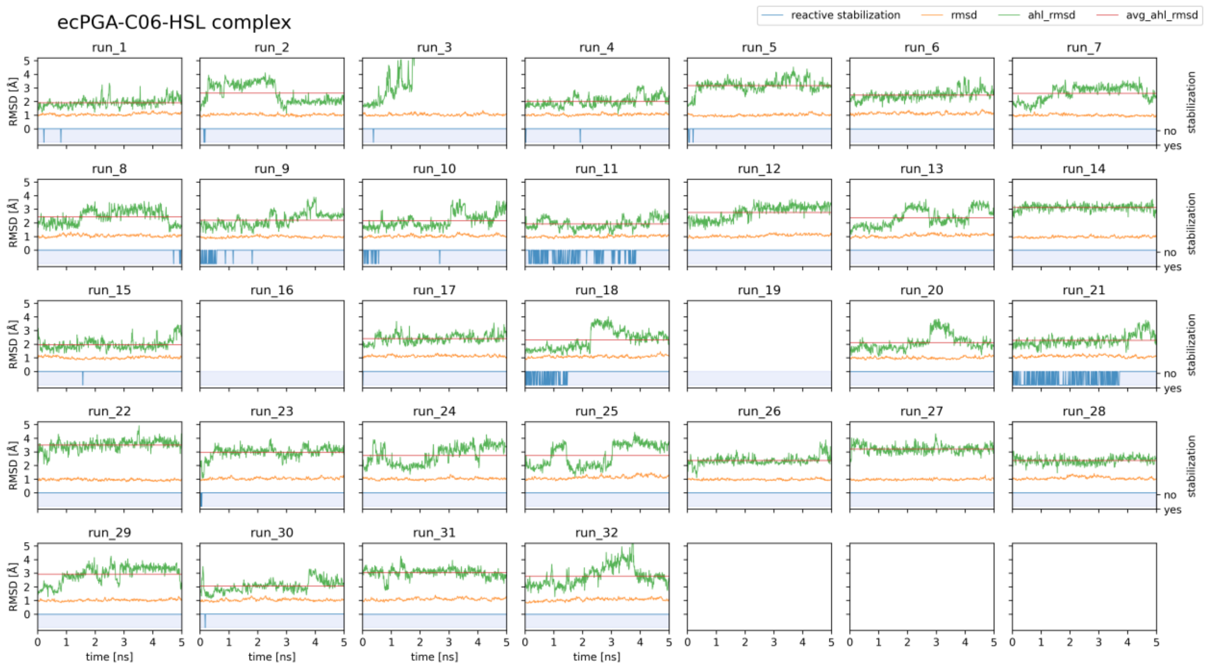
ecPGA-C06-HSL complex
reactive stabilization
rmsd
ahl_rmsd
avg_ahl_rmsd
RMSD [Å]
stabilization
no
yes
time [ns]

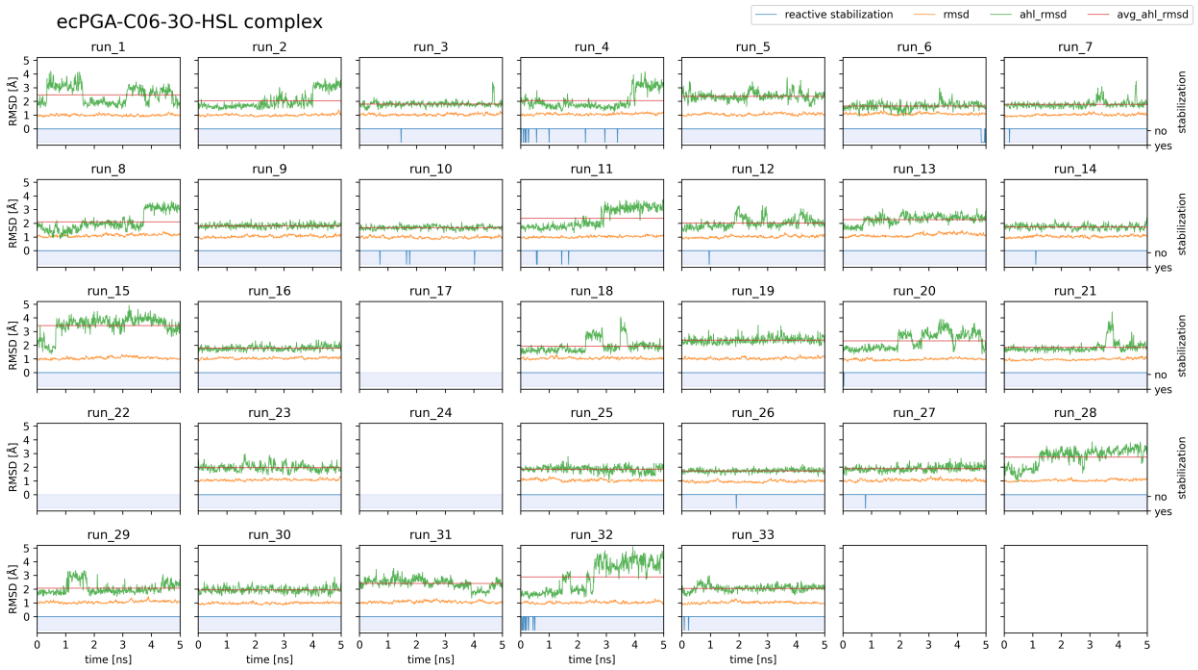
ecPGA-C06-3O-HSL complex
reactive stabilization
rmsd
ahl_rmsd
avg_ahl_rmsd
RMSD [Å]
stabilization
no
yes
time [ns]

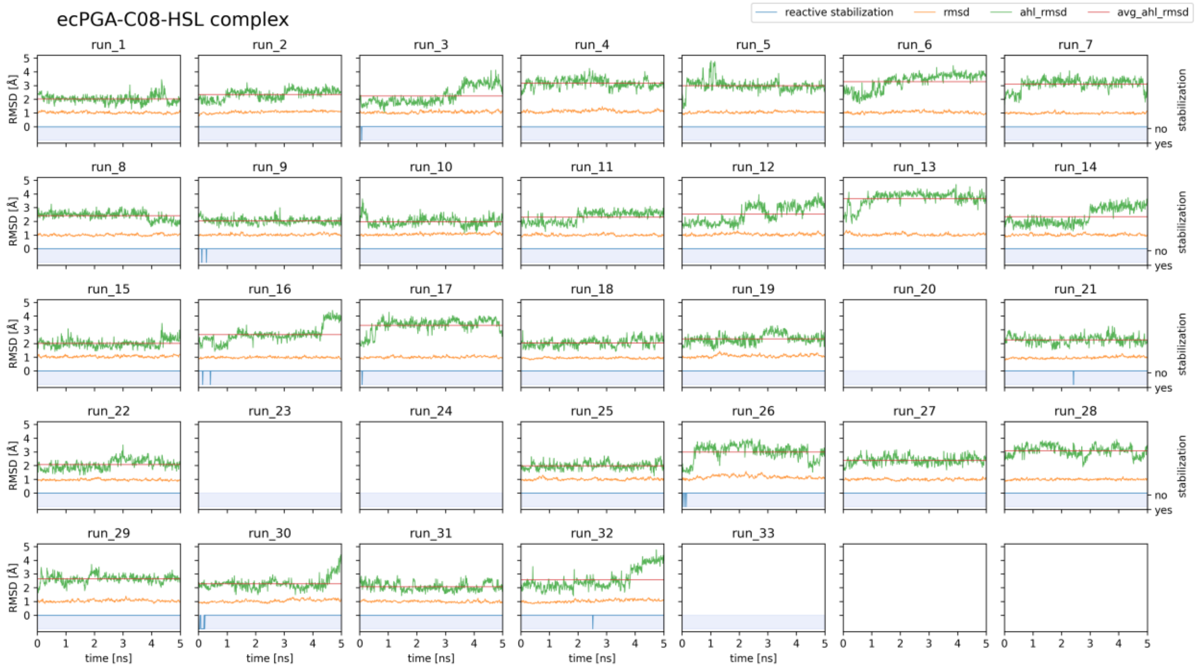
ecPGA-C08-HSL complex
reactive stabilization
rmsd
ahl_rmsd
avg_ahl_rmsd
RMSD [Å]
time [ns]
stabilization
no
yes

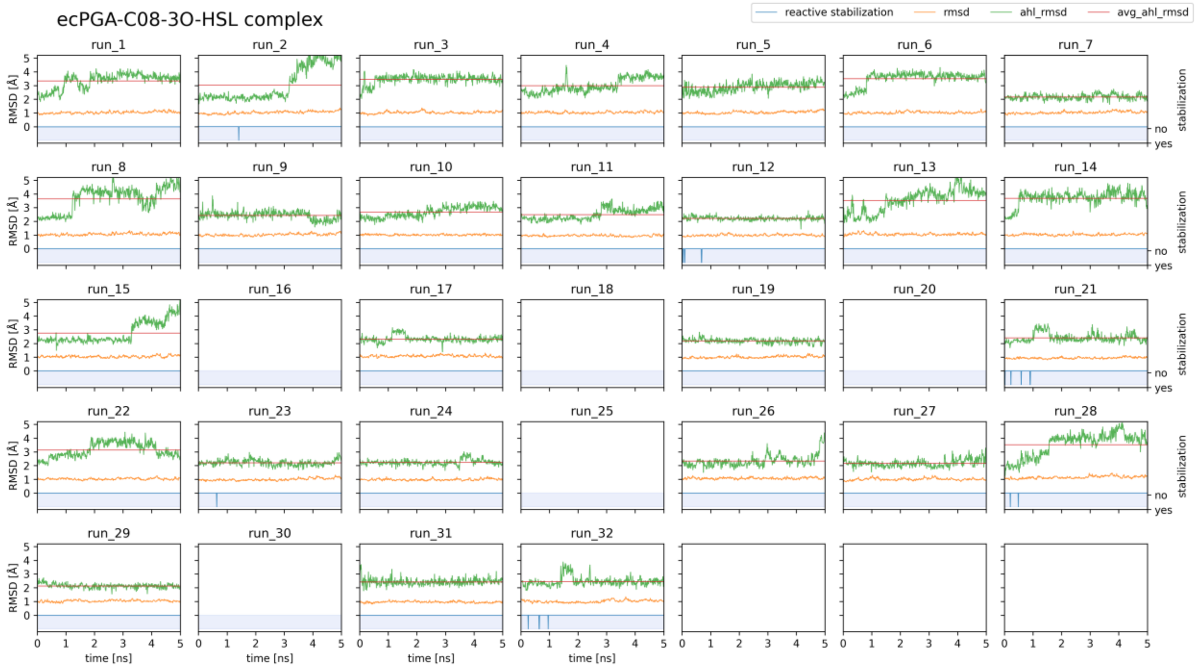
ecPGA-C08-3O-HSL complex
reactive stabilization
rmsd
ahl_rmsd
avg_ahl_rmsd
RMSD [Å]
time [ns]
stabilization
no
yes

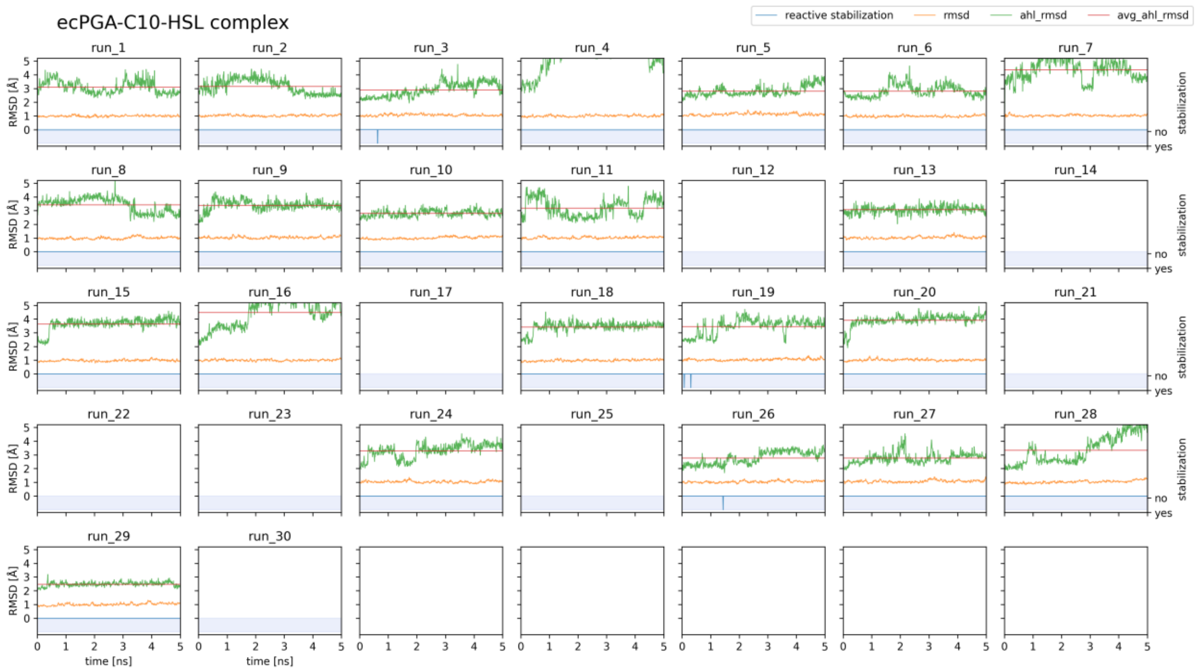
ecPGA-C10-HSL complex
reactive stabilization
rmsd
ahl_rmsd
avg_ahl_rmsd
RMSD [Å]
stabilization
no
yes
time [ns]

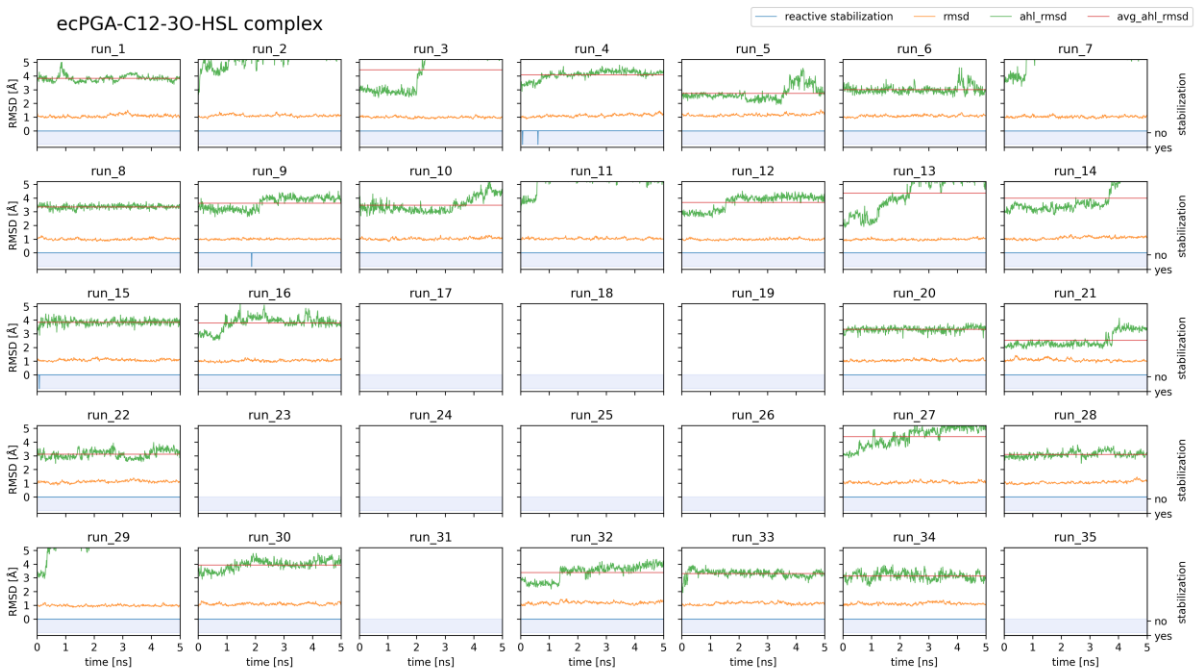
ecPGA-C12-3O-HSL complex
reactive stabilization
rmsd
ahl_rmsd
avg_ahl_rmsd
RMSD [Å]
stabilization
no
yes
time [ns]
