## Supplementary Information File 3 for "Engineering dynamic gates in binding pocket of penicillin G acylase to selectively degrade bacterial signaling molecules"

‡ These authors contributed equally.

This supplementary file includes ecPGA variants activity as a function of substrate concentration for conversion of N-acyl-homoserine lactons (AHLs).

Full names of presented protein variants:

LAF, Phe138αLeu & Met142αAla & Ile177βPhe ecPGA variant;

MAF, Phe138αMet & Met142αAla & Ile177βPhe ecPGA variant;

VAF, Phe138αVal & Met142αAla & Ile177βPhe ecPGA variant;

YAF, Phe138αTyr & Met142αAla & Ile177βPhe ecPGA variant;

LSF, Phe138αLeu & Met142αSer & Ile177βPhe ecPGA variant;

MSF, Phe138αMet & Met142αSer & Ile177βPhe ecPGA variant

Full names of presented AHLs:

C06-HSL, N-hexanoyl-L-homoserine lactone;

C06-3O-HSL, N-3-oxo-hexanoyl-L-homoserine lactone;

C08-HSL, N-octanoyl-L-homoserine lactone;

C08-3O-HSL, N-3-oxo-octanoyl-L-homoserine lactone;

C10-HSL, N-decanoyl-L-homoserine lactone;

C12-3O-HSL, N-3-oxo-dodecanoyl-L-homoserine lactone

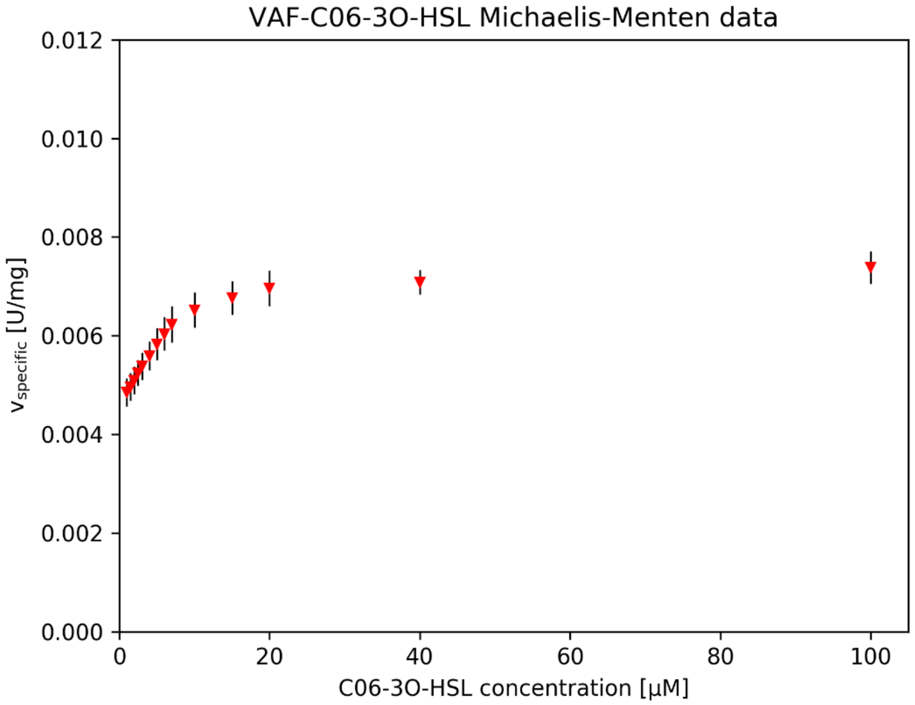

VAF-C06-3O-HSL Michaelis-Menten data
0.012
0.010
0.008
0.006
0.004
0.002
0.000
0
20
40
60
80
100
$v_{specific}$ [U/mg]
C06-3O-HSL concentration [µM]


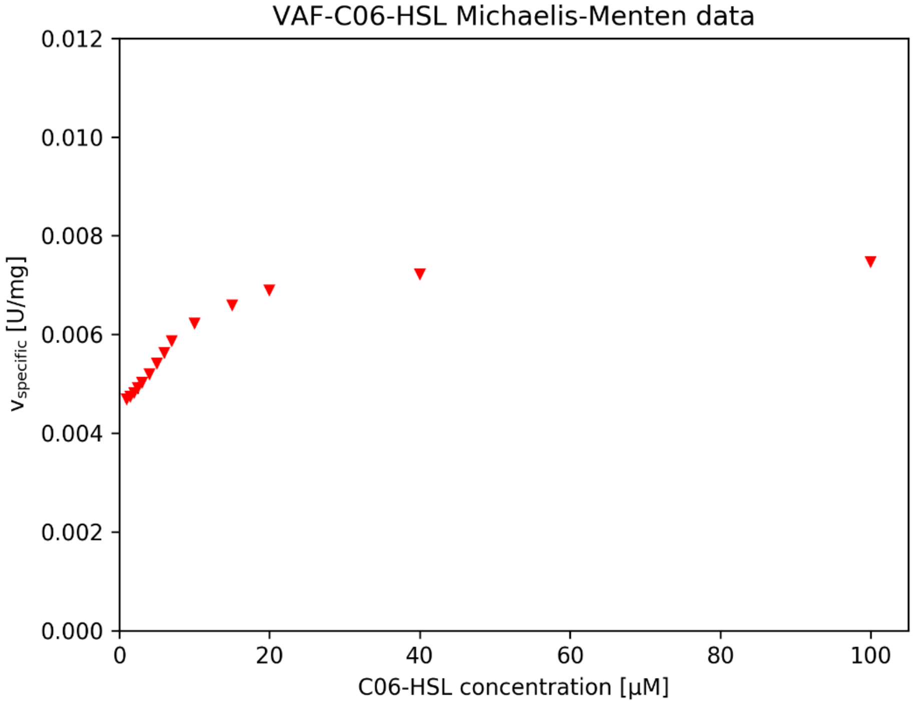

VAF-C06-HSL Michaelis-Menten data
0.012
0.010
0.008
0.006
0.004
0.002
0.000
0
20
40
60
80
100
$v_{specific}$ [U/mg]
C06-HSL concentration [µM]

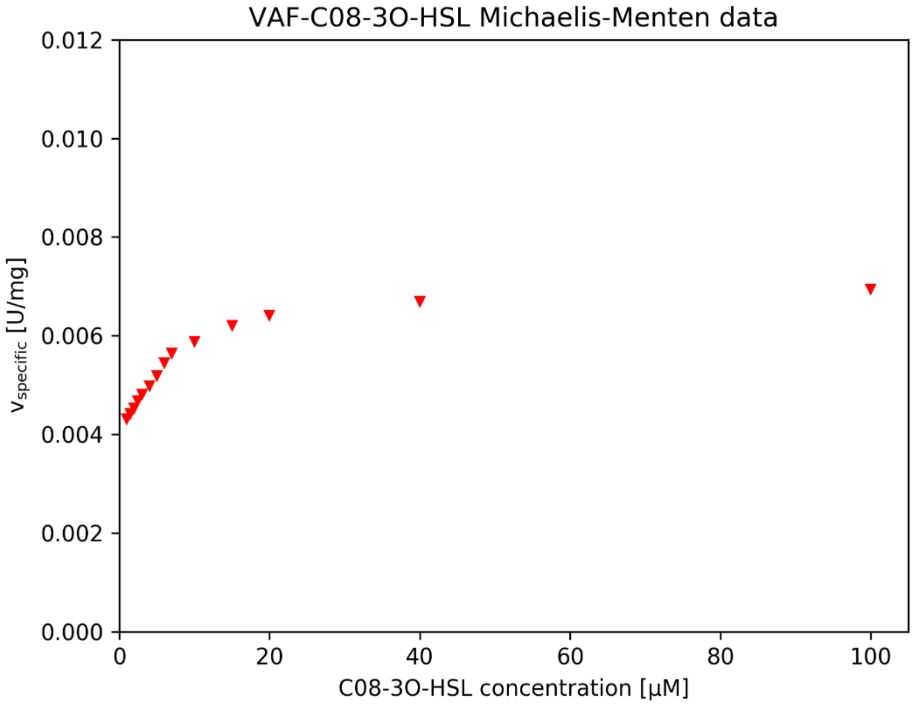
VAF-C08-3O-HSL Michaelis-Menten data
0.012
0.010
0.008
0.006
0.004
0.002
0.000
$v_{specific}$ [U/mg]
0
20
40
60
80
100
C08-3O-HSL concentration [µM]

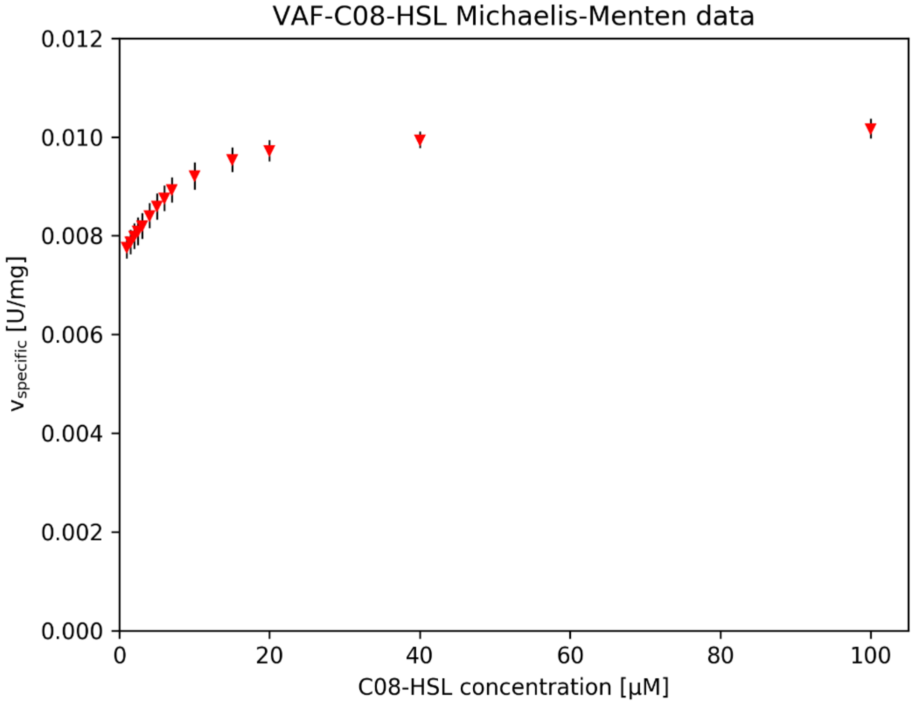
VAF-C08-HSL Michaelis-Menten data
0.012
0.010
0.008
0.006
0.004
0.002
0.000
$v_{specific}$ [U/mg]
0
20
40
60
80
100
C08-HSL concentration [µM]

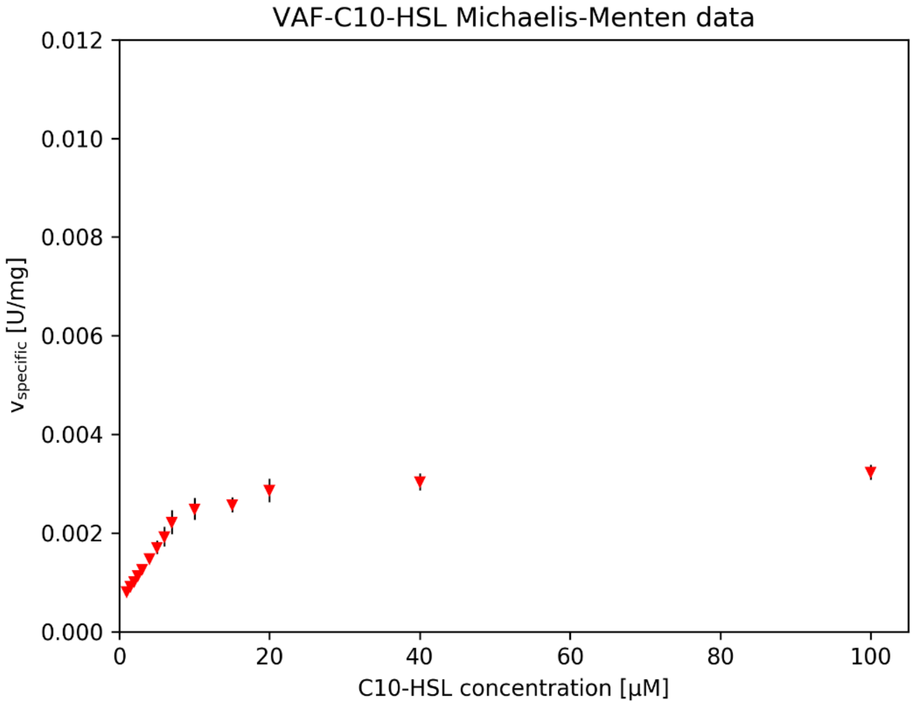
VAF-C10-HSL Michaelis-Menten data
0.012
0.010
0.008
0.006
0.004
0.002
0.000
v specific [U/mg]
0
20
40
60
80
100
C10-HSL concentration [µM]

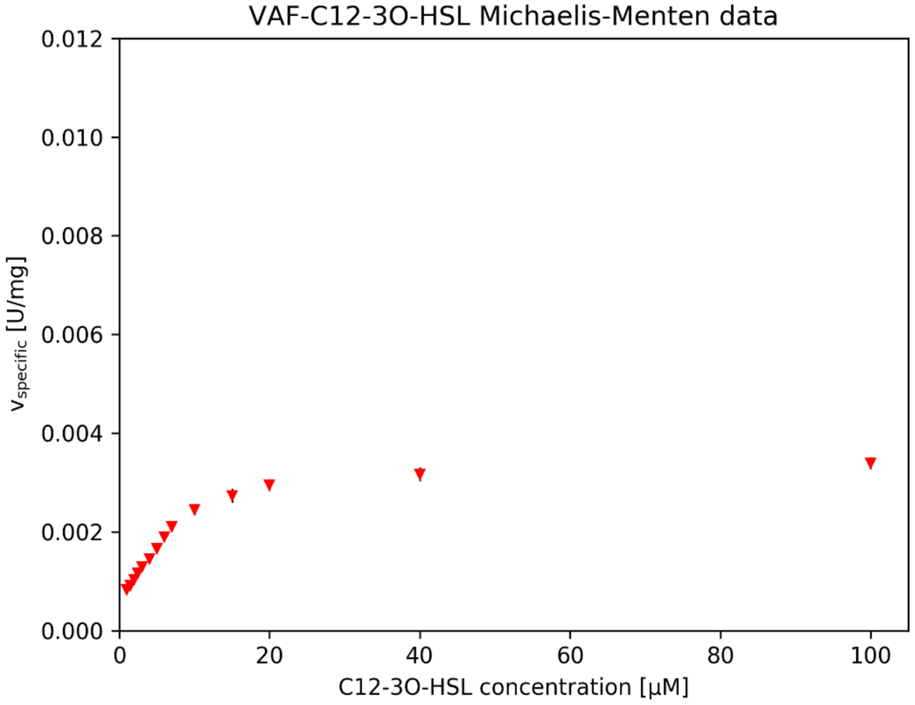
VAF-C12-3O-HSL Michaelis-Menten data
0.012
0.010
0.008
0.006
0.004
0.002
0.000
v specific [U/mg]
0
20
40
60
80
100
C12-3O-HSL concentration [µM]

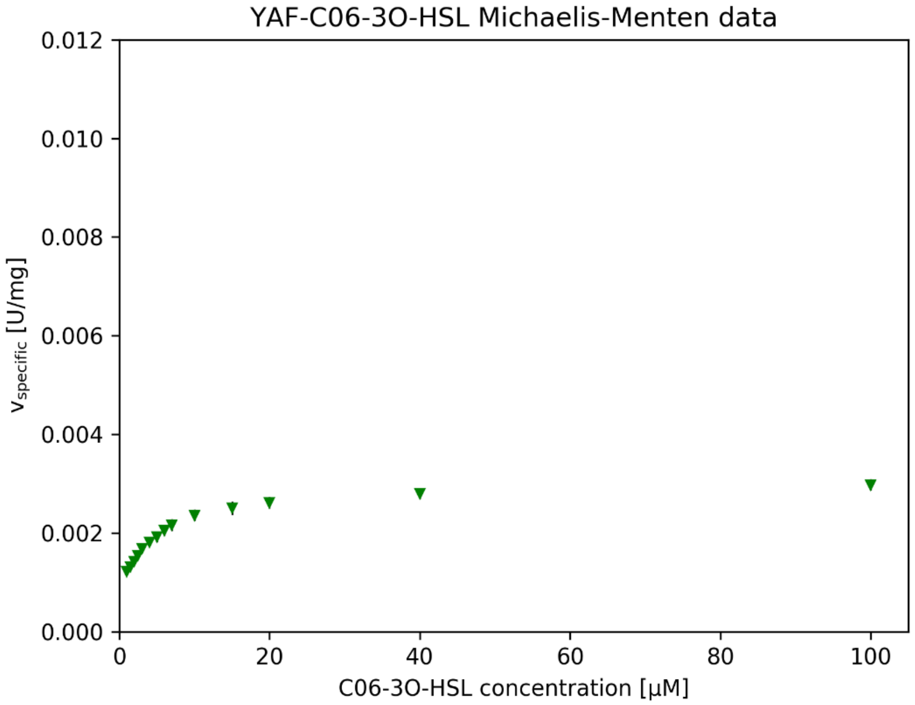
YAF-C06-3O-HSL Michaelis-Menten data
$v_{specific}$ [U/mg]
0.012
0.010
0.008
0.006
0.004
0.002
0.000
0
20
40
60
80
100
C06-3O-HSL concentration [µM]

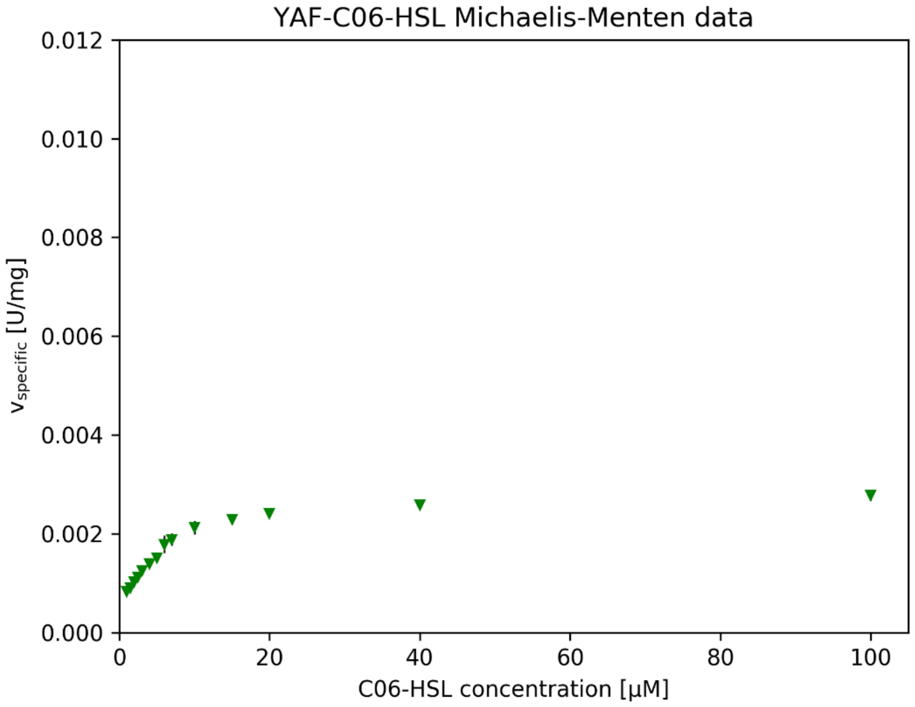
YAF-C06-HSL Michaelis-Menten data
$v_{specific}$ [U/mg]
0.012
0.010
0.008
0.006
0.004
0.002
0.000
0
20
40
60
80
100
C06-HSL concentration [µM]

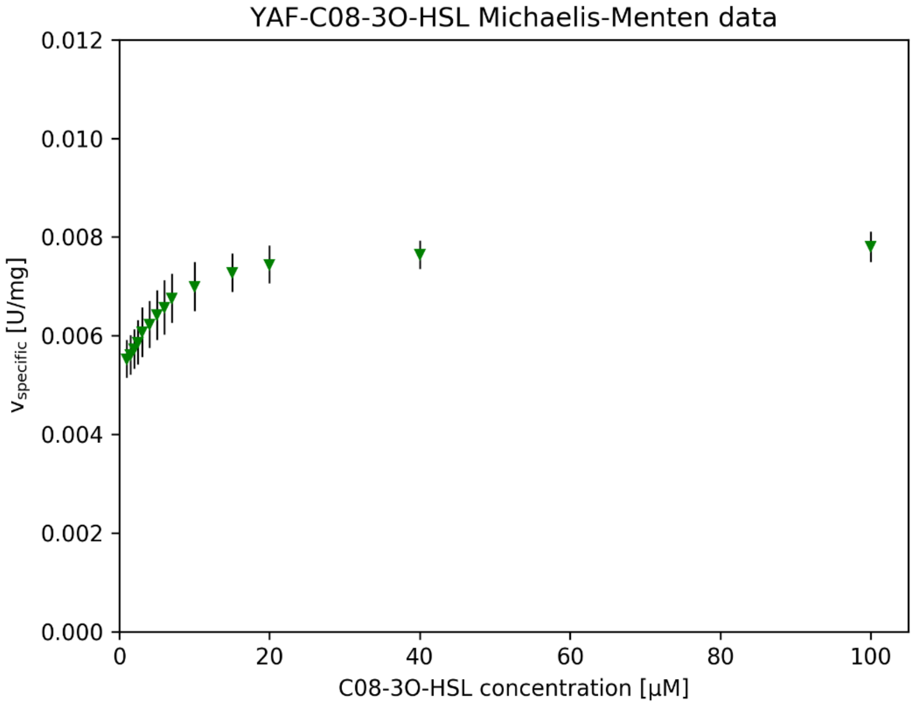
YAF-C08-3O-HSL Michaelis-Menten data
0.012
0.010
0.008
0.006
0.004
0.002
0.000
0
20
40
60
80
100
$v_{specific}$ [U/mg]
C08-3O-HSL concentration [μM]

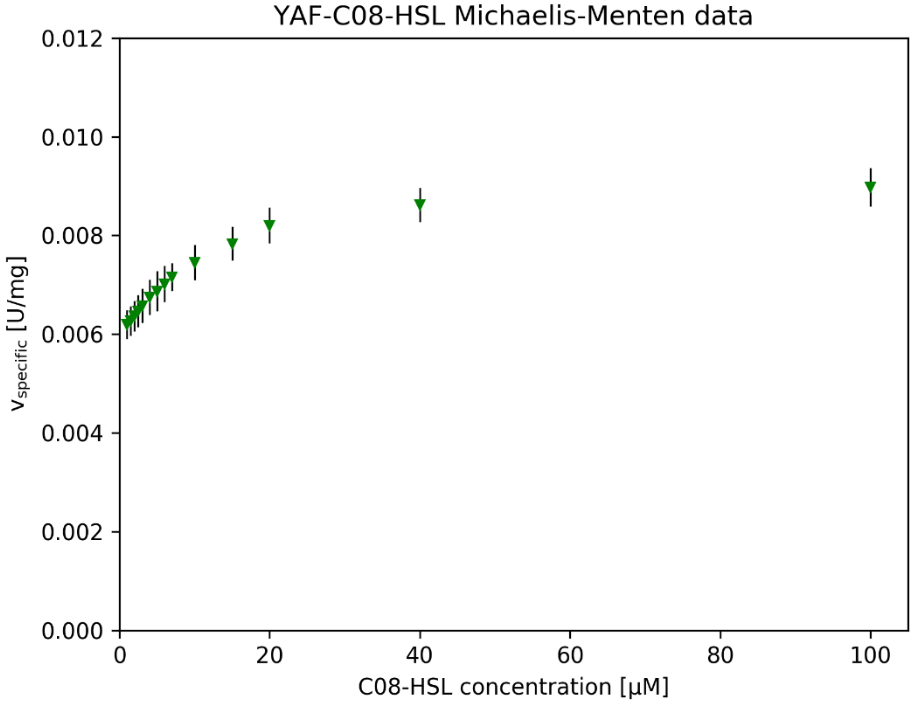
YAF-C08-HSL Michaelis-Menten data
0.012
0.010
0.008
0.006
0.004
0.002
0.000
0
20
40
60
80
100
$v_{specific}$ [U/mg]
C08-HSL concentration [μM]

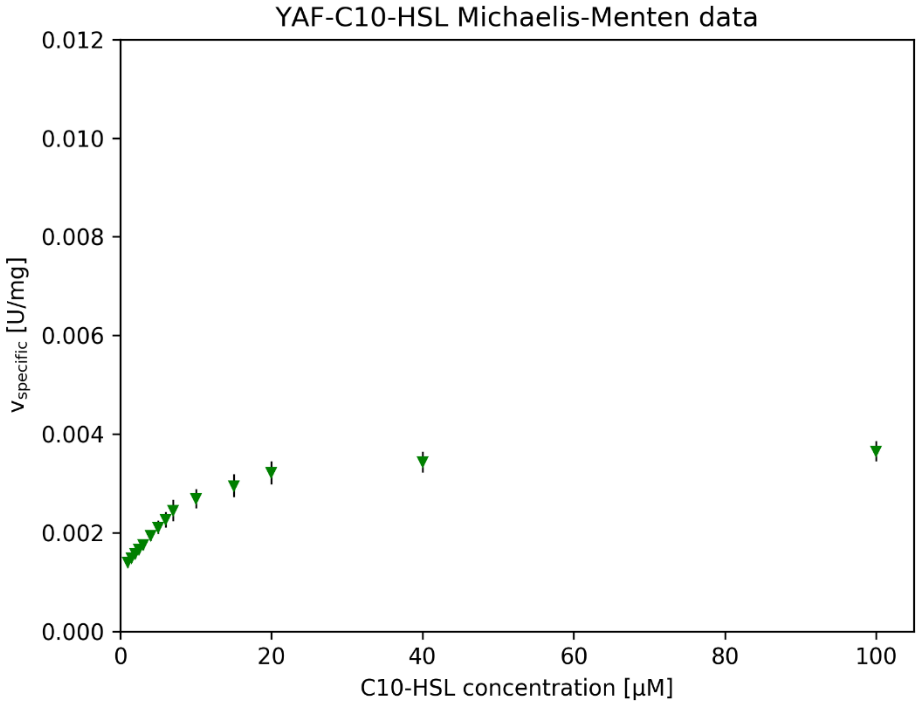
YAF-C10-HSL Michaelis-Menten data
0.012
0.010
0.008
0.006
0.004
0.002
0.000
$v_{specific}$ [U/mg]
0
20
40
60
80
100
C10-HSL concentration [µM]

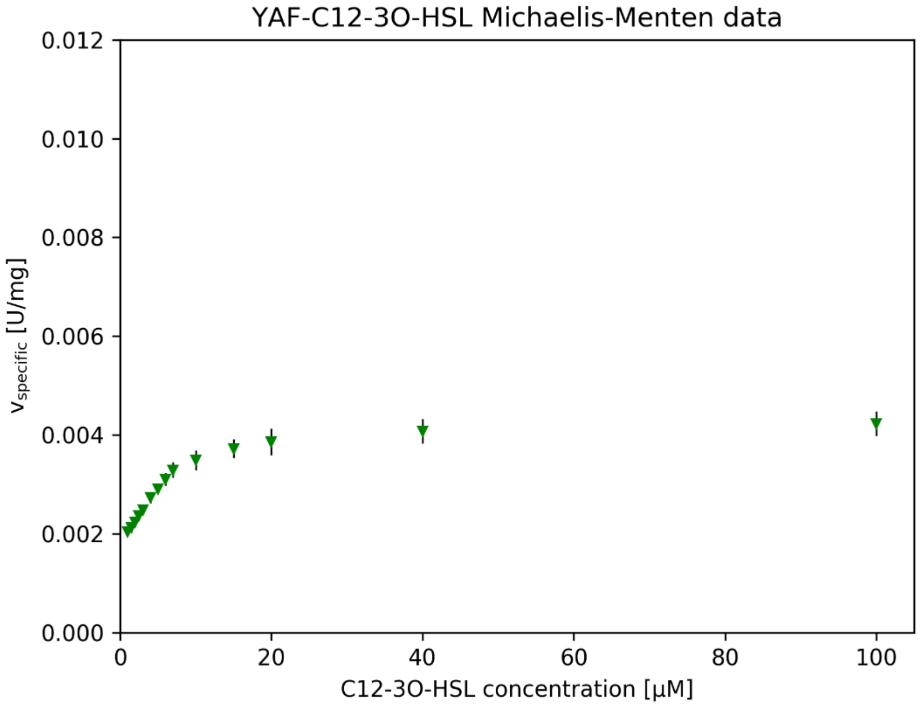
YAF-C12-3O-HSL Michaelis-Menten data
0.012
0.010
0.008
0.006
0.004
0.002
0.000
$v_{specific}$ [U/mg]
0
20
40
60
80
100
C12-3O-HSL concentration [µM]

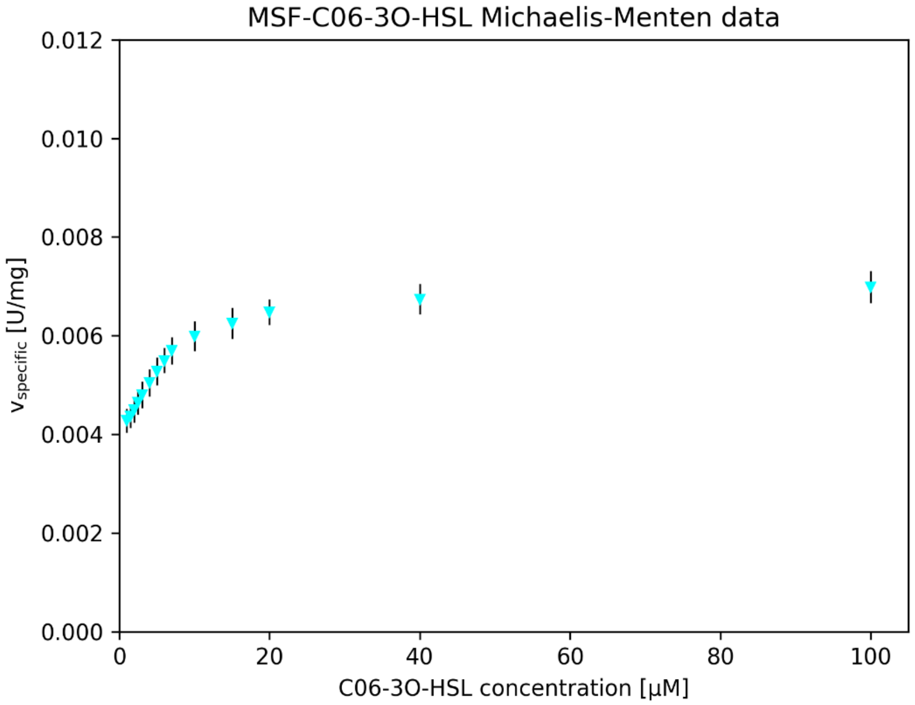
MSF-C06-3O-HSL Michaelis-Menten data
0.012
0.010
0.008
0.006
0.004
0.002
0.000
$v_{specific}$ [U/mg]
0
20
40
60
80
100
C06-3O-HSL concentration [µM]

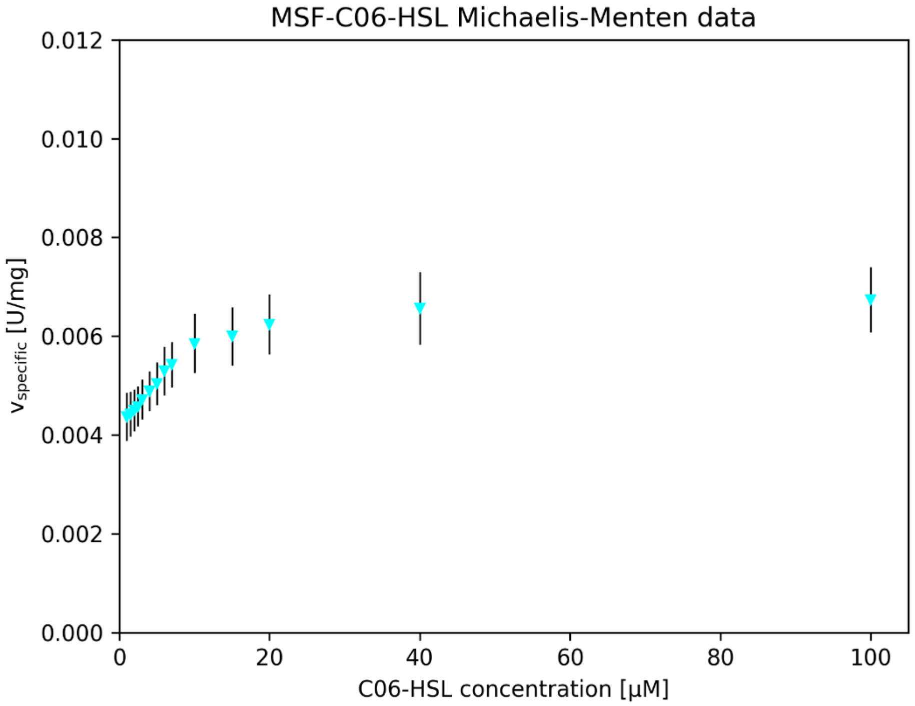
MSF-C06-HSL Michaelis-Menten data
0.012
0.010
0.008
0.006
0.004
0.002
0.000
$v_{specific}$ [U/mg]
0
20
40
60
80
100
C06-HSL concentration [µM]

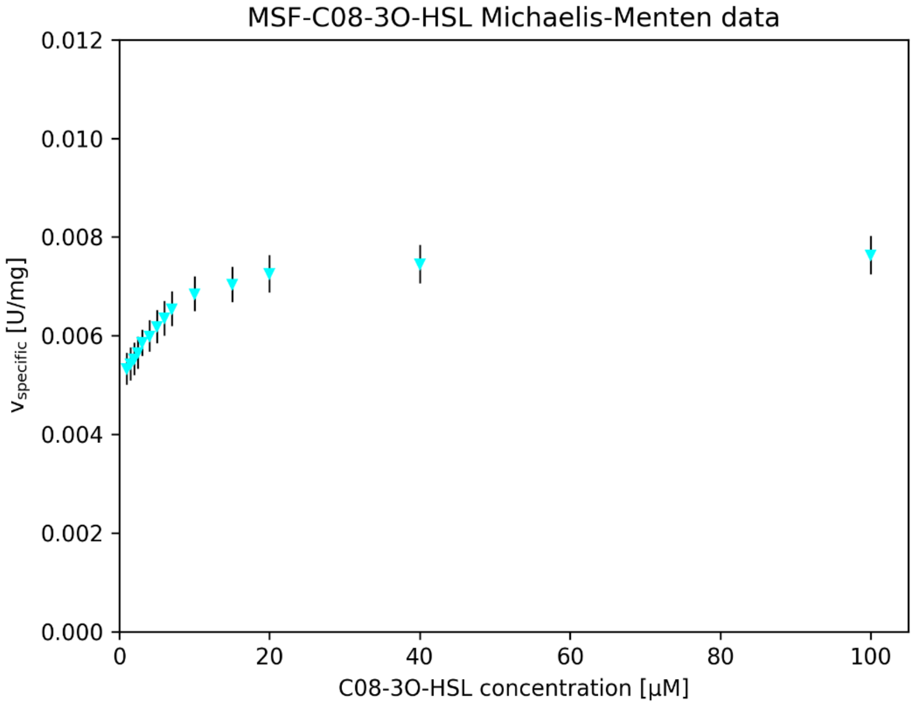
MSF-C08-3O-HSL Michaelis-Menten data
0.012
0.010
0.008
0.006
0.004
0.002
0.000
$v_{specific}$ [U/mg]
0
20
40
60
80
100
C08-3O-HSL concentration [µM]

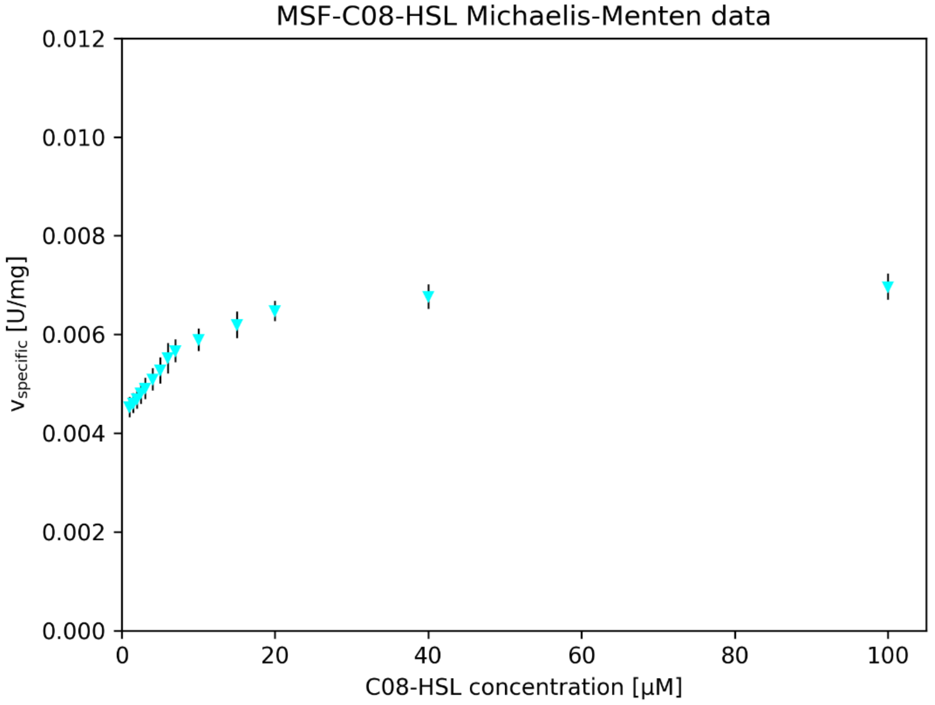
MSF-C08-HSL Michaelis-Menten data
0.012
0.010
0.008
0.006
0.004
0.002
0.000
$v_{specific}$ [U/mg]
0
20
40
60
80
100
C08-HSL concentration [µM]

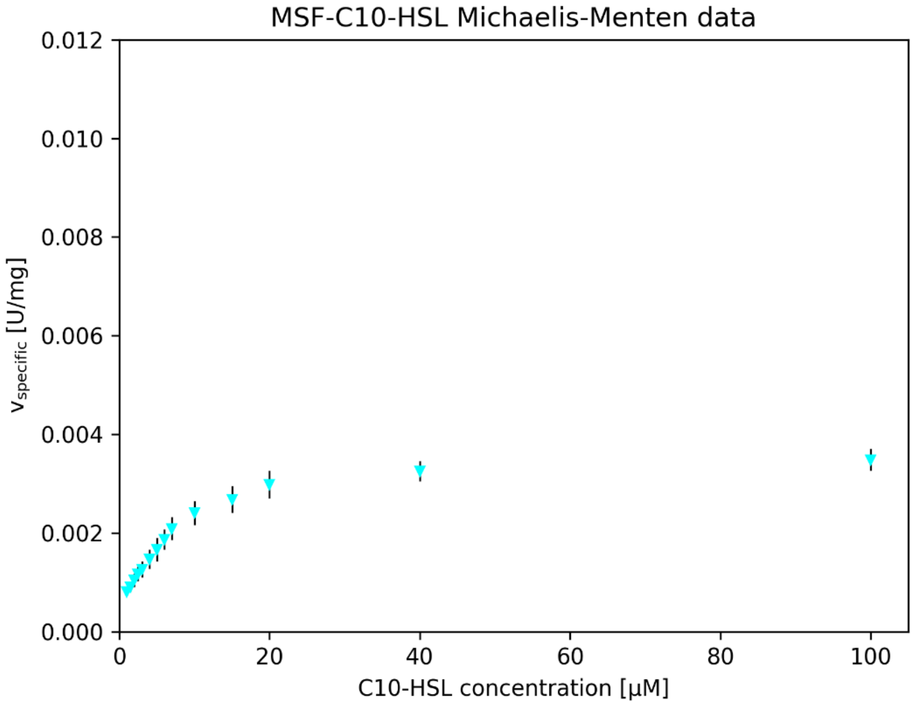

MSF-C10-HSL Michaelis-Menten data
0.012
0.010
0.008
0.006
0.004
0.002
0.000
$v_{specific}$ [U/mg]
0
20
40
60
80
100
C10-HSL concentration [μM]


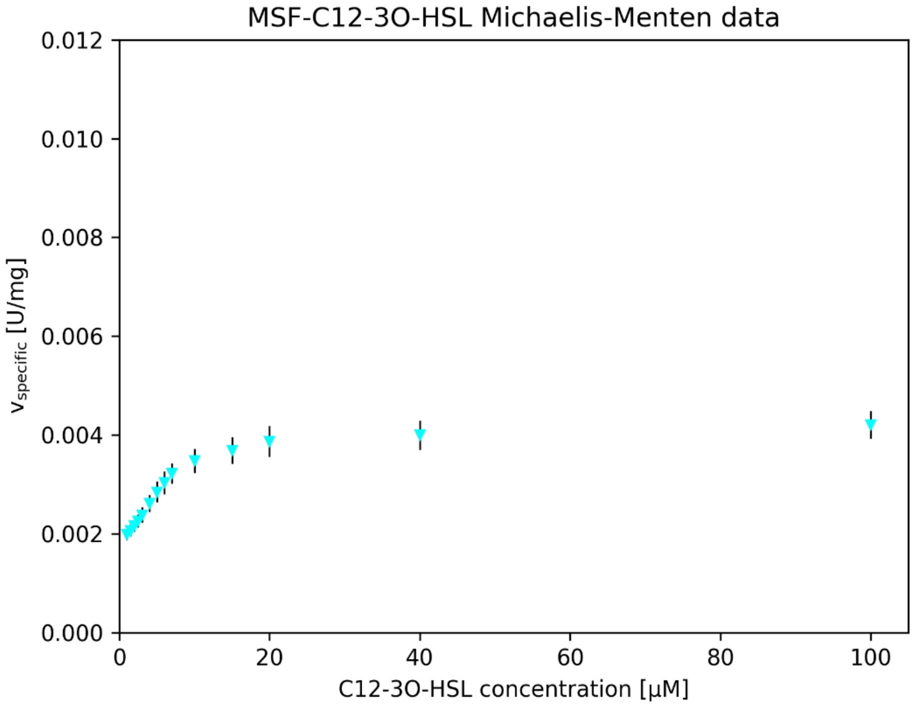

MSF-C12-3O-HSL Michaelis-Menten data
0.012
0.010
0.008
0.006
0.004
0.002
0.000
$v_{specific}$ [U/mg]
0
20
40
60
80
100
C12-3O-HSL concentration [μM]
