## Supplementary Information File 4 for "Engineering dynamic gates in binding pocket of penicillin G acylase to selectively degrade bacterial signaling molecules"

‡ These authors contributed equally.

This supplementary file includes multiplots related to all replicates of molecular dynamics simulations for protein-3-oxo-substrate complexes. On each of particular subplots green curve corresponds to N-acyl-homoserine lactone (AHL) heavy atoms root mean square deviation (RMSD), red continuous line is an AHL average RMSD (left Y-axes). Reactive stabilization defined as properly stabilized AHLs by interactions with oxyanion hole stabilizing residues Ala69β and Asn241β, and exhibiting nucleophile attack distance and angle within 3.3 Å and 75-105°, respectively, is shown as orange line on orange area of the plot (right Y-axes). Hydrogen bond of 3-oxo group of AHL with oxyanion hole stabilizing Ala69β-NH is show as blue line on blue area (right Y-axes).

Full names of presented protein variants:

LAF, Phe138αLeu & Met142αAla & Ile177βPhe ecPGA variant;

MAF, Phe138αMet & Met142αAla & Ile177βPhe ecPGA variant;

VAF, Phe138αVal & Met142αAla & Ile177βPhe ecPGA variant;

YAF, Phe138αTyr & Met142αAla & Ile177βPhe ecPGA variant;

LSF, Phe138αLeu & Met142αSer & Ile177βPhe ecPGA variant;

MSF, Phe138αMet & Met142αSer & Ile177βPhe ecPGA variant

Full names of presented AHLs:

C06-3O-HSL, N-3-oxo-hexanoyl-L-homoserine lactone;

C08-3O-HSL, N-3-oxo-octanoyl-L-homoserine lactone;

C12-3O-HSL, N-3-oxo-dodecanoyl-L-homoserine lactone

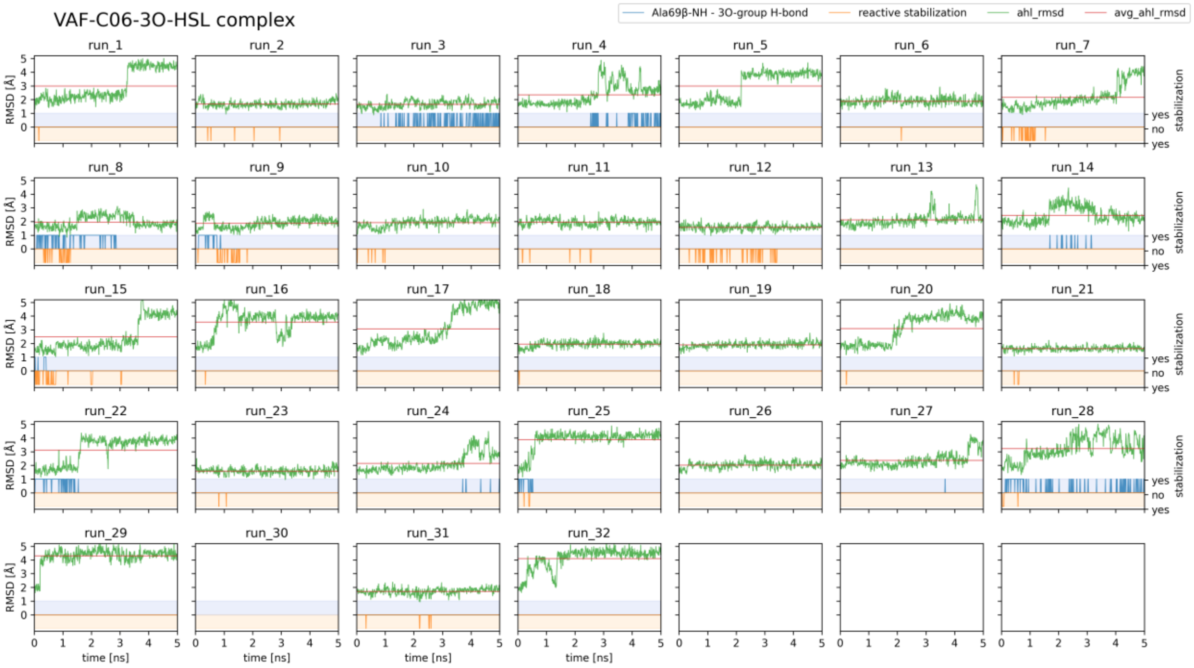
VAF-C06-3O-HSL complex
Ala69β-NH - 3O-group H-bond
reactive stabilization
ahl_rmsd
avg_ahl_rmsd
RMSD [Å]
stabilization
yes
no
time [ns]

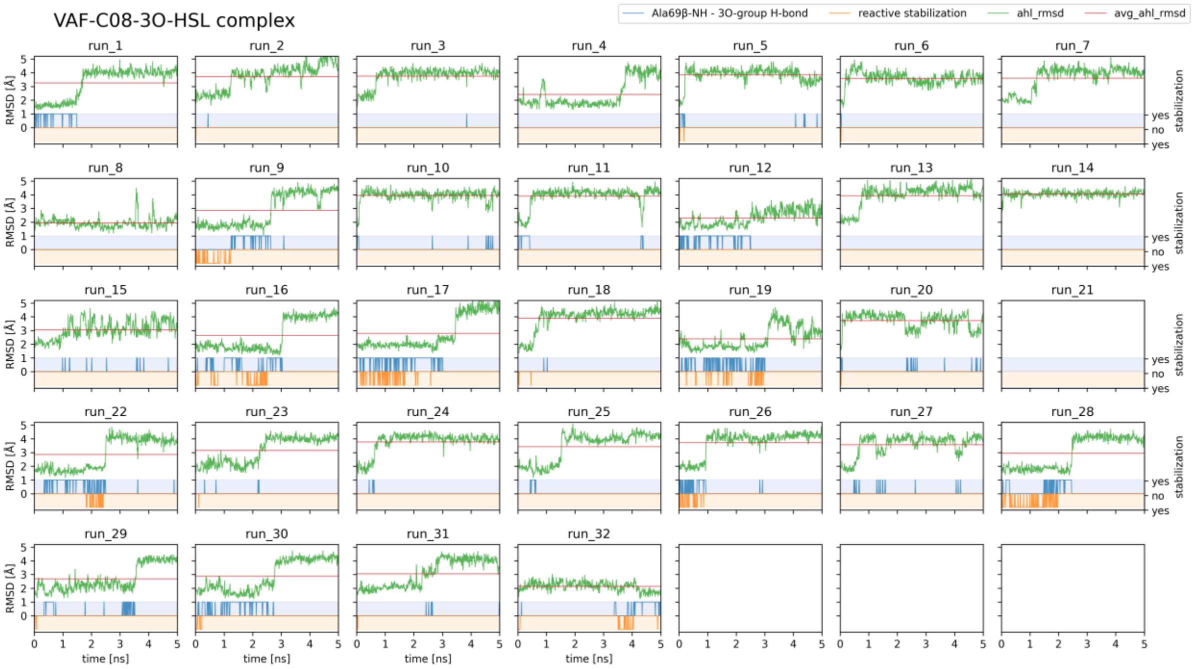
VAF-C08-3O-HSL complex
Ala69β-NH - 3O-group H-bond
reactive stabilization
ahl_rmsd
avg_ahl_rmsd
RMSD [Å]
stabilization
yes
no
time [ns]

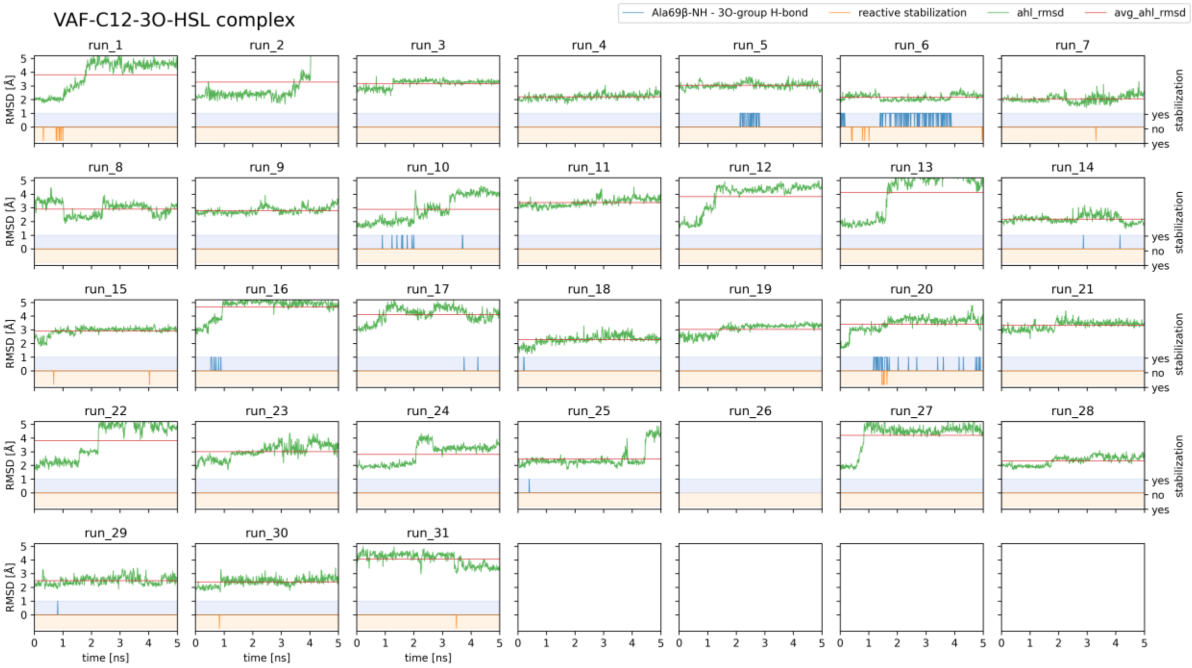
VAF-C12-3O-HSL complex
Ala69β-NH - 3O-group H-bond
reactive stabilization
ahl_rmsd
avg_ahl_rmsd
RMSD [Å]
stabilization
time [ns]

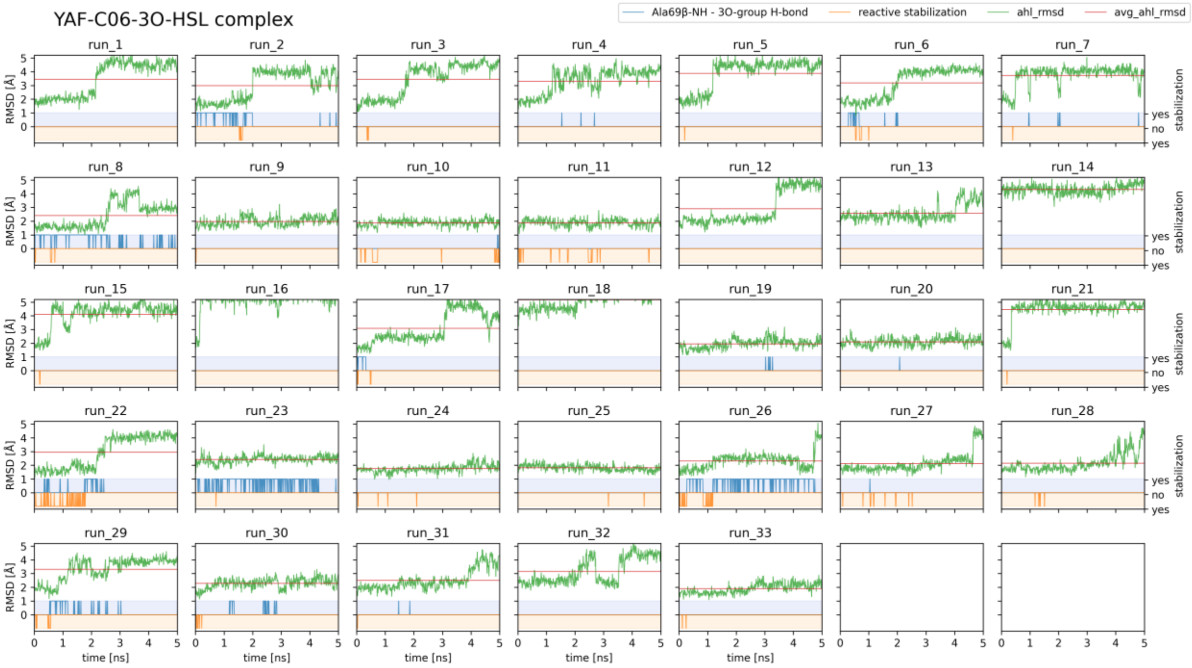
YAF-C06-3O-HSL complex
Ala69β-NH - 3O-group H-bond
reactive stabilization
ahl_rmsd
avg_ahl_rmsd
RMSD [Å]
stabilization
time [ns]

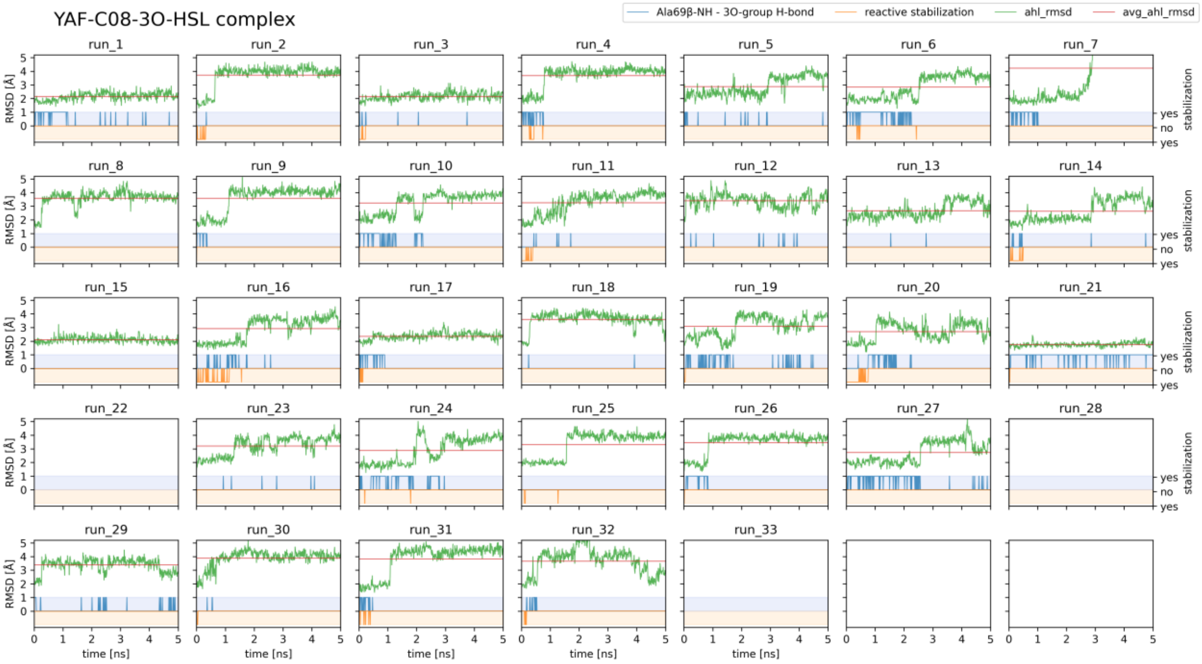
YAF-C08-3O-HSL complex
Ala69β-NH - 3O-group H-bond
reactive stabilization
ahl_rmsd
avg_ahl_rmsd
RMSD [Å]
stabilization
yes
no
time [ns]

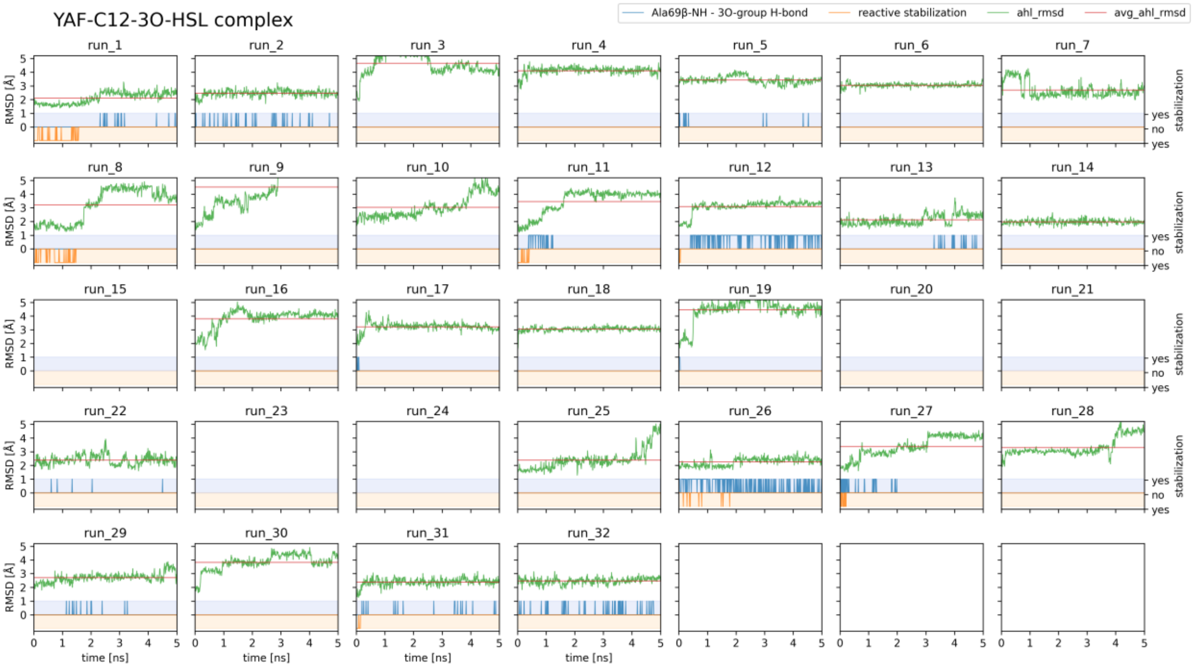
YAF-C12-3O-HSL complex
Ala69β-NH - 3O-group H-bond
reactive stabilization
ahl_rmsd
avg_ahl_rmsd
RMSD [Å]
stabilization
yes
no
time [ns]

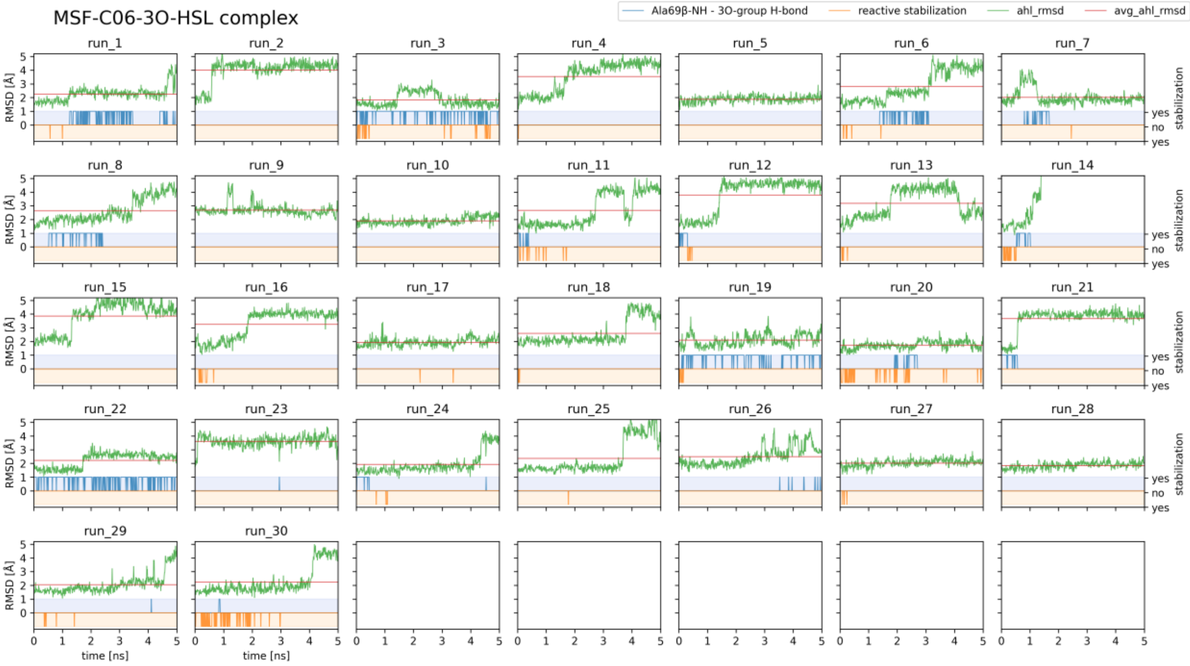
MSF-C06-3O-HSL complex
Ala69β-NH - 3O-group H-bond
reactive stabilization
ahl_rmsd
avg_ahl_rmsd
run_1 … run_30
RMSD [Å]
time [ns]
yes
no
stabilization

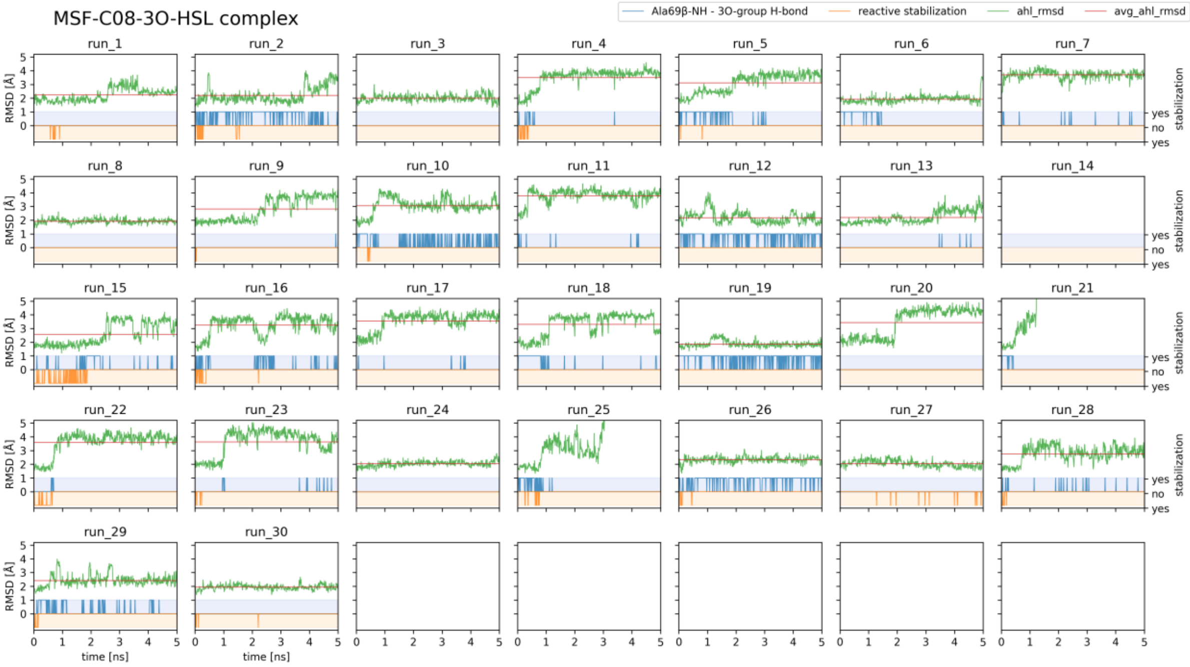
MSF-C08-3O-HSL complex
Ala69β-NH - 3O-group H-bond
reactive stabilization
ahl_rmsd
avg_ahl_rmsd
run_1 … run_30
RMSD [Å]
time [ns]
yes
no
stabilization

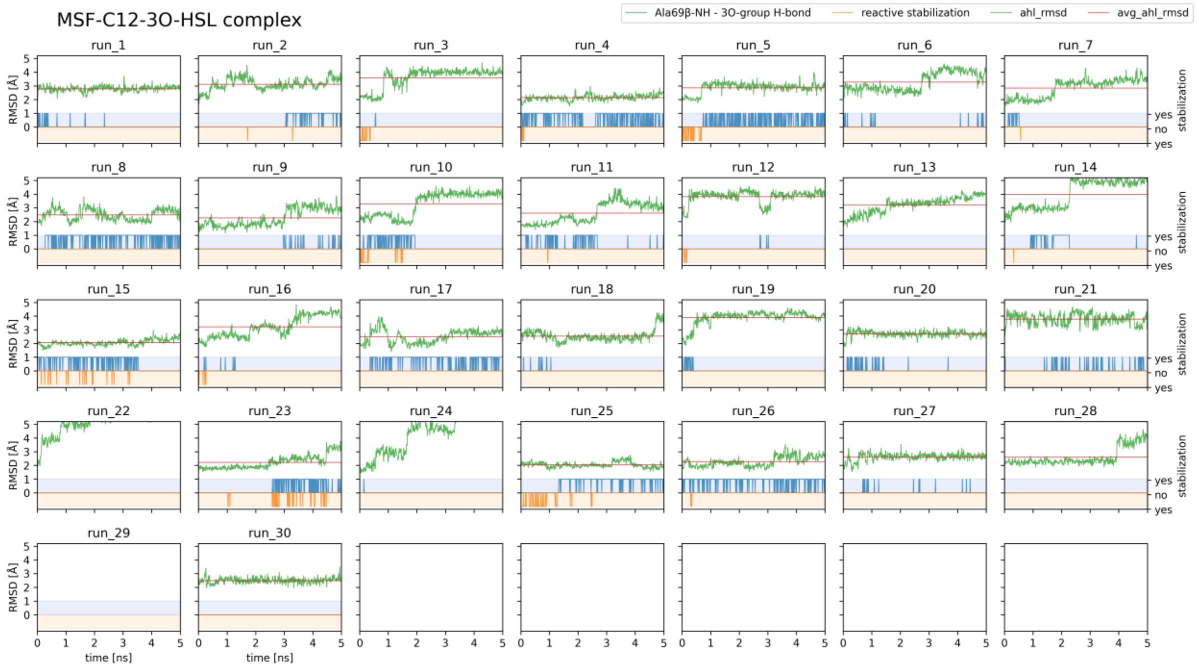
MSF-C12-3O-HSL complex
Ala69β-NH - 3O-group H-bond
reactive stabilization
ahl_rmsd
avg_ahl_rmsd
RMSD [Å]
time [ns]
stabilization
yes
no
yes

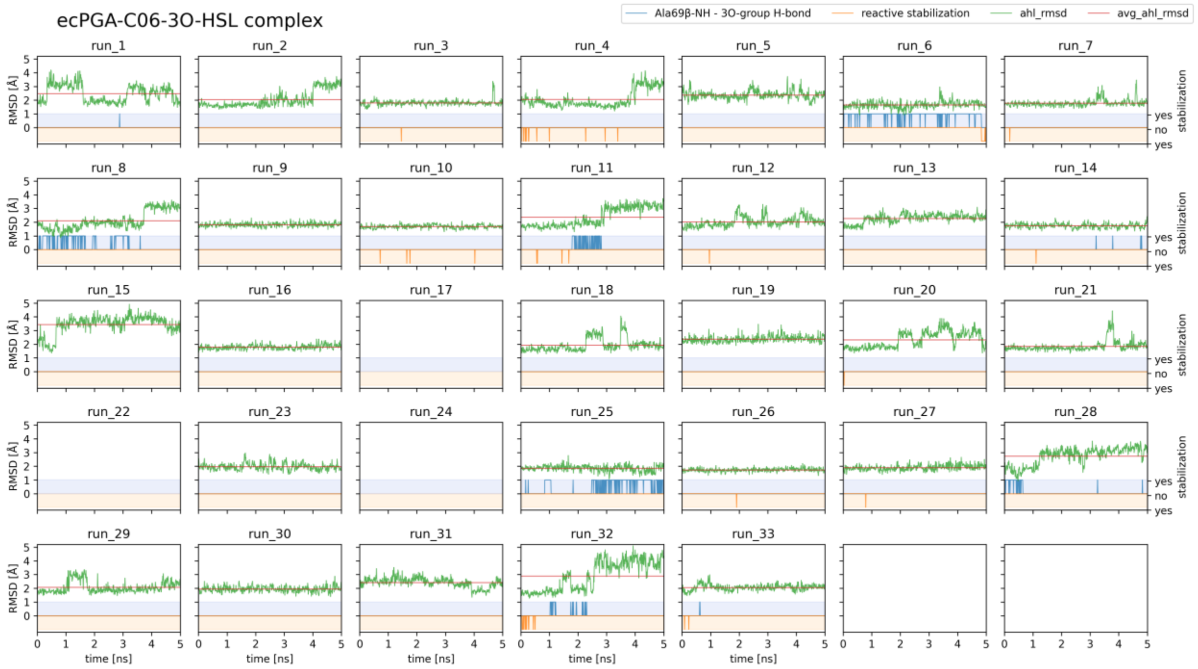
ecPGA-C06-3O-HSL complex
Ala69β-NH - 3O-group H-bond
reactive stabilization
ahl_rmsd
avg_ahl_rmsd
RMSD [Å]
time [ns]
stabilization
yes
no
yes

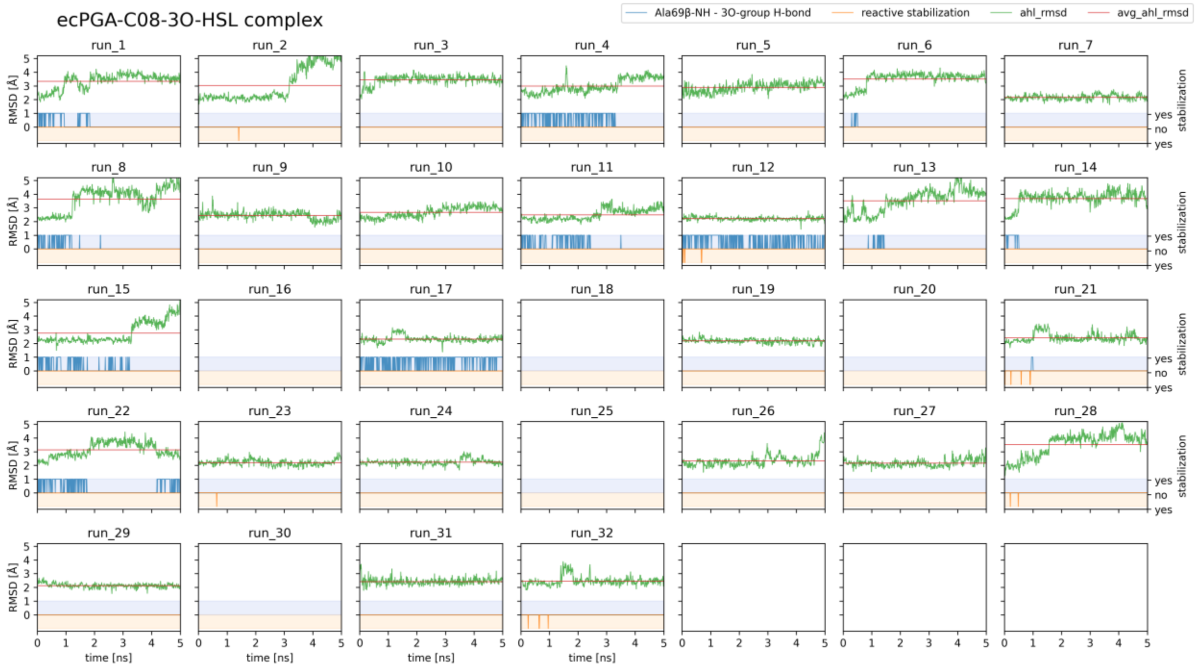
ecPGA-C08-3O-HSL complex
Ala69β-NH - 3O-group H-bond
reactive stabilization
ahl_rmsd
avg_ahl_rmsd
run_1 run_2 run_3 run_4 run_5 run_6 run_7 run_8 run_9 run_10 run_11 run_12 run_13 run_14 run_15 run_16 run_17 run_18 run_19 run_20 run_21 run_22 run_23 run_24 run_25 run_26 run_27 run_28 run_29 run_30 run_31 run_32
RMSD [Å]
stabilization
yes
no
yes
time [ns]

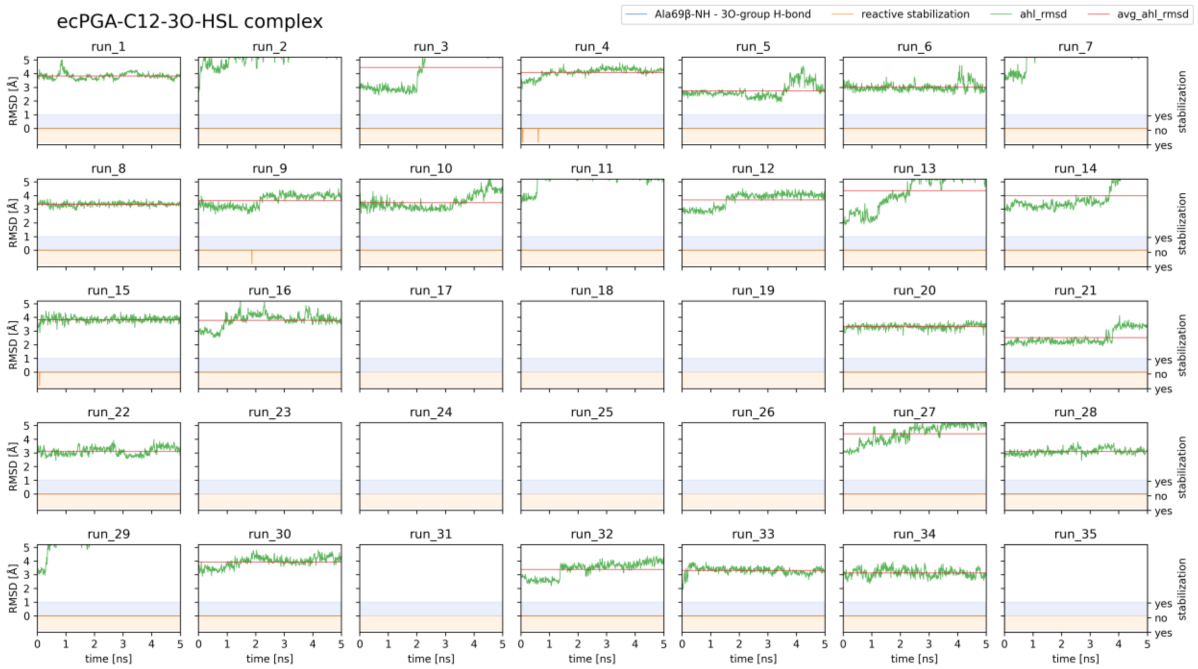
ecPGA-C12-3O-HSL complex
Ala69β-NH - 3O-group H-bond
reactive stabilization
ahl_rmsd
avg_ahl_rmsd
run_1 run_2 run_3 run_4 run_5 run_6 run_7 run_8 run_9 run_10 run_11 run_12 run_13 run_14 run_15 run_16 run_17 run_18 run_19 run_20 run_21 run_22 run_23 run_24 run_25 run_26 run_27 run_28 run_29 run_30 run_31 run_32 run_33 run_34 run_35
RMSD [Å]
stabilization
yes
no
yes
time [ns]
